# The genomics of local adaptation in trees: Are we out of the woods yet?

**DOI:** 10.1101/203307

**Authors:** Brandon M. Lind, Mitra Menon, Constance E. Bolte, Trevor M. Faske, Andrew J. Eckert

**Affiliations:** Integrative Life Sciences, Virginia Commonwealth University, Richmond, Virginia 23284 USA; Department of Biology, Virginia Commonwealth University, Richmond, Virginia 23284 USA

**Keywords:** trees, GWAS, genetic architecture, polygenic local adaptation

## Abstract

There is substantial interest in uncovering the genetic basis of the traits underlying adaptive responses in tree species, as this information will ultimately aid conservation and industrial endeavors across populations, generations, and environments. Fundamentally, the characterization of such genetic bases is within the context of a genetic architecture, which describes the mutlidimensional relationship between genotype and phenotype through the identification of causative variants, their relative location within a genome, expression, pleiotropic effect, environmental influence, and degree of dominance, epistasis, and additivity. Here, we review theory related to polygenic local adaptation and contextualize these expectations with methods often used to uncover the genetic basis of traits important to tree conservation and industry. A broad literature survey suggests that most tree traits generally exhibit considerable heritability, that underlying quantitative genetic variation (*Q*_ST_) is structured more so across populations than neutral expectations (*F*_ST_) in 69% of comparisons across the literature, and that single-locus associations often exhibit small estimated per-locus effects. Together, these results suggest differential selection across populations often acts on tree phenotypes underlain by polygenic architectures consisting of numerous small to moderate effect loci. Using this synthesis, we highlight the limits of using solely single-locus approaches to describe underlying genetic architectures and close by addressing hurdles and promising alternatives towards such goals, remark upon the current state of tree genomics, and identify future directions for this field. Importantly, we argue, the success of future endeavors should not be predicated on the shortcomings of past studies and will instead be dependent upon the application of theory to empiricism, standardized reporting, centralized open-access databases, and continual input and review of the community’s research.

## Introduction

Trees are plants with an arborescent habit, which is loosely defined as a tall-statured growth form usually producing wood (reviewed by Petit & Hampe 2006). Approximately 15% to 25% of plant taxa are classified as trees (Oldfield et al. 1998; Grandtner 2005; Wortley & Scotland 2004), with forested ecosystems accounting for approximately 30% of terrestrial vegetation (Costanza et al. 1997) and providing habitat for terrestrial biodiversity. Indeed, trees play important ecological roles in diverse communities across the globe, such as vertical structural habitat, seeds for wildlife forage, forest cover, the production of oxygen, carbon sequestration, air and water filtration, as well as the reduction of erosion, protracting snowmelt, and desertification. Of these, biological roles are ultimately defined by a set of life history characteristics common to most tree species (Petit & Hampe 2006). These include predominantly outcrossing mating systems with high levels of gene flow and fecundities, as well as long lifespans and generation times (Loehle 1988; Mitton & Williams 2006; Savolainen et al. 2007), although these may differ in, for example, clades of tropical trees. As a result, tree species typically have large effective population sizes, moderate to high levels of genetic diversity, and frequent occurrences of locally adapted ecotypes (Savolainen et al. 2007; Alberto et al. 2013; Sork et al. 2013; Boshier et al. 2015; Prunier et al. 2015; Holliday et al. 2017). Across species, however, rates of morphological and molecular evolution tend to be slow (reviewed in De La Torre et al. 2017). Additionally, genome size varies enormously across species of trees, ranging from 0.4Gbp to 31Gbp (reviewed in Neale et al. 2017). Recent sequencing efforts in gymnosperms, which represent the largest tree genomes, reveal that much of genome size variation is due to transposable element dynamics and gene family evolution (Leitch & Leitch 2012; Morse et al. 2009; Nystedt et al. 2013; Prunier et al. 2015; Neale et al. 2017) where duplication events of select gene families may contribute to the ability of trees to colonize marginalized habitats (Leitch & Leitch 2012; Prunier et al. 2015; Neale et al. 2017).

In trees, the general presence of large geographical ranges and extensive gene flow also provides an ideal setting to disentangle neutral from selective evolutionary processes (Neale & Kremer 2011). Indeed, their longevity and wide and heterogeneous geographical distributions lend trees suitable for addressing several key evolutionary questions about the importance of historical climatic fluctuations, and local adaptation involving shifts in allele frequencies (Lotterhos & Whitlock 2014; Savolainen et al. 2007, 2013; Platt et al. 2015). As we detail in subsequent sections, evidence consistent with local adaptation in trees is ubiquitous, even across fine spatial scales where it had been hypothesized that gene flow may overcome selection of locally favored alleles (e.g., Mitton et al. 1998; Budde et al. 2014; Csilléry et al. 2014; Vizcaíno-Palomar et al. 2014; Eckert et al. 2015; Holliday et al. 2016; Roschanksi et al. 2016; Lind et al. 2017).

Quantitative phenotypes are often used as a proxy for total lifetime fitness, which is composed of two broad components: survival and reproduction. Since most quantitative traits are related to some component of total lifetime fitness, they are often used to assess potential for local adaptation. For many plant taxa, selection pressures are expected to be strongest for variation in survival during the juvenile stages of development (Donohue et al. 2010), particularly for those taxa with high reproductive output, as is the case for many tree species. As such, juvenile stages in plants have been found to contribute substantially to total lifetime fitness (Postma & Agren 2016). Phenotypic traits associated with juvenile survival have thus received the majority of genetic research focus in trees, particularly due to their long-lived nature. Such studies have led to intriguing insights gained through a long history of common garden experimentation (Langlet 1971; Morgenstern 1996). For example, traits such as growth (e.g., height and diameter), form (e.g., specific gravity, straightness), phenology (e.g., bud flush, bud set), juvenile performance (e.g., germination rate, seed traits) and physiology (e.g., cold hardiness, water-use efficiency) have all been shown to be under moderate to high genetic control (reviewed in Corn-elius 1994, Howe et al. 2003, Alberto et al. 2013; this review). Variation for these traits is also often partitioned among populations (this review), despite the vast majority of neutral variation remaining within populations (Howe et al. 2003; Neale & Savolainen 2004). With few exceptions (e.g., major gene resistance in the white pine-blister rust pathosystem; Kinloch et al. 1970; Liu et al. 2017), variation for these traits forms a continuum across individuals, thus implying that the underlying genetic architecture is composed of a large number of small to moderate effect loci (i.e., a polygenic architecture; concept reviewed in Savolainen et al. 2007, 2013; Gagnaire & Gaggiotti 2016; Hoban et al. 2016; Timpson et al. 2017). There is some uncertainty, however, concerning the properties of the effect size distributions comprising polygenic architectures *(sensu* Fisher 1930, Kimura 1983, and Orr 1998), the relative importance of various forms of gene actions (e.g., dominance, epistasis) in producing trait variation (Crow 2010, Hansen 2013), how these interact to affect the evolution of polygenic architectures in natural populations (Hansen 2006), and how these factors will ultimately influence evolutionary processes and outcomes in forest trees (Savolainen et al. 2007; Sork et al. 2013; Prunier et al. 2015). Considerable strides, made in the past through genotype-phenotype-environment studies (*sensu* Sork et al. 2013), have contributed intriguing insight into the genomic basis of local adaptation for tree species. However, given the large genome size of many tree species, such methods have been criticized as lacking in power and sufficient coverage needed to detect small effect loci, which is further exacerbated by rapid decay of linkage disequilibrium (LD) in most forest trees (Mackay 2009; Savolainen et al. 2007). Despite these limitations, association studies have been moderately successful in linking genotypes and phenotypes, including providing information for making inferences about local adaptation.

In this review, we set out to summarize theory related to polygenic local adaptation and, using these expectations, contextualize the progress of describing the genetic architectures underlying traits important to conservation and industry in undomesticated tree species. We first highlight the extensive evidence for local adaptation in trees by reviewing transplant designs often used in investigations of quantitative genetic differentiation. Using an extensive literature survey across both gymnosperm and angiosperm species, we provide an overview of these transplant methods, give examples of each, and quantify the distribution of narrow sense heritability and *Q*_ST_ estimates across various trait categories. We further use this survey to establish patterns of comparative quantitative and neutral genetic differentiation (i.e., *Q*_ST_-*F*_ST_ tests) which until this review had not been suitably synthesized in trees. Before we transition into discussing common methods used to uncover loci underlying adaptation, we establish expectations for the genetic architecture of polygenic, fitness-related traits by reviewing the theory available to date. We then provide an extensive review of genotype-phenotype associations in trees and provide the distribution of the percent phenotypic variance explained by empirically associated loci. Using this distribution, we underscore the limitations of using solely single-locus approaches to uncover the loci underlying local adaptation in tree species. Given this synthesis, we highlight exemplary genomic resources available to fill knowledge gaps, identify promising avenues of future research, identify key benchmarks and necessary steps towards truly integrating studies of trees into the genomic era, and address our primary question, “Are we out of the woods yet?”.

## Identifying heritable phenotypic variation

Trees have evolved numerous adaptations as a result of their vast ecological breadth. As such, it has long been the goal of forest scientists to understand the traits important to viability and persistence. Among the most frequent designs used, common gardens and reciprocal transplants have aimed at describing genetically based differentiation of measured phenotypes among various source populations of varying sizes and across various geographic scales. Across these designs, investigators seek to better understand the phenotypes relevant to local adaptation and the selective pressures influencing these phenotypes. The exact design chosen, however, is generally based on the questions driving the research endeavor and often by the availability of resources (Morgenstern 1996; Blanquart et al. 2013; de Villemereuil et al. 2015). In this section, we briefly review these designs, identify relevant questions and inferences, highlight some of the important practical applications of these techniques, and discuss examples of past investigations in various tree species.

There is a rich history of forest scientists using the common garden approach dating back hundreds of years (Langlet 1971; Mátyás 1996). In a broad sense, a common garden design is used to test for differentiation among genetically distinct groups in a homogeneous environment. These groups can be clonal replicates or sibships (families) derived from species or hybrids sampled from various populations, provenances, varieties, cultivars, or agricultural accessions (Cheplick 2015). When individuals from various origins are grown together under the same conditions, the observed phenotypic differentiation is expected to reflect underlying genetic variation, especially when maternal effects are assumed or shown to be absent. Common garden and provenance trial designs can also establish evidence that the phenotypes under study are heritable, a prerequisite for an adaptive response to selective agents (Supplemental Box S1), and that populations exhibit quantitative genetic differentiation (i.e., *Q*_ST_; Spitz 1993). When driven by questions related to differentiation alone, a single common garden approach can be used to describe levels of quantitative genetic variation within and among genetically distinct groups. In these cases, no environmental variables are manipulated, and thus, unequivocal evidence for trait divergence among groups, and the contributing factors influencing this divergence (e.g., neutral or selective processes), is often limited because conclusions must be based on *post hoc* inferences about source environments for the materials established in the common garden. Even so, single common garden approaches can be a powerful tool to demonstrate evidence congruent with local adaptation. For instance, the white carob tree (*Prosopis alba* Griseb., Leguminosae) growing in Argentina is an ideal multipurpose tree that has potential for use in reforestation and afforestation applications in the region. However, this genus is known to invade other regions, encroach on farmland and waterways, and has a thorny growth habit that can cause sepsis in livestock. To better understand how forestry applications can balance the benefits of production and forest protection, Bessega et al. (2015) used a single common garden representing eight provenances of *P. alba* to compare estimates of neutral genetic patterns to the quantitative genetic variation of life history traits related to economic importance. They found that for most traits there existed considerable underlying genetic variation 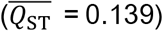. Additionally, source environments were often correlated with measured trait variation in the common garden, suggesting that the observed differentiation was driven by temperature, precipitation, wind speed, and sunshine fraction, with signals of divergent selection corroborated across *Q*_ST_-*F*_ST_ comparisons and tests for selection (e.g., S test, *sensu* Ovaskainen et al. 2011). Bessega et al. (2015) concluded that the signal of non-neutral differentiation was indicative of divergent phenotypic optima across populations, and that this variation could be used to direct future breeding programs across the region.

When there is evidence that environmental differences among source populations may be driving adaptive divergence, strong environmental candidates can be manipulated (artificially or via site selection) in a multiple common garden design to further investigate hypotheses of differentiation and adaptation. For instance, the sweet chestnut (*Castanea sativa* Mill., Fagaceae), also known for its edible fruit, is distributed across much of Minor Asia and southern Europe and is an ecologically important component of many Mediterranean systems. *Castanea sativa* exhibits ecological, physiological, morphological, and genetic variability as the range overlays a climatic transition from xeric Mediterranean conditions to wetter Euro-Siberian environments (see refs in Lauteri et al. 2004). Previous common garden experiments carried out by Lauteri and colleagues have indicated that populations across this transition are further differentiated by water use efficiency (the ratio of plant carbon gain to water loss) and carbon isotope discrimination, Δ. To further explore variability of drought-related traits, Lauteri et al. (2004) used an *ex situ* multiple common garden design using two water and temperature treatments in individual climatic chambers to assess differentiation among six populations across Spain, Italy, and Greece. They found *treatment* and *population x treatment* effects were significant, suggesting variation in drought adaptation across populations. Additionally, populations originating from dry sites generally exhibited higher values of Δ, which was also composed of significant additive genetic variation (*h*^2^= 0.15-0.52), and suggests that genetic and physiological mechanisms of drought adaptation confer a capacity to colonize a wide arrange of environmental conditions, while strong negative relationships between Δ and growth-related traits is suggestive of strong evolutionary constraints at juvenile stages.

While *ex situ* common gardens approaches (e.g., Lauteri et al. 2004) can provide strong evidence of adaptive divergence among populations, and in some cases corroborate putative drivers of observed differentiation, these studies can often exclude key environmental factors, possibly leading to confounding signals of adaptation (Kawecki & Ebert 2004). When *in situ* experimentation is feasible, site selection can be used to test for environmental drivers of local adaptation. For example, Evans et al. (2016) investigated traits related to growth and phenology in juvenile narrowleaf cottonwood (*Populus angustifolia* James, Salicaceae) by planting families from nine populations across the native range into three common gardens, one each at the northern, southern, and interior extent of the range. Using *Q*_ST_-*F*_ST_ comparisons and clinal analyses alongside the quantitative genetic analyses, Evans et al. (2016) concluded that climate cues played a major role in structuring adaptive variation across the range of *P. angustifolia,* and that future industrial and conservation applications should utilize this information to inform source environments for optimal outcomes.

As both *in situ* and *ex situ* common garden trials can include multiple environmental influences in their design, reciprocally transplanting to all source environments is not necessarily a requirement to decompose genetic variation underlying adaptive traits or to provide evidence for, or the drivers of, differentiation among populations. Thus, these designs may preclude inferences regarding local adaptation *sensu stricto.* To produce such evidence, source populations can be planted in a (full- or incomplete-factorial) reciprocal transplant design and allow for traits related to fitness to be assessed across native and non-native environments. If a population is locally adapted, individuals exposed to their native environments should show increased growth, survival, and reproduction relative to non-native genotypes (Kawecki & Ebert 2004; Leimu & Fischer 2008; Hereford 2009; Savolainen et al. 2013). For example, with the goal of delineating conservation units based on molecular and quantitative trait differentiation, Rodríguez-Quilón et al. (2016) used four reciprocally-transplanted common gardens to assess height and survival of samples from 35 natural populations of maritime pine (*Pinus pinaster* Aiton, Pinaceae). For both traits, *Q*_ST_ was consistently larger than *F*_ST_ across the four sites, a pattern suggestive of divergent selection. Six distinct gene pools based on evolutionary history of neutral markers were identified, and because high quantitative differentiation (*Q*_ST_) was found within these pools, hierarchical analyses were used to further identify ten adaptive population groups for use in conservation and breeding approaches.

Available evidence suggests that many populations of tree species have substantial heritable genetic variation, and that the quantitative traits under study often show signals of divergent selection across both broad and fine spatial scales. But how broadly can we apply this statement? Are there overall patterns of heritability and quantitative genetic structure across tree species? Because estimates of heritability and *Q*_ST_ are often only applicable to a specific set of populations, for a specific set of environments, at any specific point in time (e.g., see Figure 2D), a large sample of these estimates is therefore necessary to synthesize the current literature with regard to patterns across taxa. To accomplish this aim, we synthesized estimates from 129 published studies with estimates of narrow sense heritability (n = 114) from replicated progeny trials and/or estimates of quantitative genetic differentiation (*Q*_ST_; n = 37). However, we excluded papers that have been cited for estimates of *Q*_ST_ or heritability that were calculated *post hoc* from variance components (i.e., we only recorded estimates that were explicitly reported as *h*^2^ or *Q*_ST_ in the original publication). For comparison, we further grouped measured traits into 14 broad categories: cold hardiness, disease resistance, drought hardiness, form, growth, herbivore and insect resistance, leaf and needle properties, phenology, plant secondary metabolites, reproduction, resource allocation, seed and early germination properties, survival, and wood properties. Because sample size can influence the precision of both heritability and *Q*_ST_, for each trait category we used a weighted average where weights were equal to the number of families used to estimate variance components for each estimate of *h*^2^ and *Q*_ST_.

In agreement with Cornelius (1994), our survey found that many of the traits important to conservation and industry exhibit non-zero narrow sense heritability (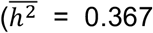; File S1; Figures S1–S4) and are thus amenable to selection. The mean weighted *Q*_ST_ across traits groups from our survey (Table S1; File S1) was between 0.10-0.28, except for drought hardiness (0.06) and disease resistance (0.04), with median values from the unweighted distribution generally falling below the weighted average for each trait group (Figure 1). This suggests that over various geographic and environmental distances, population histories, and species, there is a general pattern of substantial genetic variation underlying measured traits. Given our synthesis of *Q*_ST_ estimates in trees, we were curious of the evidence for adaptive divergence among populations (*Q*_ST_ > *F*_ST_). Of the 37 articles reporting *Q*_ST_ estimates in our review, 23 compared *Q*_ST_ with *F*_ST_ or G_ST_ estimated from the same populations under study (however, we excluded studies that used *F*_ST_ measurements taken from the literature, e.g., as in McKay & Latta 2002; Alberto et al. 2013). Indeed, as pointed out by Crnokrak & Merilä (2002), comparisons of *Q*_ST_ and estimated from different populations and/or at different time points are uninformative. Of these 23 studies, 18 compared *Q*_ST_ and *F*_ST_ in a statistical framework while the remaining five studies compared *Q*_ST_ and *F*_ST_ numerically. Across numerical and statistical comparisons combined, 67% (254 of 381 traits) exhibited higher *Q*_ST_ than *F*_ST_, with 69% (170 of 246 traits) exhibiting significantly higher *Q*_ST_ than *F*_ST_. Although we did not tally instances where *Q*_ST_ was reported to be less than *F*_ST_ (statistically or otherwise), as this was not the focus of our review, there were some instances in which this was the case. For instance, Lamy et al. (2011) found such patterns when quantifying population genetic differentiation of cavitation resistance across the species range of maritime pine (*Pinus pinaster* Aiton, Pinaceae), while Mahalovich et al. (2011) also found that *Q*_ST_ < *F*_ST_ for traits related to white pine-blister rust resistance in inoculated seedlings of whitebark pine (*Pinus albicaulis* Engelm., Pinaceae). While various explanations for such patterns were outlined by Lamy et al. (2011), canalization was argued as the most likely process driving the observed patterns, while Mahahlovich et al. (2011) offered similar arguments for selection favoring the same genotype in different environments (see Lamy et al. 2012 for more regarding this aspect).

**Figure 1.**
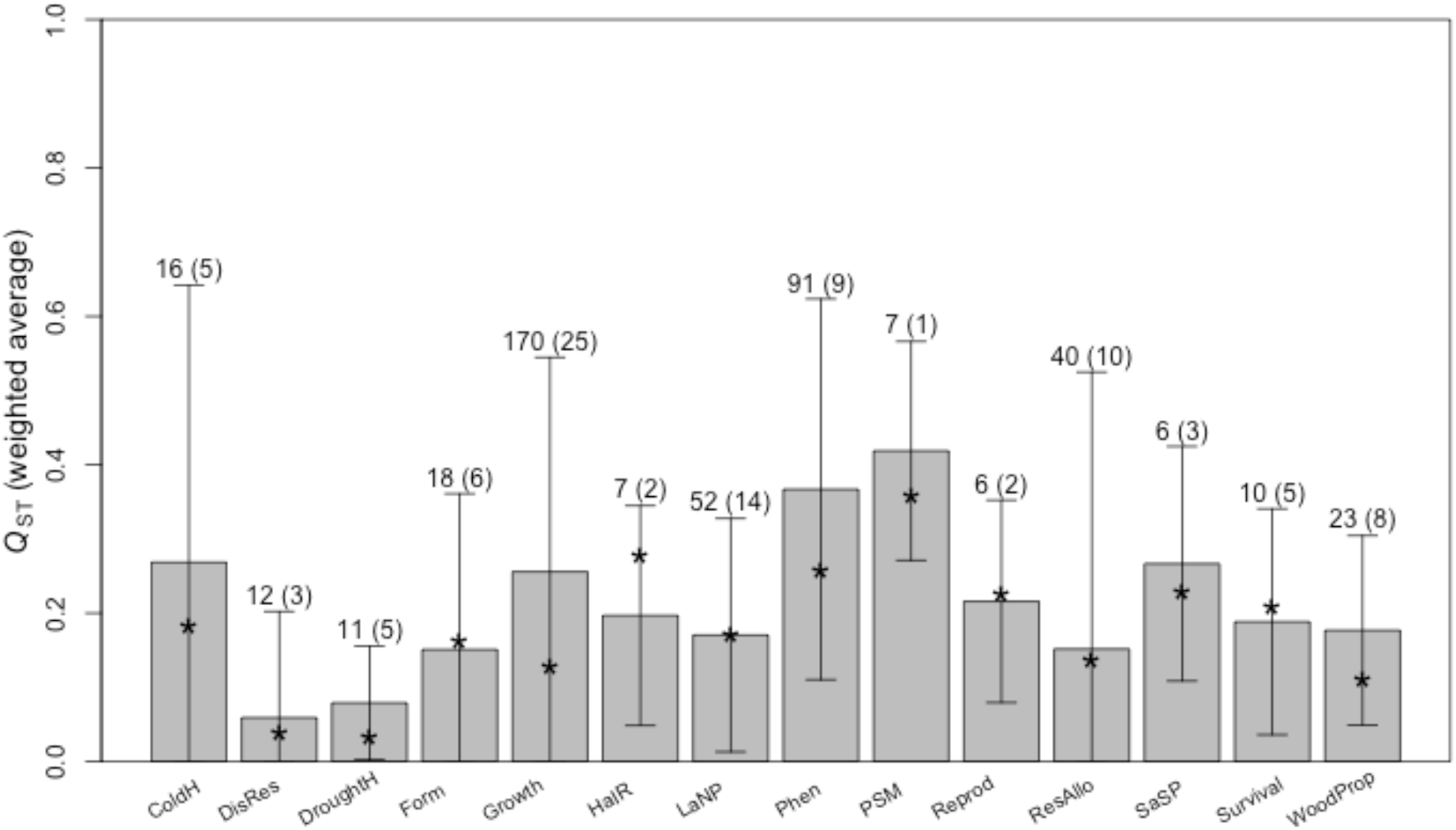
Average *Q*_ST_ for each of 14 trait categories from literature review calculated by weighting each estimate by the number of families used in the estimation. Error bars represent the standard deviation of the weighted averages. Numbers above error bars represent total number of estimates, with total number of unique species in parentheses. Asterisks indicate median values of the unweighted *Q*_ST_ distribution. ColdH = cold hardiness, DisRes = disease resistance, DroughtH = drought hardiness, HaIR = herbivore and insect resistance, LaNP = leaf and needle properties, Phen = phenology, PSM = plant secondary metabolites, Reprod = reproduction, ResAllo = resource allocation, SaSP = seed and seedling properties, WoodProp = wood properties.

Despite neutral genetic differentiation partitioned primarily within populations, adaptive genetic variation seems to be structured to a greater degree across populations, more often than not, for the various fitness-related traits reviewed here. Such a pattern is indeed consistent with local adaptation, assuming that (among other considerations such as the recency of selection) mutation rates are considerably lower than migration rates in these populations (Whitlock 1999; Hendry 2002; Leinonen et al. 2013). In any case, given an extensive literature supporting the local adaptation hypothesis in trees, our results appear consistent with patterns of selective forces acting on abundant, heritable genetic variation across populations, even in the face of gene flow (discussed further in the next section).

## Expectations for the loci underlying quantitative traits

The homogenous environments of the common garden and reciprocal transplant designs are ideally suited to test hypotheses of local adaptation in trees (Sork et al. 2013). However, uncovering the genetic basis and contributory influence of specific loci underlying these adaptive traits is a sizable endeavor on its own, and the success of such pursuits will be determined, in part, by the trait’s underlying genetic architecture (i.e., the number, effect size, type, location, expression, pleiotropic effect, environmental influence, and interaction of underlying loci), which is generally not known *a priori* (Stinchcombe & Hoekstra 2008; Rellstab et al. 2015; Savolainen et al. 2013; Hoban et al. 2016; Burghardt et al. 2017; Wadgymar et al. 2017). Much of our early understanding of the architectures of complex traits came shortly after Nilsson-Ehle (1909) and East (1910) independently demonstrated evidence for multiple-factor inheritance, where Fisher (1918) laid the groundwork for quantitative genetics by incorporating the additive properties of variance to partition phenotypic variation into components tractable to a model of Mendelian inheritance. It was this work, and that of Fisher’s geometric model (1930), which founded the basis for attributing continuous variation of phenotypes to a polygenic model of many underlying heritable components of mainly small effect. From this model, Fisher (1930) concluded that mutations of small effect were the main drivers of adaptation, suggesting large-effect substitutions to contribute little to adaptation due to negative pleiotropic effects constraining effect size. Therefore, the fate of a given locus would be conditioned on its average, marginal effect on fitness calculated across the species, with non-additive deviations from this linear model of inconsequential influence. This micro-mutationist view, to a large extent, remained the dominant thought for nearly half a century (Orr 2005). It was then that Kimura (1983) established that for an allele to contribute to adaptation, it would need to survive the stochastic nature of drift. Thus, new mutations of low frequency and effect were less likely to contribute substantially to adaptive evolution. Considering the adaptive contribution probability of large and small effect loci, Kimura concluded that mutations of moderate effect would be the most plausible. Years later, Orr (1998) showed that over the entire bout of selection via an adaptive walk, the distribution of fixed substitutions resembles an exponential distribution, with effect size decreasing with the proximity to the phenotypic optimum. In addition, the distribution of fitness effects of beneficial mutations is also expected to be exponential (Orr 2003; for more discussion on this aspect, see also Orr 2006; Eyre-Walker & Keightley 2007; Martin & Lenormand 2008, Kopp & Hermisson 2009b; Keightly & Eyre-Walker 2010, Dittmar et al. 2016).

Despite major advances in theory and technology, there still remains substantial uncertainty regarding the exact number of loci underlying many adaptive traits, the effect size distribution of these loci, and how the number of underlying loci and effect distribution may change under various evolutionary regimes (Orr 2001; Slate 2005; Hansen 2006; Mackay et al. 2009). In this section, we describe how various factors can contribute to the (perhaps, effective) number of causative loci, and the distribution of effects underlying continuously distributed adaptive traits, beginning first with aspects of the architecture itself (gene action), and concluding with explanations of how various processes (e.g., selection) play an influential role in the evolution of underlying genetic architectures. Establishing these expectations is essential for assessing common approaches and guiding future directions. In the next section we then compare these expectations with methods used in, and results from, genotype-phenotype associations in trees. While we discuss these examples in isolation, we highlight the fact that the underlying biological processes are often not independent.

### Gene action

The classical genotype-phenotype map is largely one of additive effects, and is represented by a statistical regression of the phenotype on genetic content, as developed by Fisher (1918) and extended by others (e.g., Cockerham 1954; Kempthorne 1954). Indeed, much of the work done in trees has relied on such additive effects to describe heritable and quantitative genetic variation (see previous section). In this model, the phenotypic variance is partitioned into orthogonal (i.e., independent) contributions from the genetic variance (σ_G_), environmental variance (σ_E_), and the variance due to interaction between genotype and environment (σ_GxE_; Figure 2; see Supplemental Box S1). Further, σ_G_ is also the sum of orthogonal variance components, each term representing a different form of gene action. The additive, dominance, and epistatic terms respectfully designate the associated variance contribution of independent alleles, the non-additive contribution to variance of interactions among alleles at the same locus, and the contribution to variance of non-additive interactions among alleles at different loci (the latter of which can take one of many forms such as additive-by-additive, additive-by-dominance, etc.; Lynch & Walsh 1998). As a result, non-additive gene action is minimized as non-linear contributions to the overall phenotype (Moreno 1994; Whitlock et al. 1995) which contributes little to the distinction of the different forms of dominance and epistasis (Cheverud & Routman 1995; Hansen & Wagner 2001; Hermisson et al. 2003; Hansen 2006; Mackay 2014) nor towards the inference of aspects of the underlying genetic architecture in general (Nelson et al. 2013; Huang & Mackay 2016).

**Figure 2.**
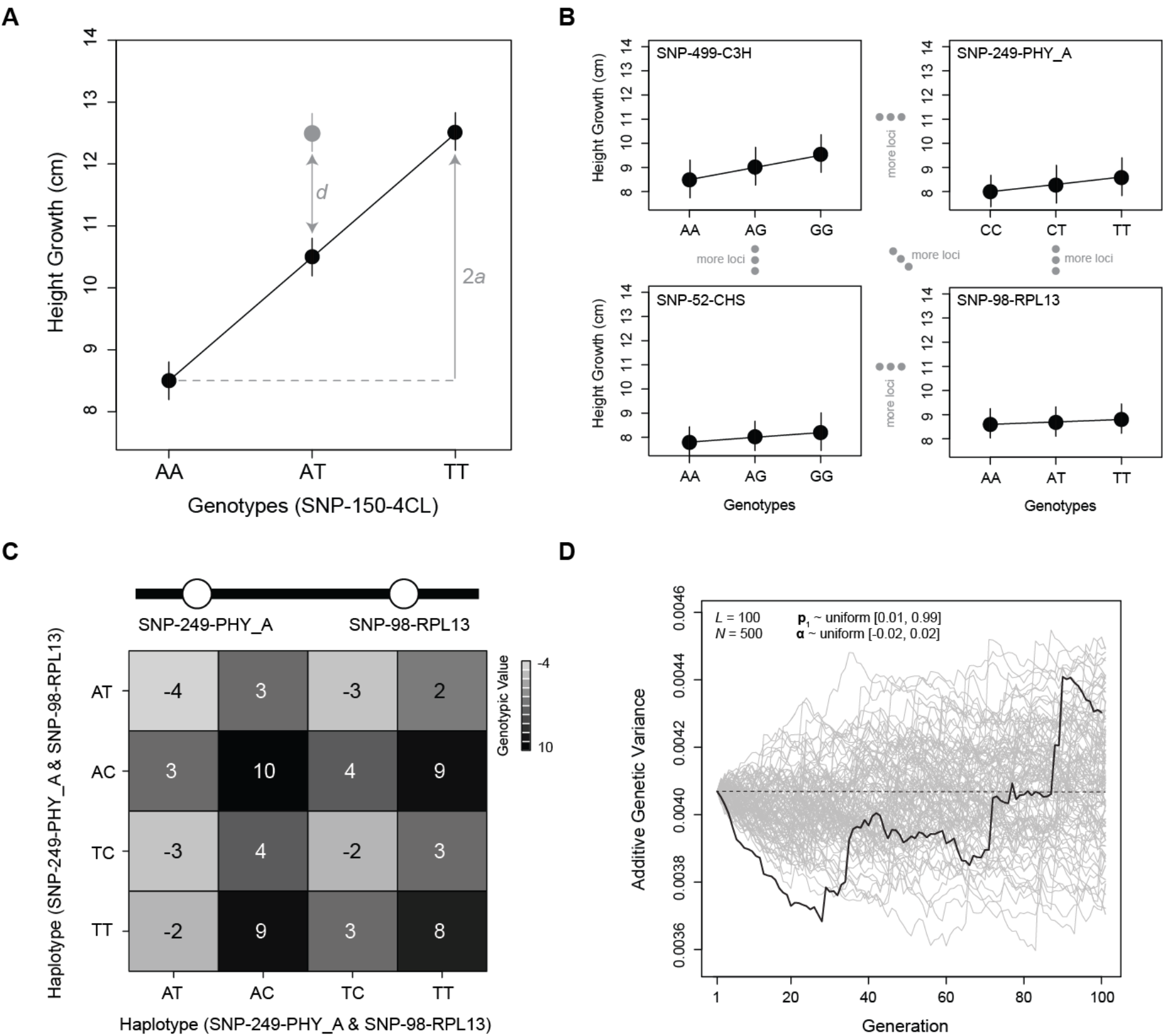
Relevant quantitative genetic concepts are needed to understand the evolution of polygenic traits. (A) Additive and non-additive effects at a single locus, where *a* is defined as the additive effect (also known as the average effect of allelic substitution [α] when there is no dominance) and *d* is defined as the dominance deviation. With dominance, α = a[1 + *k*(*p* - *q*)], where *k* is the degree of dominance (*k* = 0: additive, *k* = 1: dominance, *k* > 1: over-dominance, see Lynch & Walsh 1998). (B) Polygenic traits are determined by multiple genes, each with additive (shown) and non-additive (not shown) effects. The total additive effect is the sum of the additive effects at all causative loci. (C) Additive-by-additive epistasis, where the additive effect of an allele at the PHY_A SNP depends on what allele it is paired with at the RPL13 SNP. In this case, the effects can be thought of as dependent in the following manner using the four possible haplotypes at the PHY_A (A/T SNP) and RPL13 (C/T SNP) SNPs – AC: +5, AT: −2, TC: −1, TT: 4. (D) The effect of genetic drift on the additive genetic variance as determined by 100 independent, causative loci. Each line represents a simulation of genetic drift in a constant sized population (*n* = 500 diploids) conditioned on initial allele frequencies across loci (***p*_1_**) and effect sizes (**α**). The expected mean across all 100 simulations is given by the dashed black line. Any given simulation can deviate strongly from this expectation (solid black line). Thus, when the elements of ***p*** change over time, in this case due to genetic drift, so does the additive genetic variance. See also Supplemental Box S1.

These statistical conveniences afforded by Fisher and others led to the notion that such non-additive effects were transient (i.e., are due to LD, which will decay with the relaxation of selection), or that trends of statistical epistasis were representative of functional epistasis in general, and therefore epistasis was unimportant to evolutionary dynamics (e.g., Bulmer 1980; Crow 2008, 2010; Hill et al. 2008). While minimized in a statistical regression, this does not necessarily mean that epistasis and dominance will not have a profound impact on the genetic architecture, or towards a given population or species’ long-term evolutionary trajectory, even if statistical epistatic or dominance variance is minimal (Goodnight 1988; Chevrud & Routman 1995; Hansen & Wagner 2001; Hansen 2013; Nelson et al. 2013; Griswold 2015; Paixão & Barton 2016). Indeed, parameterizing a model in which the type I sums of squares is determined by non-additive parameters, as opposed to additive variance in the conventional regression model, the majority of genetic variation is still captured by the primary effect in the model regardless of the underlying architecture (Huang & Mackay 2016). Given the prevalence of evidence for non-additive contributions (e.g., Phillips 2008; de Visser et al. 2011; see also references in Hansen 2013), it is likely that non-additive effects will play a role in evolutionary outcomes. For instance, Huber et al. (2017) showed that the degree of dominance in *Arabidopsis* is an outcome based upon functional importance and optimal expression level. Further, Carter et al. (2005) show that, relative to a purely additive trait (or with non-directional epistasis) under directional selection, positive and negative epistasis can respectfully increase or decrease the additive genetic variance, and thus increase or decrease the rate of phenotypic response to selection (see also Le Rouzic & Álvarez-Castro 2016). As Jones et al. (2014) show, for a two-trait phenotype controlled by pleiotropic and epistatic effects, epistasis in the presence of selection can also affect the mutational architecture of complex traits, where the average allelic effect evolves to be negatively correlated with the average epistatic coefficient, the strength of which is greater in larger population sizes. Yet, as described by Barton et al. (2016), and further discussed by Barton (2017) and Paixão & Barton (2016), the infinitesimal model can be generalized to include epistatic effects, particularly when the number of underlying loci is large and selection on individual loci is weak. In the case of non-systematic, weak pairwise epistasis, and without mutation or environmental noise, the infinitesimal model holds to a good approximation (Barton et al. 2016). In the case of sparse epistasis with selection and a large number of loci, the change in the trait mean over 100 generations is greater than that under a purely additive architecture, and the decrease in additive genetic variance exceeds, to an extent, that of the neutral case after about 30 generations (which is exacerbated with simpler architectures), with a reduction of the frequency of segregating alleles with positive effect on the trait (Barton et al. 2016; Barton 2017).

Despite an ongoing debate within the literature (Wright 1932; Whitlock 1995; Crow 2008, 2010; Gibson 2012; Zuk et al. 2012; Hansen 2013; Hemani et al. 2013; Nelson et al. 2013; Mäki-Tanila & Hill 2014; Ávila et al. 2014; Paixão & Barton 2016), and given that there seems to be no general prevalence of either positive or negative epistatic interactions (Mackay 2014), the infinitesimal model is likely to continue to contribute to our understanding of the evolution of complex traits, as exemplified in its application towards breeding applications (Turelli & Barton 1994) and specifically those successfully applied to trees (Savolainen et al. 2007; Thavamanikumar et al. 2013; Isik et al. 2015; Grattapaglia 2017). Ultimately, the success of such models will be conditioned on the context, as well as the distinction between physiological and statistical gene action. Here, (higher order) non-additive contributions to phenotypic variance will likely have minimal deviations from the limit of the infinitesimal model in the shortterm, particularly if this is primarily due to independent, low-order interactions, and should thus be applied with this in mind. As such, while short-term evolutionary processes are likely to hold in this limit, identifying the non-additive loci which underlie the trait, and their respective gene action, may still need further inquiry (Grattapaglia 2017). Indeed, it is often argued that nonadditive gene action is too often neglected in studies of complex traits (e.g., Carlborg & Haley 2004), possibly due to the large sample sizes required to detect significant interactions, and lack of statistical power incurred due to multiple hypothesis testing (Mackay 2014). Given the recent reduced cost of sequencing technology and availability of novel computational and laboratory tools, future studies incorporating investigations of epistasis and dominance (where appropriate and feasible) would contribute to our understanding of genetic architectures, quantitative trait evolution, and breeding applications in trees (Vitezica et al. 2017). For example, breeding applications assessing hybridization across divergent backgrounds, as is also prevalent across species in nature, have shown the importance of non-additive effects in phenotypic outcomes (as in *Eucalyptus,* e.g., Tan et al. 2017, and *Pinus,* e.g., Dungey 2001). Even so, the additive model is still a powerful tool to describe the loci underlying adaptive traits.

Pleiotropy is another considerable factor influencing the expectations of the genetic architecture of quantitative traits, its evolution or evolvability, and indeed the genotype-phenotype map (Hansen 2003; Orr 2006; Chevin et al. 2010b; Tenallion 2014). While multiple definitions exist across the literature (see Paaby & Rockman 2013), pleiotropy is generally identified as a single locus influencing multiple phenotypic traits. Other than linkage disequilibrium, pleiotropy is the fundamental cause of genetic covariance among phenotypes (Lande 1980). Given that the number of independent traits under selection is likely limited (Barton 1990), pleiotropy likely plays a substantial role in evolutionary dynamics. It is expected that as the number of traits, *n*, influenced by a locus increases, the probability of a beneficial mutation will decrease with the effect size of a mutation; where the effect size, *r,* relative to the distance to the phenotypic optimum, *d · n*^−1/2^, must be (much) less than *d* in order to be beneficial (Fisher 1930; the so-called ‘cost of complexity’: Orr 2000). Yet, empirical data seem to contradict this hypothetical cost, as the effect size of mutations often do not scale with pleiotropy in this way, and instead increase with the dimensionality of targeted traits (Wagner et al. 2008; Wang et al. 2010). Additionally, universal pleiotropy, where all mutations affect all phenotypes, and where there is no net directionality of mutations (i.e., mutational isotropy; both aspects as in Fisher 1930), has also been challenged by findings which suggest that only a fraction of phenotypic traits are affected by pleiotropic loci (Wagner et al. 2008; Wang et al. 2010). Relaxation of such assumptions from Fisher’s geometric model have shown that the total number of traits affected by pleiotropy has a relatively decreased effect on the rate of evolution in more general models (e.g., Martin & Lenormand 2006; see also Simons et al. 2017, and references in Wagner & Zhang 2011 and Tenaillon 2014). It seems that if model organisms (e.g., Pickrell et al. 2016, Smith 2016) are taken as a bellwether for expectations in trees, pleiotropy is likely a contributing factor for many quantitative traits. Thus, the fraction of beneficial mutations is likely limited when the number of traits influenced is large, suggesting that the cost of complexity (or, more precisely, pleiotropy) may be generally robust (Welch & Waxman 2003), particularly when a population is close to its phenotypic optimum where selection acts against dimensionality of pleiotropic effects (Zhang 2012). Thus, the degrees of pleiotropy across underlying loci, distance from phenotypic optima, and covariance among traits under selection can have profound effects on evolutionary outcomes (e.g., as in *Pinus contorta,* Lotterhos et al. 2017). This is particularly true for the evolvability of architectures and distribution of effect sizes, which further depends on the variational autonomy of the traits affected by pleiotropy and the modularity of mutations, the former of which is ultimately determined by the direction and size of effect among a set of pleiotropic loci across a set of characters (see Arnold 1992; Wagner & Altenberg 1996; Hansen 2003, 2006; Wagner et al. 2007; Chevin et al. 2010b; Wagner & Zhang 2011; MacPherson et al. 2015).

In many investigations of local adaptation, the primary interest is in trait evolution and thus the underlying genetic components. As such, environmental effects and interactions are not often pursued, or perhaps even detected (Yoder & Tiffin 2017), particularly in studies of a single common garden or environment, and are instead treated in much the same way as epistatic interactions discussed above. Nonetheless, genotypic effects can evolve through genotype-by-environment interactions with a changing environment just as is the case for the evolution of non-additive interactions with a changing genetic background (Hansen 2006). Indeed, it is likely that consistent fluctuations in the environment would select for environmentally-perceptive responses, which seems to be the case across many tree species (Li et al. 2017). The contribution to the effect size distribution from GxE interactions will be a function of the variation in selection across the environments experienced by the interacting allele(s) as well as the level of gene flow between environments and fitness differences among various genetic backgrounds, but to our knowledge such information (to the extent of that for e.g., selective sweeps) is lacking within the literature.

### Negative selection

Negative selection acts against deleterious mutations that arise within populations. It is one, but not the only, mechanism that underlies stabilizing selection, defined at the level of the phenotype where deviations from an optimal value are selected against. Optima in this framework can be thought of either globally (i.e., across all individuals) or locally (i.e., individuals within a population), where the latter can have varying optima across populations. The nature of the optima (i.e., being local or global) affects the detectable trait architecture. For example, trait architecture should be composed of rare alleles with a negative relationship between effect size and allele frequency (cf. Eyre-Walker 2010 and references therein), where this relationship can also be confounded with degree of dominance and gene expression network connectivity (Huber et al. 2017), under models of a single global optimum. From a population genetic perspective, the ubiquity of negative selection is encapsulated in the name background selection, which has extensive reviews about its presence in natural systems (Charlesworth 2013), its importance for the neutral and nearly neutral theories of molecular evolution (Ohta 1992, 1996), and its contribution to observable patterns of hitchhiking (Stephan 2010). Important for the study of polygenic adaptation and its architecture, however, is that loci identified using GWAS may also include segregating deleterious variation (as argued and hinted at in Eckert et al. 2013b; cf. Yang et al. 2017; Gazal et al. 2017) as this creates trait variance, with little known about their prevalence (including differential prevalence across traits), differentiation in frequencies across populations (but see Zhang et al. 2016), and effects on downstream inferences about divergent selection pressures across populations. It is sets of GWAS loci, though, that are currently analyzed for signatures of local adaptation via spatially divergent (i.e., locally positive) natural selection (e.g., Berg & Coop 2014).

Recent exemplary work with expression networks in *Populus tremula* L. (Salicaceae; Mähler et al. 2017) and the herbaceous *Capsella grandiflora* Boiss. (Brassicaceae; Josephs et al. 2015, 2017a) have revealed intriguing insight into the effects of negative selection on the architecture of complex traits in plants, as well as the relationship between network connectivity and the strength of negative selection. In *P. tremula*, genes with expression levels that were significantly associated with sequence variation were found more often in the periphery of the co-expression network (lower network connectivity) than within network module hubs (higher connectivity), while expression-associated SNPs were negatively correlated with network connectivity and effect size, a pattern also found between connectivity and expression variance, and minor allele frequency and QTL effect size (Mähler et al. 2017). Genes associated with sequence variation had less skewed site-frequency spectra (i.e., the frequency distribution of allelic variants) and lower estimates of nonsynonymous to synonymous divergence (d_N_/d_S_) than genes not associated with sequence variation, together suggesting that genes within the periphery of co-expression networks are likely under less selective constraint than those genes with high network connectivity which likely experience greater intensities of purifying selection. These genes thus tend to have more segregating variation and may be those most likely to be detected with current sample sizes utilized in GWAS, which has implications for estimation of trait architecture and its ‘degree’ of polygenicity. Even so, while there is prevalent evidence of negative selection in trees (e.g., Krutovsky & Neale 2005, Palmé et al. 2009, Eckert et al. 2013a, b; De La Torre et al. 2017), more inquiry is needed.

### Positive selection

The temporal and spatial heterogeneity of selection can impact the evolution of genetic architectures underlying adaptation. These impacts are often thought of on a spectrum of tradeoffs, with one end being antagonistic pleiotropy where allelic effects vary between positive and negative on fitness across populations, and the other being conditional neutrality where allelic effects on fitness are positive in one or more populations and nearly zero in others (Anderson et al. 2012, Savolainen et al. 2013). For instance, alleles incorporated into a population after a shift in environmental influence can increase from low to high frequency via positive selection. The existence of such a beneficial allele can manifest in several ways: from new mutations, introgression through gene flow, or molecular reorganization through novel recombination, inversion, transposition, copy number variation, or insertion-deletion events. If there is strong selection acting on this allele (*N*_e_*s* >> 1), it will sweep to high frequency creating a signature of reduced polymorphism at neutral sites physically linked to the allele (‘genetic hitchhiking’, Maynard Smith & Haigh 1974) resulting in a hard ‘selective sweep’ (Berry et al. 1991). However, in structured populations with limited gene flow, this process can take significantly longer to reach fixation, resulting in incomplete sweeps (Whitlock 2003). Additionally, Pavlidis et al. (2012) found that, in congruence with Chevin & Hospital (2008), a multilocus genotype often prevents the trajectories of individual alleles from sweeping to fixation, with an increasing number of loci leading to decreasing probability of fixation, and as a result, an altered selective signature at such loci (see also Jain & Stephan 2017). As such, hard selective sweeps in a polygenic architecture are expected to be rare (but not completely absent) under most circumstances, particularly when the shift in environment causes a relatively small deviation from the phenotypic optimum. Thus, hard sweeps most likely apply to loci with relatively large effect above a calculated, context-dependent threshold value (Orr 2005; de Vladar & Barton 2014; Stephan 2015; see specifically Jain & Stephan 2015, 2017).

While early literature (Maynard Smith & Haigh 1974; Kaplan et al. 1989) focused on the rapid sweep of an allele incorporated into a population after an environmental shift, research within the last few decades have focused on ‘soft sweeps’ resulting from neutral or deleterious mutations that are present in the standing genetic variation prior to the change in the selective environment, wherein the selection coefficient changes with the environmental shift such that the allele(s) become evolutionarily advantageous (reviewed in Hermisson & Pennings 2005, Barret & Schluter 2008, Messer & Petrov 2013, and Hermisson & Pennings 2017; see also Jensen 2014). These allele(s) could manifest via a single low-frequency variant, multiple variants caused by parallel recurrent mutation/reorganization on multiple haplotypes, or multiple unique alleles that arise independently within, perhaps multiple, populations. In such cases where selection acts via soft sweeps, the rate of evolution at the phenotypic level is expected to exceed those of hard sweeps because the alleles under selection have escaped the stochastic nature of drift to a greater degree and are segregating within multiple individuals and genetic backgrounds within the population. The extent to which soft sweeps alter the effect size distributions underlying the genetic architecture is likely dependent upon both the strength of selection and effect size before and after the environmental change (Messer & Petrov 2013; Matuszewski et al. 2015; Jain & Stephan 2017), while the frequency before selection influences the likelihood of subsequent detection (Innan & Kim 2004). Additionally, if multiple mutations are segregating during the sweep, the probability of fixation for any given locus also decreases (Pennings & Hermisson 2006a, 2006b; Chevin & Hospital 2008; Ralph & Coop 2010). Evidence for hard sweeps in tree species exist within the literature, although they are rare (e.g., disease response genes in *Pinus taeda* Ersoz et al. 2010; see also Table 2 in Siol et al. 2010). However, for many species of trees, which often experience high gene flow and strong diversifying selection across populations, adaptive divergence for polygenic traits is expected to result more often from soft sweeps than hard sweeps, affecting phenotypes by subtle allele frequency changes across populations, such that allele frequency differences of individual loci across populations for neutral and selective sites will often be nearly indistinguishable (Latta 1998, 2003; Barton 1999; Le Corre & Kremer 2012; Stephan 2015; Yeaman 2015; Jain & Stephan 2015, 2017). Indeed, the large effective population sizes found in most tree species would permit large effective mutation rates (or reorganization events) necessary for a soft selective sweep from multiple unique variants, particularly when the phenotype is underlain by a large mutational target. Even so, and as highlighted by Stephan (2015) and Bailey & Bataillon (2016), the extent to which scientists can detect the influence of demographic processes on soft versus hard sweeps, and vice versa, remains challenging (Jensen et al. 2005; Chevin & Hospital 2008; Schrider et al. 2015, 2016; Schrider & Kern 2016; Hermisson & Pennings 2017).

While discrete directional selection events are likely to be a common evolutionary influence across taxa, fluctuating or sustained directional selection (i.e., moving optima) are also likely to be contributory factors influencing the genetic architecture of quantitative traits (reviewed in Kopp & Matuszewski 2013; see also McCandlish & Stoltzfus 2014). For a sustained moving optimum, the effect size distribution of beneficial alleles is expected to be dependent upon the effect distribution of standing or *de novo* mutations as well as the strength of selection: if the rate of change is dramatic, adaptation from new mutations is expected to occur through intermediate to large-effect loci (Kopp & Hermisson 2009a; Matuszewski et al. 2014) or from small-effect loci when adaptation occurs via standing variation (particularly when epistasis is considered, Matuszewski et al. 2015). Under lesser rates of environmental change, adaptation is expected to proceed through mainly alleles of small-effect (Collins et al. 2007; Kopp & Hermisson 2009a, 2009b) where intermediate effects will dominate the long-term distribution of effect sizes (Kopp & Hermisson 2009b). In the case of fluctuating environments, outcomes often depend directly on the degree of temporal autocorrelation of the changing environment. In such cases of stochastic fluctuation around a linear trend of environmental change, extinction risk increases relative to that of the strictly linear trend (Bürger & Lynch 1995) where local adaptation lags, to some degree, behind any given contemporaneous scenario. In comparison, and similar in some ways, stochastic fluctuations around a constant mean are expected to resemble the dramatic environmental change scenario described above, characterized by strong selection pressures, maladaptation between generations, and a large lag load (Bürger 1999; Chevin 2012; Kopp & Matuszewski 2013). In the case of autocorrelated shifts, the ‘predictability’ of such fluctuations may decrease the possibility of extinction, increase probability of local adaptation, and lead to similar scenarios as discussed for gradual changes in the environment (Kopp & Matuszewski 2013).

### Gene flow

Gene flow, to the extent that would be appreciable to that found in trees (reviewed in Savolainen et al. 2007), is also an important component shaping quantitative expectations. Indeed, since the early 1900s we have known that gene flow can disrupt adaptation if selection is not strong enough to overcome the loss of beneficial alleles (Haldane 1930; Wright 1931; Slatkin 1987; reviewed in Felsenstein 1976, Lenormand 2002, Savolainen et al. 2007, 2013, Feder et al. 2012a, and Tigano & Friesen 2016). Particularly when gene flow is asymmetric between core and peripheral populations, adaptation can be inhibited in marginal habitats (Kirkpatrick & Barton 1997; Kawecki 2008). Even so, there is abundant evidence that gene flow can promote adaptation and maintain polymorphisms within populations, including white sand lizards (Laurent et al. 2016), stick insects (Comeault et al. 2014, 2015), cichlid fishes (Meier et al. 2017), Darwin’s finches (Lamichhaney et al. 2015), and lodgepole pine (Yeaman & Jarvis 2006).

The magnitude of gene flow between populations can also impact the distribution of effect sizes, for when gene flow falls below a critical threshold, and over many thousands of generations, there is an increase in the probability of establishment and persistence times of large-effect alleles, thus reducing the proportion of the polymorphism due to small-effect loci (Yeaman and Otto 2011; Yeaman and Whitlock 2011). These dynamics are further influenced by the susceptibility of alleles to ‘swamping’ (Slatktin 1975; Bürger & Akerman 2011; Lenormand 2002; Yeaman 2015; *sensu* Haldane 1930). For alleles that are prone to swamping, adaptive phenotypic divergence depends on genetic variation and is driven by allelic covariance among populations particularly when the underlying architecture is highly polygenic, the mutation rate is high, and the number of loci underlying the trait exceeds the number needed to achieve the local optimum phenotype (genetic redundancy; Yeaman 2015). Conversely, when there is little genetic redundancy underlying the trait, limited divergence is observed unless the effect size of a given swamping-prone allele exceeds the critical migration threshold. In these cases where swamping-prone alleles contribute to adaptive divergence, the genetic architecture is transient and any given locus contributes ephemerally to phenotypic divergence, even for loci of relatively large effect (Yeaman 2015). In the case of swamping-resistant alleles, the evolved architecture is enriched for large-effect loci and adaptive divergence can be maintained with little genetic variation or input from mutation. Yet while the contribution from such loci can last many thousands of generations, the architecture can again become transient as the genetic redundancy or mutation rate increases (Yeaman and Whitlock 2011; Yeaman 2015).

Physical linkage and reduction of recombination between adaptive loci can also play a considerable role in adaptive processes in the face of gene flow (Feder & Nosil 2010; Feder et al. 2012a, b; Yeaman 2013; references therein). In such cases, loci that are tightly linked to other loci already under selection will have an increased probability of contributing to local adaptation, both because of physical linkage as well as by reducing the effective recombination among loci within the sequence block. For instance, Yeaman & Whitlock (2011) showed that under divergent selection with gene flow, the number of contributing loci decreases with increasing recombination while small effect loci tend to cluster in groups that act as a single large effect locus (see also Remington 2015), and strong selection can maintain these clusters of linked loci over greater map distances than can weak selection. More recently, Yeaman (2013) employed individual-based simulations to provide evidence that the clustering of alleles throughout a bout of adaptation is unlikely to be driven mainly by divergence hitchhiking alone, and that instead competition between genetic architectures and chromosomal rearrangements occurring throughout adaptive processes under a range of environmental fluctuation scenarios can lead to the evolution of tightly clustered adaptive loci which persist in the event of gene flow, unlike the clusters identified by Yeaman & Whitlock (2011). Yeaman (2013) found that the level of clustering was a function of the temporal fluctuation period, the rate of rearrangement itself is an important determinant on the evolution of clustered architectures, and clusters can in some cases be evolutionarily disadvantageous. Together, these results suggest that genomic rearrangements (reviewed in Ortiz-Barrientos et al. 2016), including inversions (Kirkpatrick & Barton 2006; reviewed in Hoffman & Rieseberg 2008), which decrease the effective rates of gene flow among adaptive sequences can be an essential component of local adaptation, and indeed some cases of speciation, in the face of gene flow.

### Summary

While we provided an overview of the factors that can influence the genetic architecture of local adaptation, we acknowledge that it is far from exhaustive. Because the phenotypes used in studies of local adaptation (particularly those assumed or corroborated to be a component of total lifetime fitness) often have a continuous distribution, and are thus quantitative in nature, the underlying genetic basis for these traits is likely polygenic and is predicted to be underlain by multiple (often many) segregating loci, many of which may confer small phenotypic effects (and are thus unlikely to be detected using single-locus approaches). Even so, a continuum exists, where the true genetic architecture (the number of contributing loci, as well as their relative locations within the genome, phenotypic effects, and interactions) underlying a given complex trait is itself determined by a combination of evolutionary forces that encompass an interplay between the strength, timing, and direction of (background) selection against the homogenizing effects of gene flow and recombination, disruptive effects of drift, linkage, transposition, inversion, and mutation, interactions between underlying loci as well as between these loci and the environment, structural variation, relationship to gene expression networks, as well as other factors related to life history. Consequently, the contemporary genetic architecture is a result of past evolutionary processes, while the adaptive response to future evolutionary dynamics is influenced in part by the contemporary architecture and genetic variance at hand.

## The genomics of local adaptation in trees

### Common approaches used to identify adaptive loci

Across taxa, and specifically in trees, the predominant association and outlier methods for uncovering sets of loci underlying local adaptation have relied upon single-locus population genetic approaches. Putatively adaptive loci are often identified by elevated allele frequency differences among populations relative to patterns genome-wide. Yet, as revealed in the previous section, loci underlying polygenic traits will often be indistinguishable from non-causative sites in this way. Further, outlier tests based on *F*_ST_ (*sensu* Lewontin & Krakaur 1973) do not incorporate information regarding putative phenotypic targets of selection nor environmental drivers of differentiation, often do not correct for neutral population structure (but see Lotterhos & Whitlock 2015), and will inevitably isolate a biased set of candidate loci (Hermisson 2009; Cruickshank & Hahn 2014). In the case of single-locus genotype-environment associations (reviewed in Rellstab et al. 2015; see also De Mita et al. 2013), information about possible environmental drivers is incorporated by assessing the association between allele frequencies and environmental heterogeneity, yet without information regarding traits hypothesized to be influenced by selection (Schoville et al. 2012). Single-locus genome wide association studies (see next section; Supplemental Box S2) and quantitative trait loci (QTL) experiments (reviewed in Ritland et al. 2011, Hall et al. 2016) have also been used in trees, quantifying the differential effects of typed alleles on a given phenotype. Despite the shortcomings of these methods, such studies provide candidate loci that can be investigated in further detail (Tiffin & Ross-Ibarra 2014), which is particularly advantageous when resources are limited. Indeed, as discussed below, these approaches dominate the methods used to uncover complex traits (adaptive or otherwise) in trees.

### Current progress in trees

In light of the expectations outlined above for the architecture of quantitative traits under various evolutionary regimes, and the methods commonly used to detect these loci, we reviewed the literature of single-locus genotype-phenotype associations (GPAs, which included associations to gene expression levels) from studies in forest trees. In doing so, we identified 52 articles across 10 genera and 24 species with a total of 2113 GPAs (Supplemental Table S2, Supplemental File F2). Because most studies in trees do not report phenotypic effect sizes of individual loci (i.e., regression coefficients), we report *r*^2^ values which can be used to quantify the percent phenotypic variance explained by the associated locus. In cases where multiple SNPs from a given locus (e.g., a gene or scaffold) were associated to a trait, we averaged the *r*^2^ values for that locus. As with our review of trait heritability and *Q*_ST_, we grouped phenotypic traits used in associations into twelve broad categories (in this case, no phenotypes fell into Survival or Seed and Seedling Properties groups). If traits important to tree conservation and industry are often of a polygenic basis, we would expect small to moderate effects from loci empirically associated to phenotype. Indeed, across the trait groups considered here, the mean *r*^2^ was 0.039, where 80.79% (n = 1707) of recorded estimates had *r*^2^ values less than 0.05, 18.78% (n = 397) of *r*^2^ values falling between [0.05,0.22], and nine values of *r*^2^ greater than 0.22, which were all related to *Cronartium ribicola* resistance in *Pinus monticola* Douglas ex. D. Don (Figure 3a).

**Figure 3A.**
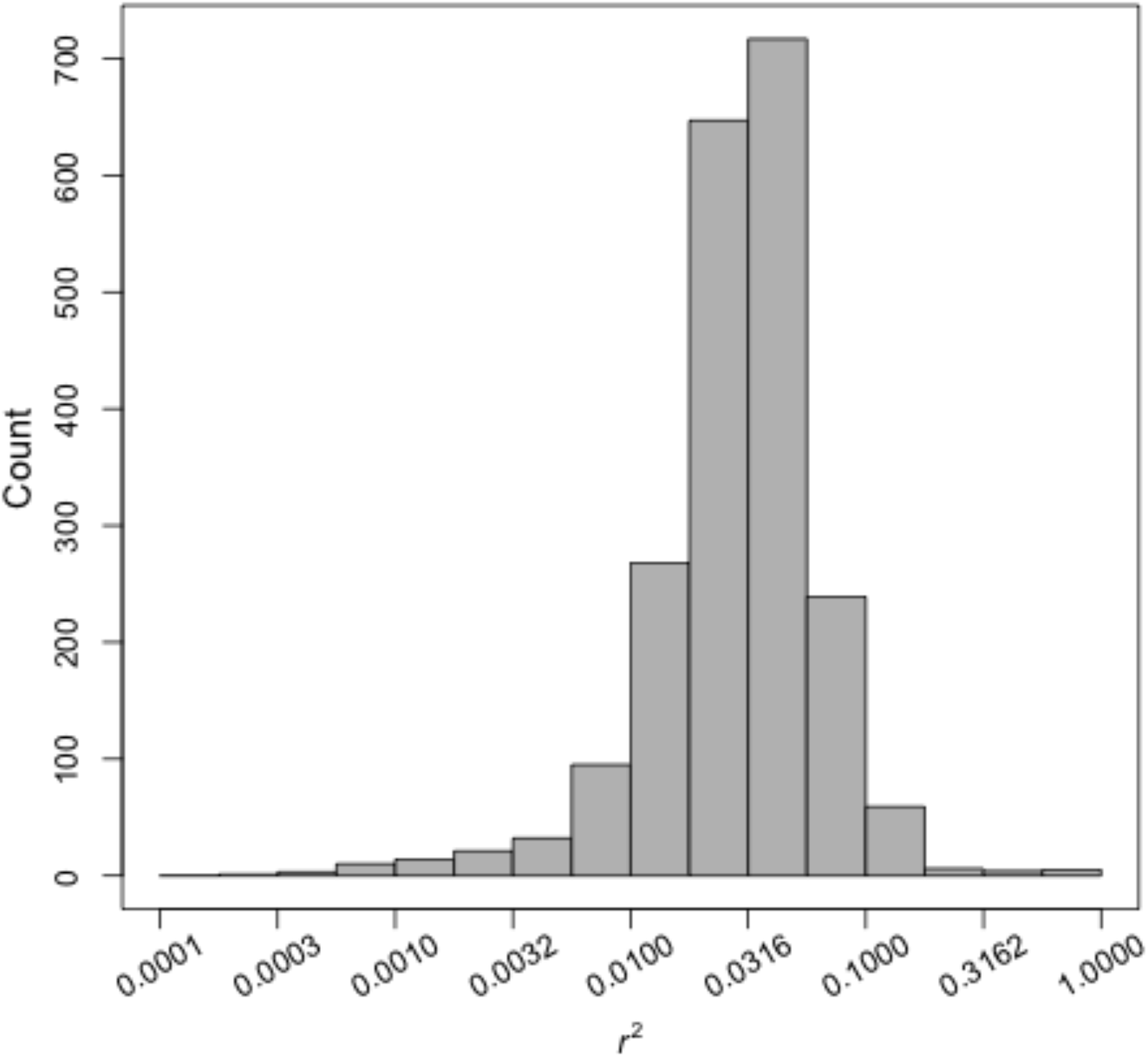
Counts of per-locus percent variance explained (*r*^2^) estimates from single-locus genotype-phenotype associations from literature review. Note logarithmic x-axis.

Of the twelve trait groups, all but those traits relating to both reproduction and herbivore and insect resistance had *r*^2^ estimates greater than 0.10, with traits relating to disease resistance, growth, leaf and needle properties, phenology, and wood properties each contributing over 10% of these outliers (Figure 3b). These small effects tend to also not account for much of the observed heritability, but can explain sizeable fractions in some instances (e.g., primary metabolites in Eckert et al. 2012). Of the loci associated with expression levels, *r*^2^ estimates were between 0.05 and 0.152 in all but one case (n = 54). We also assessed the propensity of individual loci to be associated to more than one phenotype or expression level across our literature review. Without correcting for the multiple associations of a locus to yearly phenotypes (e.g., bud flush 2009, bud flush 2010), we found that the average number of loci associated to multiple phenotypes per study was 6.00, while after correcting for multiple years the average number decreased to 5.42. The median number of SNPs utilized for association per study was 206, where 75% (39/52) of studies used less than 1,000 SNPs, eight studies using between 1,000-10,000 SNPs, four studies using between 29,000-35,000 SNPs, and one study utilizing 2,822,609 SNPs for association (all studies with greater than 10,000 SNPs were from either *Pinus* or *Populus* species).

**Figure 3B.**
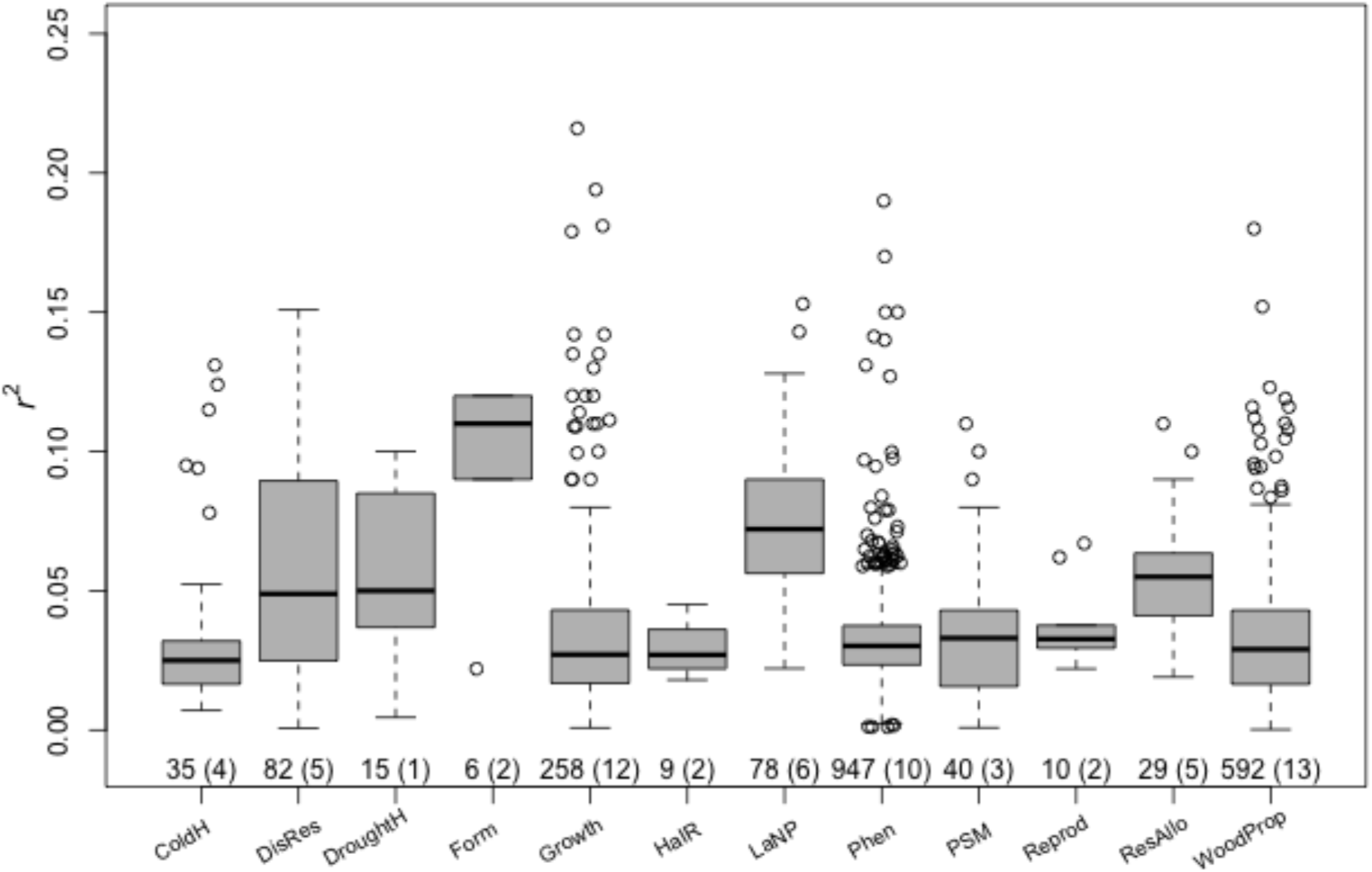
Distribution of per-locus percent variance explained (*r*^2^) values for trait groups within genotype-phenotype literature review. Values along x-axis are total number of estimates and number of species across estimates. Not shown are nine outliers for disease resistance to *Cronartium ribicola* in *Pinus monticola* (range = [0.402, 1.0]) from Lui et al. 2017. Abbreviations as in Figure 1.

### Are we out of the woods yet?

From insight gained from the literature review of genotype-phenotype associations it seems that the vast majority of the genetic architecture of local adaptation and complex traits in trees remains largely unexplained using common GWAS methods (see also Box 1), a consistent pattern across the past decade of research in trees (Neale & Savolainen 2004; Savolainen et al. 2007; Ćalić et al. 2015; Hall et al. 2017). Furthermore, it is likely that the estimates for percent variance explained are inflated due to a combination of QTLs that break down into smaller effect loci (Remington 2015), the Beavis effect (Beavis 1994; Xu 2003), and the Winner’s Curse (Görning et al. 2001; Zöllner & Pritchard 2007) where locus effects are inflated by using the same data for both gene identification and phenotypic prediction (see Box 1 in Josephs et al. 2017b for a detailed synopsis of these biases). Such a pattern suggests that, indeed, many of the traits important to evolutionary, breeding, and conservation insight in trees are likely of a polygenic basis and that future studies must take this into account when seeking to identify the underlying loci.

Even within studies of model organisms, missing heritability is nothing new. Across taxa, missing heritability is less frequent within phenotypes of mono-to oligogenic bases (as seen for the *Cr2* major-gene resistant locus in *Pinus monticola,* Liu et al. 2017), as would be expected, and is a recurrent, pervasive shortcoming from genotype-phenotype associations of complex traits, particularly those maintaining single-locus perspectives. A number of explanations have been put forth to explain the missing heritability, such as epistasis (Hemani et al. 2013) and its inflationary effect on heritability estimates (Zuk et al. 2012), environmental or epigenetic interactions (Feldman & Lewontin 1975) as well as their inflationary effect on heritability estimates (Zuk et al. 2012), (unmeasured) low-frequency variants of large effect (Dickson et al. 2010), genetic or variance heterogeneity of individual alleles (Leiserson et al. 2013; *cf.* Box 1 in Nelson et al. 2013), or common variants with effect size below detection thresholds (Yang et al. 2010). As such, here we avoid supporting one causative hypothesis over another, particularly given the ongoing discussion within the literature, for which strengths and weakness for any viewpoint are apparent (e.g., Gibson et al. 2010), and because of the progress yet to be made in trees.

Indeed, the dissection of the genetic architectures underlying complex traits in trees is still in its nascency compared to the progress of model organisms (for which missing heritability is still an issue), and beyond issues of coverage, genomic saturation, and genomic resources (discussed below in The Path Forward), we must approach this issue with all possibilities in mind. Given the unique properties of the life histories, genome size and organization of many tree species, and the limited numbers of studies with large sets of molecular markers, causative sources of the missing heritability should be ruled out, or supported, as with any other hypothesis, particularly as we gain information from contemporary studies of trees that address shortcomings of those in the past. Further, we must keep in mind differences between functional and statistical gene action (Álvarez-Castro et al. 2007; Nelson et al. 2013; Huang & Mackay 2016; Huber et al. 2017). In any case, it seems that sample sizes of single-locus approaches will need to be increased (Hall et al. 2016), albeit with diminishing returns (Boyle et al. 2017; Simons et al. 2017), to discover a higher proportion of the underlying loci in trees due to small to moderate additive effects. Alongside suggestions outlined in The Path Forward, incorporating investigations into such aspects of epistasis, dominance, pleiotropy, expression, GxE effects, and network analyses (when appropriate), may be a worthwhile complement (e.g., Lotterhos et al. 2017, Mähler et al. 2017, Mizrachi et al. 2017; Tan et al. 2017).

While the infinitesimal model will continue to prove to be immensely useful for breeding programs and for short-term evolutionary predictions, and we may find that the missing heritability in trees is truly due to consequences of the infinitesimal regime (as is often cited to be the majority consensus across taxa for missing heritability), it has been argued that the analysis paradigm for such studies is near its limits in describing the functional genetic architecture of quantitative traits, and that it is therefore necessary to move beyond single-locus perspectives and reconsider common practices (Pritchard & Di Rienzo 2010; Nelson et al. 2013; Sork et al. 2013; Tiffin & Ross-Ibarra 2014; Wadgymar et al. 2017). At this stage, it seems that we investigators seeking to describe the genetic architecture of quantitative traits in trees have some ways yet to go before we are truly out of the woods. In the next section, we describe the path forward to describing genetic architectures from a polygenic and functional perspective, identify resources available to advance our knowledge and fill knowledge gaps, as well as future directions for this research area.

## The Path Forward

As we have outlined, there is still ample room for improvement in our description and understanding of the genetic architecture of quantitative traits in trees (see Table 1 and Box 1). Importantly, methods used to uncover causative loci should take into consideration the expected degree of polygenicity, the relative contributions of various forms of gene action, as well as how past evolutionary phenomena has likely shaped current adaptive expectations. In this section, we orient our path forward by first highlighting utilities available to, and underused within, the forest genetics community to describe the genetic architecture of complex traits. We then outline several suggestions to facilitate further progress and advocate for prospective perspectives in future studies such that information and data may continue to be used easily in subsequent syntheses across pathways, environments, species, and towards insight to identify future needed resources as our understanding progresses. While our recommendations are specific to the tree community, we also acknowledge other valuable recommendations from recent reviews (e.g., Savolainen et al. 2013; Tiffin & Ross-Ibarra 2014; Lotterhos & Whitlock 2015; Gagnaire & Gaggiotti 2016; Hoban et al. 2016; Wellenreuther et al. 2016; Burghardt et al. 2017; Wadgymar et al. 2017).

### Stepping off the path – what’s in our pack?

The genetic architecture underlying local adaptation and complex traits likely has a polygenic basis composed of many loci of relatively weak effect yet many of the common association or outlier methods will often fail to detect many of the causative loci of small to moderate influence. Such investigations have so far led to an incomplete description of studied architectures, and, in many cases, have limited our understanding of complex traits in trees to a handful of loci. While we do not advocate that such single-locus methods be avoided in future studies (considered further in the next section), here we outline underused and promising approaches to identify and describe underlying loci that explicitly take into account the polygenic basis of such traits and may help advance our understanding in future studies, including some of the questions we have outlined in Table 1. Multivariate, multiple regression, and machine learning techniques are three such examples, and differ from univariate analyses by analyzing patterns among multiple loci simultaneously.

The Bayesian sparse linear mixed model (BSLMM), for instance, such as that deployed in the software package `GEMMA` (Zhou et al. 2013), is developed for both genomic prediction (see also Box 1) and mapping of complex traits that offers considerable advantages over single-locus genotype-phenotype approaches (Guan & Stephans 2011; Ehret et al. 2012; Zhou et al. 2013; Moser et al. 2015). This analysis has gained in popularity recently, being used across diverse taxa such as stick insects (Comeault et al. 2015, Riesch et al. 2017), butterflies (Gompert et al. 2015), Darwin’s finches (Chaves et al. 2016), and trees (Lind et al. 2017). BSLMM is a hybrid of LMM and Bayesian variable regression that extends the Lande & Arnold (1983) multiple regression approach in an attempt to address the sparsity of common data sets used in genotype associations, where the number of model parameters (loci) is often much greater than the number of observations (sampled individuals; Zhou et al. 2013; Gompert et al. 2016). Specifically, the model takes into account relatedness among individuals and provides a means to summarize estimates of selection across the genome such as the proportion of phenotypic variation explained (PVE) across genotyped markers by estimating the combined influence of markers with either polygenic (infinitesimal) or measureable (moderate to large) effect, the proportion of PVE explained by genetic loci with measurable effects (PGE), and the number of loci with measurable effects that underlie the trait (for more details see Guan & Stephens 2011; Zhou et al. 2014; Gompert et al. 2016). Additionally, `GEMMA` returns the posterior inclusion probability for each marker providing evidence for association with the phenotype. While the approach remains promising considering its performance in the context of genomic prediction and inference of PVE (e.g., Zhou et al. 2013, Speed & Balding 2014), there has been no attempts, to our knowledge, to assess the approach under various demographic histories, genetic architectures, and sampling designs. A close approximation to this comes from analyses carried out by Gompert et al. (2016), in which `GEMMA` was evaluated for PVE estimation, estimated effects of causative loci, and the estimated number of underlying SNPs based on various author-specified numbers of causal loci, underlying heritability ranges, and numbers of sampled individuals. In short, the authors convey that `GEMMA` is promising, but that there are important limitations to consider (Gompert et al. 2016). However, because the authors simulated architectures by randomly assigning effects to loci from an empirically-derived sequence data set, and while they were thorough in their data exploration, we encourage these results be replicated *in silico* through full modeling of genomic loci across various demographic, LD, sampling, and architecture scenarios to ensure underlying allele frequencies among populations and LD (within and among populations) reflect realistic patterns which may have an effect on model performance. Such additional analyses will also allow for more specific insight into model performance based on *a priori* biological insight available to investigators, allowing more informed decisions when choosing an appropriate genotype-phenotype association method such as BSLMM.

Random Forests (Breiman et al. 2001) is a machine learning algorithm used to identify patterns in highly dimensional data sets to further generate predictive models. Alongside uses outside of evolutionary biology, the Random Forests algorithm has gained popularity in association studies across taxa as well as in trees such as that of genotype-phenotype associations in Sitka spruce (*Picea sitchensis;* Holliday et al. 2012) and genotype-environment associations in white spruce (*P. glauca;* Hornoy et al. 2015). Random Forests is based upon classification (for discrete variables, e.g., soil type) and regression (continuous variables; e.g., temperature or phenotypic measurements) trees (so-called CART models). During its implementation, Random Forests creates these decision trees using two layers of stochasticity: the first layer is used to grow each tree by using a bootstrap sample of observations (environmental or phenotypic) while the second uses a random subset of predictors (marker loci) to create a node which is then split based on the best split of the observations across permutations of predictors using the residual mean square error (see Figure 2 in Hornoy et al. 2015). The observations that were not used as training data to create the model are then used to estimate model accuracy, which can be further used to assess variable importance (Holliday et al. 2012; Hornoy et al. 2015; Forester et al. 2017).

While creating a promising alternative to univariate approaches, until recently the Random Forests algorithm has not been fully explored to assess model performance for use in association studies. Forester et al. (2017) provide a thorough analytical assessment using simulated data to remark on performance for use in genotype-environment association studies (GEA). In their analysis, they used published simulations of multilocus selection (Lotterhos & Whitlock 2014, 2015) of various demographic histories and selection intensities across 100 causative (with 9900 neutral) loci to compare the Random Forests algorithm to the multivariate approaches of constrained ordination (redundancy analysis, RDA, and distance-based RDA, dbRDA - both of which are mechanistically described in Legendre & Legendre 2012, but are multivariate analogs of multiple regression on raw or distance-based data) and to the univariate latent factor mixed model (LFMM). In short, Forester et al. (2017) found that LFMM performed better than Random Forests as a GEA, while constrained ordinations resulted in relatively lower false positive and higher true positive rates across levels of selection than both Random Forests and LFMM. Additionally, the authors found that correction for population structure had little influence on true and false positive rates of ordination methods, but considerably reduced true positive rates of Random Forests. They also note that further testing is needed across various evolutionary scenarios. Even so, constrained ordination provides an effective means by which to detect loci under a range of both strong and weak selection (Forester et al. 2017). While promising under a GEA framework, future analyses may provide evidence that such methods also perform well in genotype-phenotype associations as well. Empirically, it has been used in trees to explore multivariate relationships between phenotypes, genotypes, and environments (e.g., Sork et al. 2016). Additionally, there have been many extensions of the original Random Forests model, such that extensions with purportedly better performance should be assessed alongside other popular association methods in the future.

Once a set of candidate loci have been identified to putatively underlie a phenotype or environment of interest, these loci can be used to further test the hypothesis of polygenic local adaptation. For instance, Berg & Coop (2014) use the significant hits from GWAS data sets to estimate within-population additive genetic values by calculating the frequency-weighted sum of effects across these loci. These values are then compared to a null model of genetic drift that accounts for population structure to test for an excess of variance among populations, ultimately identifying the populations most strongly contributing to this signal. The excess variance statistic (Q_x_) is analogous to *Q*_ST_ and is composed of two quantities - an *F*_ST_-like component describing allele frequency differentiation across populations and a LD-like component describing coordinated and subtle allele frequency shifts across populations. This method thus allows explicit hypothesis tests related to the expected polygenic architecture of local adaptation across populations of trees. It is also noteworthy in that it combines aspects of the genotype-environment-phenotypic spectrum that underlies local adaptation within a single methodological framework (cf. Sork et al. 2013). Prior attempts take a pairwise approach examining each pairwise combination of the genotype-environment-phenotype spectrum (e.g., Eckert et al. 2015). Despite the promising insight from this method, it has not been used widely outside of model organisms. Future applications in trees should consider the number of causal loci identified to be associated with quantitative phenotypes (driven somewhat by the number of loci used in mapping studies), the number of populations needed to increase power, especially in the correlation of genetic values to environmental data, and the ability to reliably estimate genotypic effects.

### At the trail junction – where to next?

While we have outlined methods above that have not yet realized their full potential in describing genetic architecture of complex traits in trees, there are several matters that we, as a field, must keep in mind such that we can continue to progress our understanding in the most efficient manner. Here we believe the path forward lies in three critical areas which we discuss in further detail below: 1) needed data, 2) standardized data reporting, and 3) empirical studies in trees designed to test theoretical expectations of genetic architectures.

### Needed data

While the common garden approach can facilitate understanding of evolutionary processes without specifically identifying underlying loci (Rausher & Delph 2015), identifying features of the genetic architecture will ultimately inform breeding applications important to management, conservation, and industry, and thus requires knowledge about underlying loci. Consequently, we have not yet had sufficient sampling of both marker densities and studies amenable to replication across systems to truly exhaust the use of single-locus approaches, particularly as the sample size of markers, individuals, and populations increase in the near future. Indeed, Hall et al. (2016) estimated that the number of causative loci underlying quantitative traits in trees is likely in the several hundreds, and to capture 50% of the heritable genetic variation using single-locus approaches, population sizes of about 200 will be needed for mapping disease traits, and about 25,000 for traits such as growth. Even so, we recommend that such single-locus associations should not be used as the sole method of architecture description as we carry out future studies unless justified a *priori* based on biological principles, knowledge of the expected architecture, and/or for testing specific hypotheses. While the limits of such methods should be considered, these approaches can be used alongside other lines of evidence to either support or spur further testing of underlying loci (*sensu* Sork et al. 2013). For instance, there is little downside to performing both a single-locus association and a multivariate analysis in the same study, even if some or all of the results for a given technique are excluded to the supplement (e.g., Sork et al. 2016). Further, contextualizing genotype-phenotype and genotype-environment relationships with results that describe local adaptation (e.g., phenotype-environment, *Q*_ST_-*F*_ST_ comparisons) can also stimulate further understanding particularly for data that is made publically available for future synthesis. Specifically, studies which do so within the context of comparisons within and across species (e.g., Yeaman et al. 2016) or environments (Holliday et al. 2016), offer unique circumstances under which to advance our understanding of complex traits in trees (Table 1; Lotterhos & Whitlock 2015; Ćalić et al. 2016; Hoban et al. 2016; Ingvarsson et al. 2016; Mahler et al. 2017).

Isozymes (Adams & Joly 1980), restriction fragment length polymorphisms (Devey et al. 1994), randomly amplified DNA (Grattapaglia & Sederoff 1994), and expressed sequence tag polymorphisms (Temesgen et al. 2001) were among the first used to test evolutionary hypotheses in trees related to genome organization and the mapping of complex traits (discussed in Eckert et al. 2009). Marker technology has progressed considerably since this time (dozens of markers) to include markers capable of more densely sampling tree genomes (up to millions of markers). For example, array-based designs (Silva-Junior et al. 2015) and exome capture (Suren et al. 2016) allow for hundreds to tens of thousands of both genic and intergenic markers (which can be dwarfed by the number of subsequently called SNPs) whereas RADseq (reviewed in Parchman et al. in review) is in the range of tens-to hundreds of thousands of markers (e.g., Parchman et al. 2012) and whole genome sequencing in the range of millions (e.g., Stölting et al. 2015). However, while the continual advent of sequencing technology will likely allow for more SNPs and longer sequences, it is ultimately the concordance between polygenic expectations and analytical methods of marker data that will determine the success of such endeavors. With this in mind, future studies aimed at answering outstanding questions (Table 1) will benefit from a diverse set of markers that represent both functional proteins (genic regions) as well as those which control aspects of their expression or post-transcriptional regulation. If one lesson is to be gained from the recent discussion of the applicability of reduced representation techniques (Lowry et al. 2016, 2017; Catchen et al. 2017; McKinney et al. 2017), it is that genomic resources are paramount to advancement of knowledge, especially when developed with knowledge of patterns of linkage disequilibrium or, if not with this knowledge, with goal of quantifying it. However, RADseq remains one of the most cost-effective approaches available to trees and should thus be assessed in the specific context of tree species, particularly when strengths and limitations are understood and addressed (as reviewed in Parchman et al. forthcoming). No matter the approach used for association, some aspect of the architecture is likely to be missed in trees. For example, RADseq-based markers developed within large genomes are not enriched within genic regions where structural changes to proteins are expected to affect phenotypes, although choice of enzymes can affect the relative proportion of genic regions in tree genomes, as evidenced from *in silico* digestions of reference genomes from *Populus, Eucalyptus, Amborella, Pseudotsuga,* and *Pinus* species (Parchman et al. forthcoming). In contrast, exome based approaches are anchored within coding regions thus excluding putative regulatory elements outside of the exomic regions used to develop probes. Recent marker development approaches, such as RAPTURE (Ali et al. 2016), however, have blurred the lines between RADseq and exome based approaches and, in addition to targeted capture approaches, may offer a promising, cost-effective path forward that explicitly avoids biased assumptions about the importance of exomic versus intergenic loci comprising the architecture of local adaptation.

Beyond dense genetic linkage maps (e.g., Friedline et al. 2015) and reference genomes, which undoubtedly should be among our top priorities, other techniques outside of traditional genomics, such as transcriptomics, have the potential to complement genomic studies in many ways without great need for existing species-specific resources (reviewed in Romero et al. 2012, Strickler et al. 2012; Vialette-Guiraud et al. 2016). For instance, comparative transcriptomic techniques in trees can be used to identify putatively orthologous sets of markers (e.g., Wachowiak et al. 2015; Yeaman et al. 2016) that can be used to describe the evolution of architecture (e.g., shared orthologs versus paralogs across species) or for comparative linkage mapping (Ritland et al. 2011) across systems. Additionally, with the appropriate study design, transcriptomics can be implemented in tree species to describe various aspects of differential expression (Cohen et al. 2010; Carrasco et al. 2017; Cronn et al. 2017), selective constraint (Mähler et al. 2017), prevailing selective forces (Hodgins et al. 2016), mapping of disease resistance (Liu et al. 2016; Liu et al. 2017), and regulatory networks (Zinkgraf et al. 2017). The multilocus paradigm of transcriptomics is amenable to identifying and testing hypotheses of the genetic architecture of complex traits in a network framework (Jansen et al. 2009; Leiserson et al. 2013; Civelek & Lusis 2014; Feltus 2014) and will no doubt provide valuable contributions for tree evolutionary biologists. Other areas amenable to network description such as metabolomics and proteomics would also be a complement (Feltus 2014; Cowen et al. 2017), particularly if genetic studies contextualize results with findings from such approaches and vice versa. Ultimately the goal is to use *a priori* knowledge synthesized across past studies, techniques, and perspectives to guide further hypotheses about underlying architecture, as exemplified by Mizrachi et al. (2017) and Lotterhos et al. (2017). Finally, high-throughput phenotyping as well as environmental measures at fine spatial scales below square-kilometers will also facilitate and advance our understanding of complex traits in trees (Sork et al. 2013; Rellstab et al. 2015; Leempoel et al. 2017), particularly when measured phenotypes well represent those experiencing selection pressure, and environmental measures well represent the multivariate environment imposing selection (Lotterhos et al. 2017).

### Standardized data reporting

As we continue to accrue genotype-phenotype, genotype-environment, and phenotype-environment relationships within and across tree species, authors should consider how their results can most effectively be used in further studies and syntheses, both for the purpose of validation or comparison as well as novel insights yet to be seen. Here we outline a few suggestions that can be broken down into reporting within manuscripts and metadata. For instance, in our survey of common garden studies used to estimate *h*^2^ and *Q*_ST_, in many cases the exact design of the study could not be replicated with the information from the manuscript alone. While an abbreviated design may be suitable for the main text, authors can provide much more detail in supplemental materials that can facilitate replication and comparison across studies (e.g., total individuals per garden, family, or block – as opposed to averages or ranges), which will ultimately facilitate syntheses regarding future directions. Further, future studies would benefit from estimating relatedness using marker data which will ultimately improve the precision of *h*^2^, *Q*_ST_, and missing heritability estimates (de Villemereuil et al. 2016) including those estimates made in the field (Castellanos et al. 2015). For cases in which estimating relatedness from markers is not appropriate or feasible, the field would benefit by authors exploring a range of underlying sibships (e.g., Eckert et al. 2015), which are often assumed to be half-sib relationships. While some studies in our survey assumed a mixed sibship relationship for open-pollinated sources, ultimately such assumptions without data exploration will affect the outcome or conclusions for any given study. A recently released R package by Gilbert and Whitlock (2014) allows for such an exploration of effects of mixed sibships on inference of *Q*_ST_ and its magnitude relative to *F*_ST_. Inclusion of such exploration, even in the supplement, will help contextualize such studies as they are published. For studies estimating causality for genotype to phenotype, it would be worthwhile to include the regression coefficients or other estimates of effect size (e.g., odds ratios) in addition to PVE (*r*^2^). Importantly, the units of the effect size must be explicitly reported (e.g., Julian days versus phenotypic standard deviations), with the standard deviation also reported. For all association studies, supplemental tab- or comma-delimited text files (outside of a word processing document) easily analyzed with programming languages would also facilitate synthesis (even if providing redundant information from the main text), particularly if such files are well described with a README file and contained data regarding marker position, putative orthogroups, hits to reference genomes, effect size, PVE, genotypes by individual identifiers, individual population assignments, and if the sequence or marker was significantly associated to phenotype or environment. Such an operating procedure may work well in the short term, however in the long term such information will need to be easily accessible from one or a central hub of repositories.

Data standardization, the inclusion of meta-information, and compilation of these data specific to trees into a database with common terminology will be crucial to future inquiries with the purpose of synthesizing evidence for underlying architectures across species and environmental systems (e.g., as for human GWAS data: https://www.ebi.ac.uk/gwas/). If the data generated by tree biologists is disparate and housed across databases and journal supplements this impedes synthesis first by forcing scientists to collate information across sources, which may be further impeded by data redundancies or inconsistencies in data format and utilized nomenclature (Wegrzyn et al. 2012). While many journals have required submission of sequence data to repositories such as NCBI, such databases are lacking with regard to information pertaining to phenotypic, environmental, and geographic information upon which much of the foundation of our field is built. Submissions to Dryad somewhat overcome this, but there is no standardization within the community for content for such submissions and important information may be lacking. Currently, this information is often appended in supplemental files that cannot be readily accessed, compared, or queried in an efficient manner. Hierarchical ontologies can be used to ease this burden. Gene Ontology is likely the most recognizable to evolutionary biologists, but there also exist Plant Ontologies for organismal structure and developmental stages, Environmental Ontologies for habitat categorization, and Phenotypic, Attribute, and Trait Ontologies for the annotation of phenotypes. Such ontologies not only standardize nomenclature, but also assist in database queries. The utilization of such databases will no doubt encourage comparative studies and syntheses, as infrastructure and data accessibility are essential to the comparative approach (Neale et al. 2013; Ingvarsson et al. 2016; Plomion et al. 2016). Luckily, such a database exists for the broader tree genetics community. The open-source genomics and phenomics database, called TreeGenes (treegenesdb.org), is part of a central hub of repositories, including the Hardwood Genomics Project (hardwoodgenomics.org) and the Genome Database for Roseaceae (rosaceae.org), that communicate and integrate data from each other. Unlike many other repositories for tree genomic data, TreeGenes is not project or institution specific. The data and metadata for roughly 1700 species housed on TreeGenes can be accessed, queried, and visualized through DiversiTree, a web-based, desktop-style interface (Wegrzyn et al. 2008). DiversiTree connects to the geographical interface CartograTree (Vasquez-Gross et al. 2013) to encourage comparative synthesis by providing technology to filter and visualize geo-referenced biotic and abiotic data housed on TreeGenes. As promising as such database hubs are, they are only as useful as the data that is deposited to them. While TreeGenes will regularly import and enhance data from public repositories (through e.g., sequence alignment to published genomes, or data from Genbank, Phytozome, PLAZA, etc), often pertinent metadata necessary for comparative synthesis is lacking (Wegrzn et al. 2008, 2012). Indeed, from our survey of published GPA since the release of the database in 2008, less than 13% (6/48) of the studies submitted their data directly to TreeGenes. To better prepare for future synthesis, we advocate that authors submit their data to the TreeGenes database and that reviewers and editors enforce this habit, as currently implemented for linkage maps published in *Tree Genetics & Genomes.* Consolidated, open-source resources will be crucial to the advancement of this field (Neale et al. 2013), and will no doubt spur knowledge that would not have been recognized otherwise. Prime examples of advancement to knowledge because of these types of resources and community-wide efforts come from the human GWAS literature where such resources provide crucial information necessary to study polygenic adaptation (e.g., Berg & Coop 2014).

### Empirical tests of theory

In combination with the development of truly genome-wide public resources, there is need to use these resources to validate and better characterize foundational ideas and assumptions in the theory of polygenic adaptation relative to the life history strategies of tree species. For example, Gagnaire & Gaggiotti (2016) highlight that the degree of polygenicity can be tested as a function of the number of GWAS hits relative to the length of contigs or chromosomes containing these markers. Simple models of polygenicity predict that there should be a positive correlation between these quantities. Thus, rather than assuming some functional form of a polygenic architecture (i.e., an approximate infinitesimal model) during analysis, researchers can strive to characterize, or at least exclude some forms of, the underlying genetic architecture prior to interpretation. In a related fashion, publically available data sets would spur comparisons across species and study systems to test hypotheses about polygenic architectures (e.g., the modularity of genetic architectures as in Lotterhos et al. 2017, or perhaps genomic organization or effect size distribution) due to the relative timing of selection, degree of environmental contrast (e.g., diversifying selection and changes to the strength of negative selection), selection strength, and level of gene flow across diverging lineages. As an example, much of the theory of polygenic adaptation requires assumptions about simplistic demographics (where violations have consequences for standing levels of non-neutral diversity, e.g., Wang et al. 2017) and the equilibration among co-acting evolutionary forces over a large number of generations (Brandvain & Wright 2016). Indeed, differing architectures are expected as a function of the timing for the onset of selection (Le Corre & Kremer 2003; Kremer & Le Corre 2012), with subtle allele frequency shifts across populations dominating architectures near the onset of selection and larger allele frequency shifts much later in time. While there is need for empirical validation of this theory, there is also a need to characterize the prevalence of its predicted patterns across differing clades of tree species. In other words, researchers could imagine testing the theory itself in natural populations (e.g., as begun by Le Corre & Kremer 2012) or assuming its validity and characterizing the circumstances under which to expect large shifts in allele frequencies across tree species with differing life history strategies. Little of any of this (Table 1), however, will be possible without development of needed data and its deposition into publically available, standardized databases.

## Concluding Remarks

The path forward provides a means by which we can most efficiently describe the underlying genetic architectures of traits important to management, conservation, and industry, which can ultimately be used to expedite breeding projects (Box 1). The past evolutionary history will have a profound effect on the underlying genetic architecture of such traits, and thus strengths and weakness of the data and methods used to uncover such architecture should be specifically addressed in the future, particularly in how utilized methods perform across various demographic and architecture scenarios. Insights gained from empirically testing theory will also contribute to the advancement of this field and will ultimately quantify the variation in architecture across environments and species and inform effective management. Importantly, the success of future genotype-phenotype efforts should not be predicated on past studies using single-locus approaches and small numbers of markers, and instead on overcoming such shortcomings by applying theoretical expectations to empirical inquiry. Even so, until sequencing technologies allow for cost-effective whole genome sequencing of individual trees, most genotype-phenotype studies (GS included) will be carried out via reduced representation techniques (i.e., a subset of all sites within the genome). Therefore, it is essential that processed data be uploaded to a repository that, in addition to raw sequences, includes genotypic, environmental, and spatial data, facilitates user-friendly queries, and allows for future meta-analysis. The future is bright, but we are not yet out of the woods. As such, efficient advancement in this field relies on community efforts, standardized reporting, centralized open-access databases, and continual input and review within the community’s research.

## Acknowledgements

The authors would like to thank S. González-Martínez for inviting this review, Chris Friedline for stimulating conversations when developing content, and Justin Bagley, Jill Wegrzyn, and two anonymous reviewers for providing helpful comments to earlier versions of this manuscript. Brandon Lind is supported through a Dissertation Fellowship provided by the Graduate School of Virginia Commonwealth University. Andrew Eckert is supported through the National Science Foundation (EF-1442486) and the United States Department of Agriculture (USDA 2016-6701324469).

## Author Contributions

BML and AJE conceived the review, with contributions from MM, CEB, and TMF. BML, MM, CEB, and TMF contributed to the literature search and survey which was analyzed by BML. CEB summarized *Q*_ST_ and *F*_ST_ comparisons. BML wrote the manuscript with contributions from MM and AJE. All authors contributed to the editing of the manuscript.

## Tables

***Table 1.*** Where to next? The Path Forward identifies meaningful ways in which we can progress our understanding of the architecture underlying complex traits in trees. Here we outline some questions that can be used to guide future inquiry as the number of markers and sequence length increase, and annotation becomes more precise and specific to tree biology.

1. Composition and evolution of genetic architectures in trees

a. How prevalent are non-additive contributions to underlying genetic architectures in trees? Are there patterns across similar phenotypes or regulatory networks? Is there evidence that such non-additive effects have either constrained or facilitated local adaptation?
b. Are adaptive loci most prevalent in areas of low recombination or repetitive sequences (e.g., retrotransposons, clustered gene families)? Do loci of similar effect sizes, expression profiles, or pleiotropic effect (Lotterhos et al. 2017) experience elevated LD within the genome? Should genome size influence our expectations for underlying architectures (Mei et al. 2017)?
c. At what frequency does local adaptation result in fitness tradeoffs across environments (Tiffin & Ross-Ibarra 2014; Wadgymar et al. 2017)? And does this interact with demographic history in trees?
d. Does pleiotropy play a predictable role in underlying tree genetic architectures (Lotterhos et al. 2017)?
e. Which aspects of genetic architectures in trees are likely to exhibit deleterious variation? And how much of this signal are we capturing in genotype-phenotype applications?
2. Inter- and intraspecific variation of genetic architectures in trees

a. Which aspects of the genetic architecture should we expect to vary across populations or environments?
b. Under what conditions in trees are we likely to observe genomic reorganization across species or ecotypes (e.g., physical linkage or dispersion)? Will reference genomes be suitable to assess this question across species or diverged populations, or can long-read sequencing technologies (reviewed in Jiao & Schneeberger 2017) offer appropriate resources?
c. What is the degree of convergent and parallel adaptation within polygenic architectures across tree populations and species?
d. At what level of the genetic architecture do we see patterns of convergence, parallelism, and divergence? Within core hubs, or perhaps within aspects of the periphery? What does the comparison of the topologies from such architectures tell us about influential evolutionary processes?
e. How often are architectures influenced by variation in expression levels rather than structural variation in proteins? Do architectures differ in predictable ways with the prevalence of one or the other? How can we utilize knowledge synthesized across past approaches to spur understanding of underlying genetic architectures in trees (Mizrachi et al. 2017)?

## Box 1 A step in the right direction: Synergism between GWAS and Genomic Selection

Early simulations showcased the promise of predicting breeding values from marker data to accelerate domestication and breeding of plants and animals (Meuwissen et al. 2001; Bernardo & Yu 2007; Heffner et al. 2009; Zhong et al. 2009), and particularly under the framework of genomic selection (GS) in trees (Wong & Bernardo 2008; Grattapaglia & Resende 2011; Iwata et al. 2011; defined and reviewed by Grattapaglia 2017). Much of the early exploration into the applicability of GS in trees discounted the utility of marker-assisted selection (MAS) because of the small estimated effects for the few loci significantly associated via single-locus approaches at the time, as well as having concerns related to replication because of the identification of markers across limited parental (genetic) backgrounds (Grattapaglia & Resende 2011; Iwata et al. 2011; Resende et al. 2012a, 2012b). Based on these arguments and results from simulations, genomic selection was identified as a more promising endeavor than MAS, particularly if the breeding cycle can be reduced via efforts such as grafting (Grattapaglia & Resende 2011) or somatic embryogenesis (Resende et al. 2012a).

While GS techniques often can explain a considerable proportion of narrow sense heritability, current implementation of GS in trees is often on par with, or marginally better than, traditional phenotypic selection when evaluating potential within the same generation and environment (see Table 9.1 in Grattapaglia 2017). Further, the predictive accuracy of various models are a function of underlying architecture (Resende et al. 2012c; Grattapaglia 2017). As pointed out by Grattapaglia (2017), current marker densities have produced satisfactory results due to the capture of relatedness between training and validation populations. Here, this success is likely due to the ability of markers to reasonably represent large haplotype blocks (and thus cumulative action of causative effects) due to the high level of relatedness between training and validation populations. Even so, Grattapaglia (2017) recommends higher marker densities so that markers also capture true marker-QTL LD and thus sustain long-term accuracies across generations and environments. We also believe GWAS applications (in the broadest sense) in trees will also see improvements through increased marker densities, the results of which can then be used to further test specific hypotheses regarding underlying architectures and to increase predictive accuracies of GS as well. Incorporating markers that putatively underlie the trait of interest into model prediction may spur opportunities that do not require high degrees of relatedness between training and validation populations, perhaps to the extent of incorporating material from outbred stands using predictive approaches (*sensu* Bérénos et al. 2014; Bontemps 2016) and heritability validation (*sensu* Castellanos et al. 2015) in the field.

In the end, the realized progress of our understanding regarding the genomics of complex traits in trees will therefore be enhanced by the deposition of data from both GS and GWAS (as well as other ‘omics’) approaches into a centralized open-access database hub such as TreeGenes (treegenesdb.org). Future meta-analyses can then synthesize past inquiry to summarize our current understanding of underlying genetic architectures, ultimately incorporating this knowledge towards future applications in industry and conservation (see The Path Forward; Table 1).

## Supplemental Information

### Supplemental Figures

**Figure S1.**
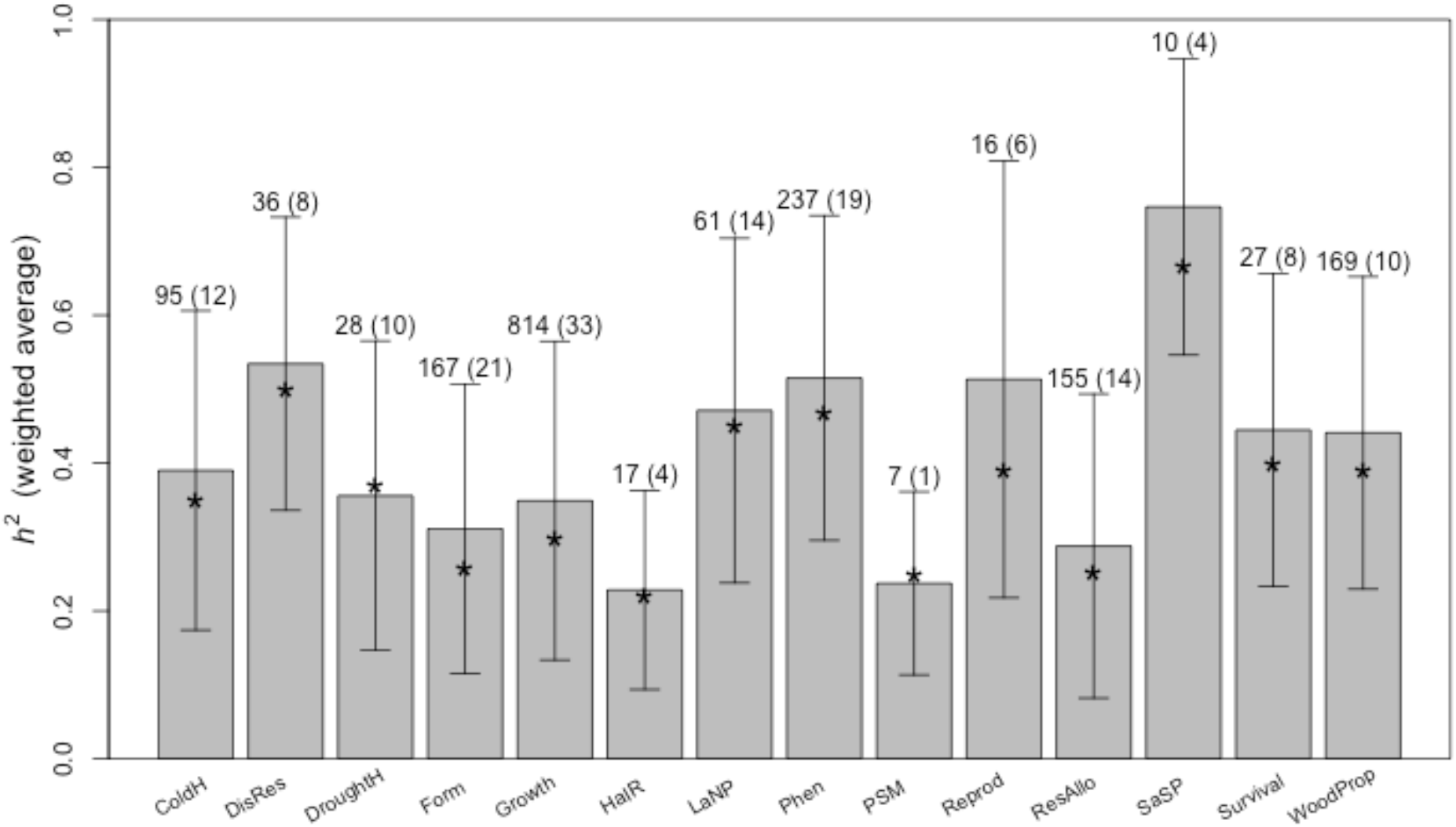
Averages of narrow sense heritability calculated by weighting the number of families used in each estimate of heritability. Error bars represent the standard deviation of the weighted averages. Note that genetic variances in juvenile traits may be inflated due to instances of maternal effects, which we did not control for in our literature survey. Abbreviations as in Figure 1 of the main text.

**Figure S1.**
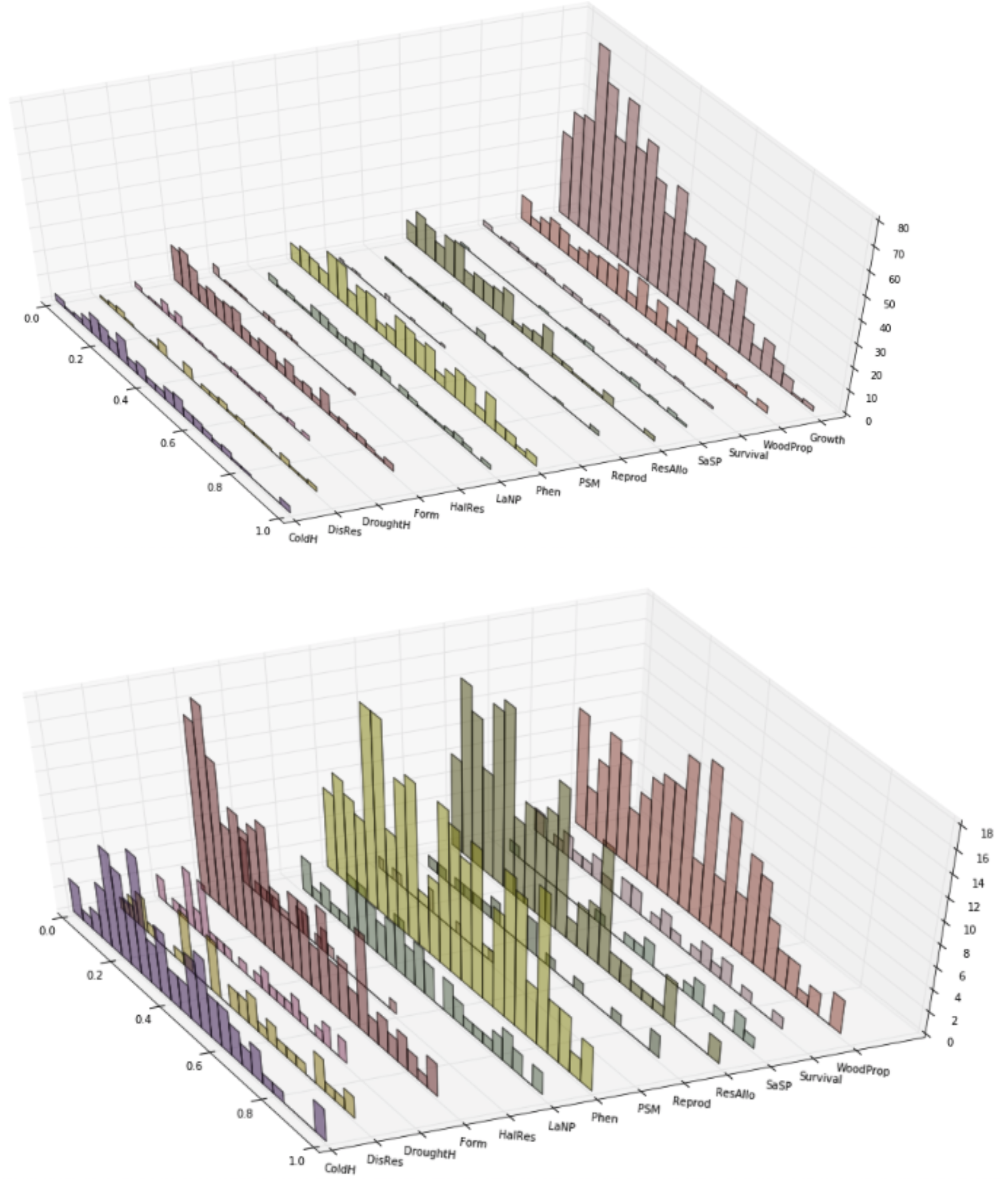
Distributions of unweighted narrow sense heritability with (A) and without (B) inclusion of the Growth distribution. Trait abbreviations as in Figure 1 of the main text.

**Figure S2.**
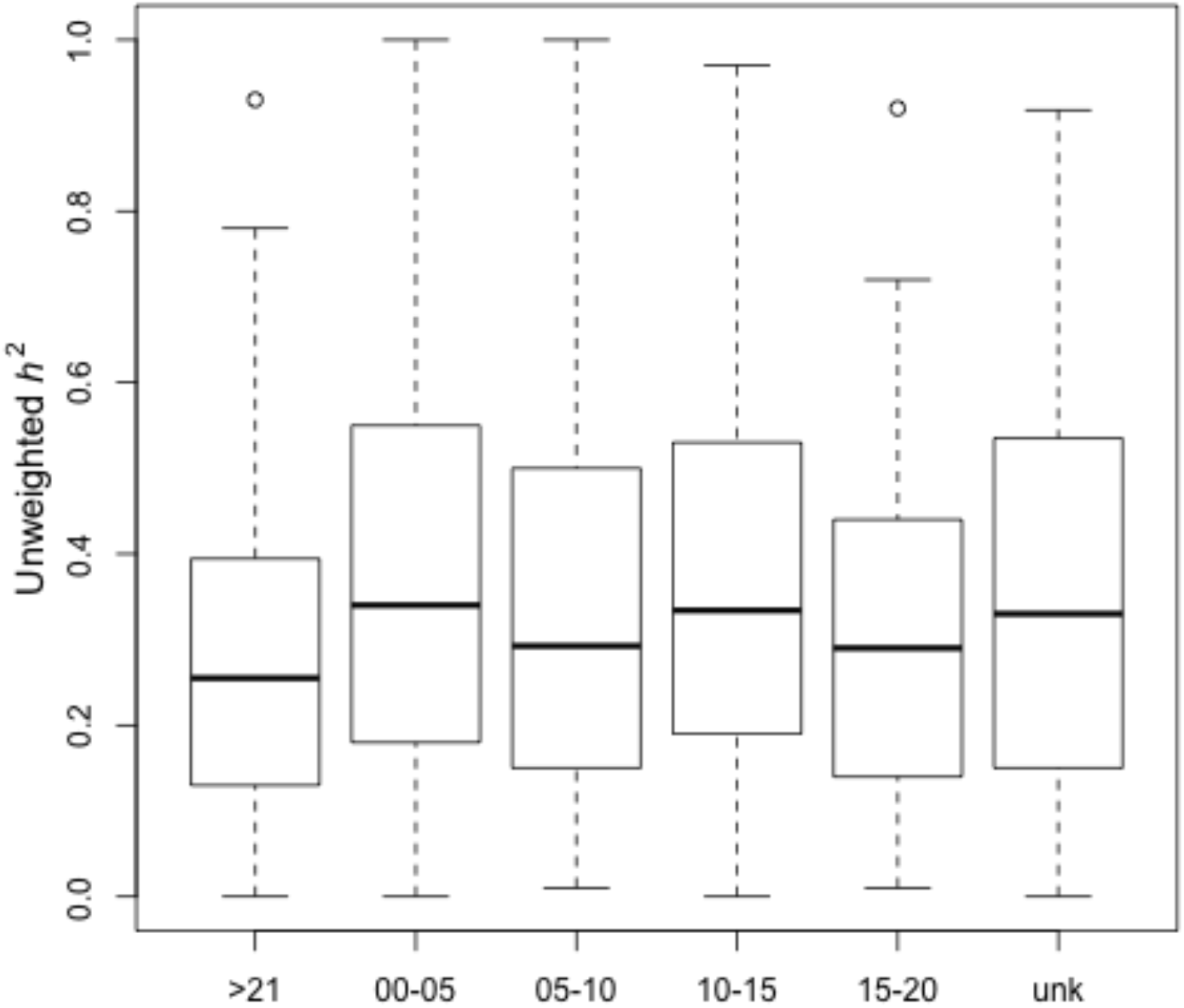
Unweighted narrow sense heritability distributions by age (years). Unk = unknown age (i.e., not specified by article).

**Figure S3.**
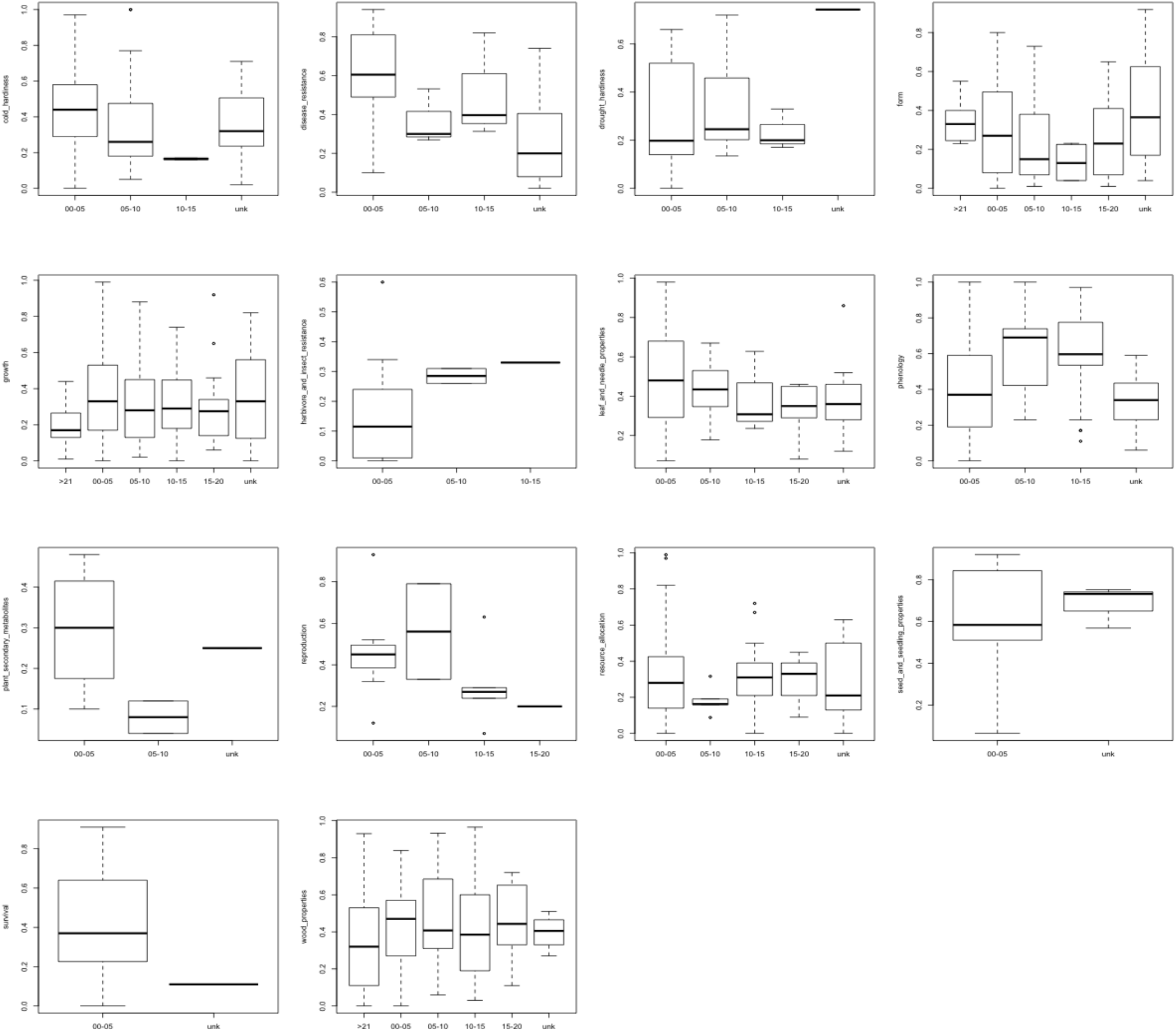
Unweighted narrow sense heritability distributions by age (years) and by trait category. Unk = unknown (i.e., not specified by article)

**Figure S4.**
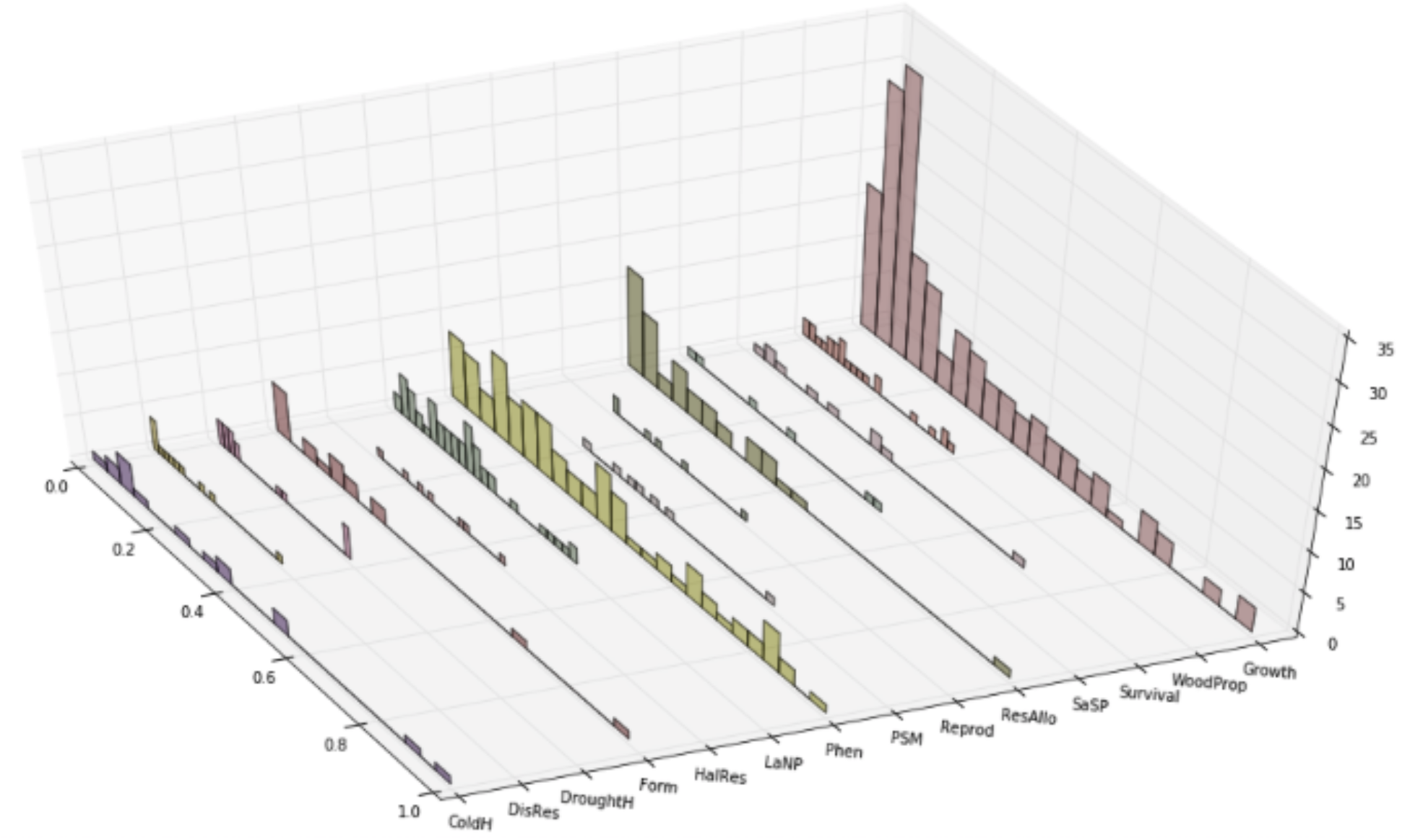
Distributions of unweighted QST estimates from literature survey. Abbreviations as in Figure 1 of the main text.

### Supplemental Tables

**Table 2.**
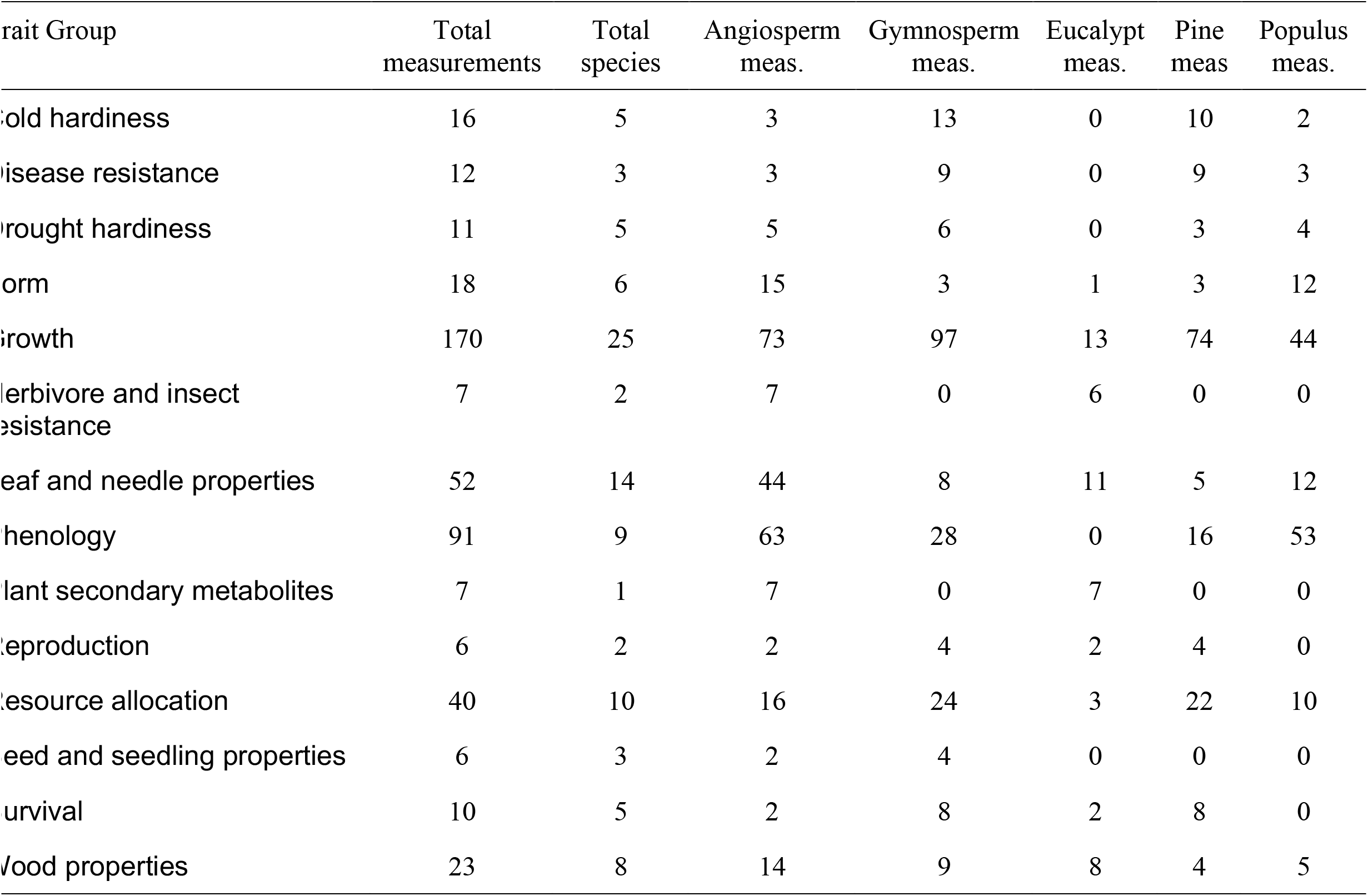
Summary of total and per-species measurements used in literature review of differentiation of quantitative genetic variation (*Q*_ST_). The Total Species column is the number of unique species in our survey, whereas the remaining columns provide the total number of measurements per category.

**Table 3.**
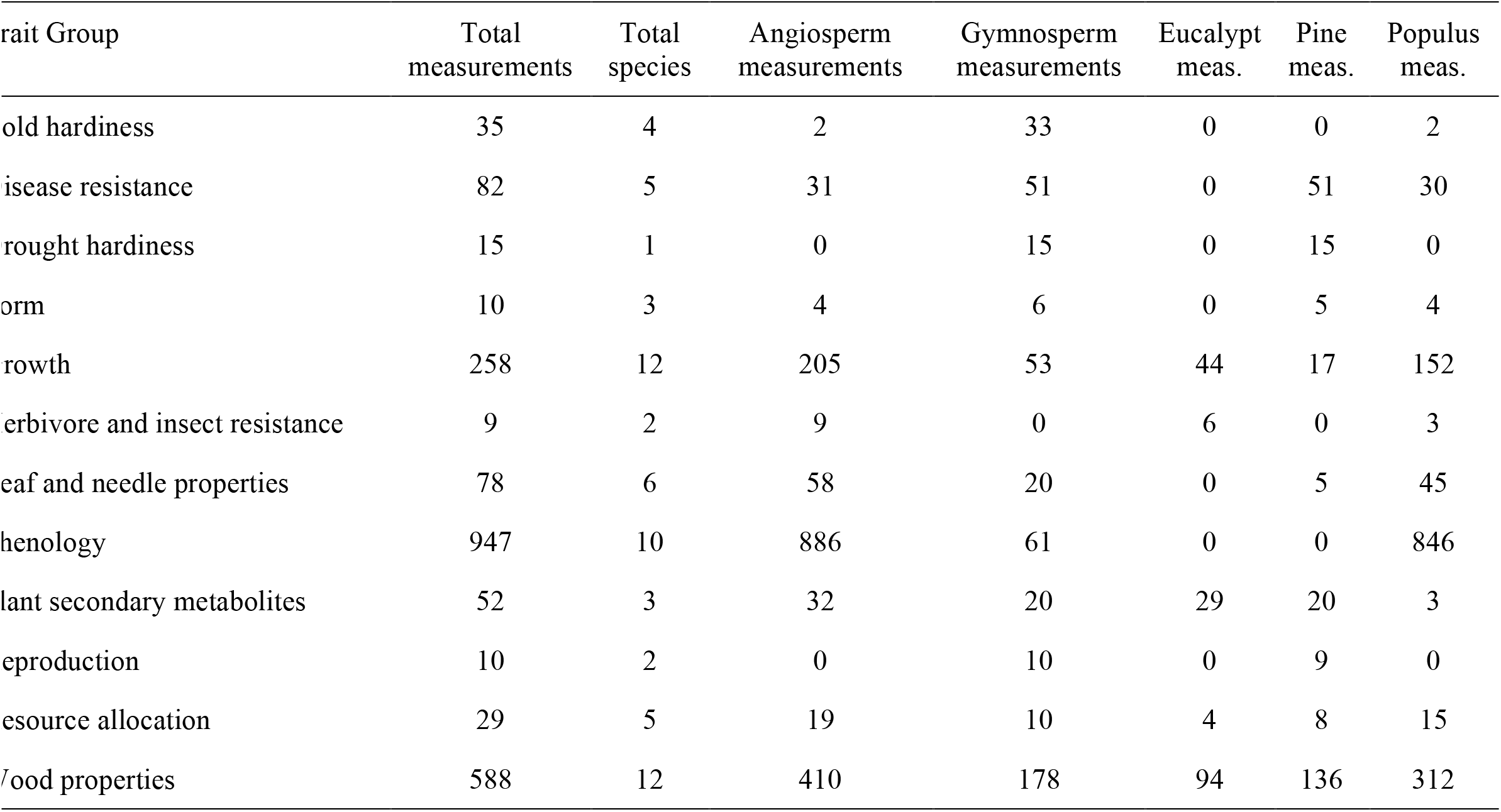
Summary of total and per-species measurements used in literature review of percent phenotypic variance explained by associated markers (*r*^2^). The Total Species column is the number of unique species in our survey, whereas the remaining columns provide the total number of measurements per category.

## BOXES

### Supplemental Box S1

#### Basic Concepts from Quantitative Genetics

We follow the traditional decomposition of phenotypes into genetic and environmental components, which forms the basis of quantitative genetics (Fisher 1918, Lynch & Walsh 1998, Charlesworth & Charlesworth 2010, reviewed by Hill 2010). The phenotype of an individual (P) can be decomposed into effects from its genotype (G), its environment (E), and the interaction between its genotype and environment (GxE). Typically, this is thought of as deviations from the population mean, with each causative locus having two alleles. Using this framework, phenotypic variance (σ^2^_P_) can be decomposed into genotypic variance (σ^2^_G_), environmental variance (σ^2^_E_) and the variance due to the interaction between genotypes and environments (σ^2^_GxE_):

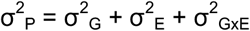

For a single locus, σ^2^_G_ can be decomposed into variances arising from additive (σ^2^_A_) and dominance (σ^2^_D_) effects (Fig. 1). For multiple loci, σ^2^_G_ can be decomposed into variances arising from additive, dominance, and epistatic (σ^2^_I_) effects, with the total additive effect across loci being the summation of the effects at each of the causative loci. Dominance and epistatic effects are jointly termed non-additive effects. Thus, the previous equation can be expanded to the following:

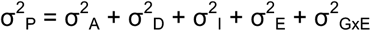

The decomposition of σ^2^_G_ into different types of effects provides a way of estimating narrow-sense heritability (*h*^2^), which is defined as the ratio of additive genetic variance (σ^2^_A_) to total phenotypic variance (σ^2^_P_). For tree populations, this is often accomplished through variance partitioning techniques (Namkoong 1979) using half-sib designs in common gardens (White et al. 2007) or using molecular markers to estimate relatedness in the field (cf. Ritland & Ritland 1996). In the case of half-sib designs, if the assumptions of free recombination and little epistasis among causative loci, random mating, and lack of environmental covariance among sibs are satisfied, σ^2^_A_ is given by (Lynch & Walsh 1998):

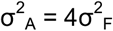

where σ^2^_F_ is the variance due to family (e.g., as extracted from a linear mixed model). Hence, for a half-sib design, *h*^2^ = 4σ^2^_F_/σ^2^_P_. Other sibling designs are possible, with the 4 in the previous equation replaced by 1/*r*, where *r* is the coefficient of relationship (e.g., Whitlock & Gilbert 2012). Clonal and controlled mating designs are also often used for estimation of heritability, often broad-sense heritability (Namkoong 1979; White et al. 2007). When families are nested into populations, and an estimate of the among population variance component is made, these are the components also used to estimate *Q*_ST_ (Spize 1993; Prout & Barker 1993). When compared against estimates of F_ST_ using a similar variance decomposition procedure (e.g., Yang 1998) and a method suitable to account for the substantial variance associated with these components (e.g., Whitlock & Guillaume 2009) conclusions about local adaptation can be reached.

Heritability estimates are population, environment, and time specific, as evidenced by the relationship between σ^2^_A_ and allele frequencies within populations (Lynch & Walsh 1998; e.g. Berg & Coop 2014):

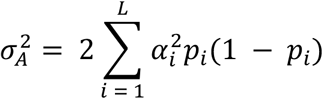

where the summation is over the number of causative loci (*L*), *α* is the average effect of allelic substitution at each locus (Fig. I), and *p_i_* is the frequency of one of the alleles at each of the causative loci. Thus, any evolutionary force altering *p* at some or all of the causative loci will change σ^2^_A_ (cf. Box 3.7 in Charlesworth & Charlesworth 2010). Heritability is also uninformative about the underlying architecture itself, as are the relative magnitudes of the different variance components themselves (Huang & Mackay 2016), and can often be misleading about evolutionary potential (Hansen et al. 2011).

### Supplemental Box S2

#### Brief introduction to methods for single-locus genetic association analysis

Detecting associations between genetic markers and complex trait variation relies on fitting and evaluating linear models, typically of the form:

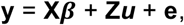

where **y** is a vector of observed or inferred phenotypic values, ***β*** and **u** are vectors of random and fixed effects, respectively, **X** and **Z** are design matrices associated with ***β*** and **u**, and **e** is a vector of residuals (Yu et al. 2005). In the simplest model, the phenotype (**y**) is modeled as a function of genetic effects at a single locus, represented by marker genotypes for the samples comprising values in **y**, and covariates describing relatedness among sampled trees and the structure among populations from which those trees were sampled. Genetic effects are encoded based on *a priori* assumptions about the underlying architecture of the phenotypic trait under consideration, with the most frequent encoding being that for additive effects (e.g., counts of a reference allele) considered as either fixed or random effects (Goddard et al. 2009). Phenotypic values are often estimates derived through analysis of materials established within common gardens, either from clones or sibships, from which estimates of the genetic values of unmeasured trees (e.g., maternal trees for which markers have been genotyped) are made using the theory of Best Linear Unbiased Predictors (BLUPs; Henderson 1975; Searle et al. 1992; Piepho et al. 2008). Inclusion of only fixed effects results in a General Linear Model (GLM), whereas a mixture of fixed and random effects results in a Mixed Linear Model (MLM or LMM). The use of covariates is necessary to avoid identification of false positive associations arising from the confounding between neutral genetic and phenotypic variation due to demographic history and the analysis of relatives (Devlin & Roeder 1999; Yu et al. 2005; Price et al. 2006).

Models as described above are typically fitted and evaluated using restricted maximum likelihood (REML, Patterson & Thompson 1971), although Bayesian methods are available and have the advantage of specifying *a priori* assumptions more clearly, remove the distinction between fixed and random effects, and are more applicable to testing biologically realistic models (Stephens & Balding 2009). Output from these models include estimates of effect sizes for markers (e.g. *r*^2^, coefficients for random effects, genotypic trait means) and, when used in a frequentist framework, probability values (p-values) of observing test statistics under a null model. Bayesian methods, in contrast, provide strength of evidence measures such as Bayes Factors for the association of each marker to the phenotype of interest. The ability to discover and correctly quantify effect sizes of true positives (i.e. causative markers or indirect associations resulting from linkage to causative markers) is dependent upon experimental design, including design of genotyping assays, and sample sizes (Long & Langley 1999; Zöllner and Pritchard 2007; Spencer et al. 2009), as well as genome-wide patterns of linkage disequilibrium relative to the density of markers in the genome, the genetic distance between the indirectly associated maker and the causative locus, and the true underlying genetic architecture of the phenotypic trait under consideration (Platt et al. 2010; Prichard et al. 2010; Caballero et al. 2015).

One model is typically fitted and evaluated per marker-phenotypic trait combination (but see e.g. Wegrzyn et al. 2010 for haplotype analysis). Even without the issue of confounding described above, this increases the likelihood of false positives arising solely from performing many statistical tests. A variety of methods exist to deal with multiple testing, with the most popular methods being those based on the false discovery rate (Storey & Tibshirani 2003) and permutation (Hirschhorn & Daly 2005). Additional methods exist for situations where the multiple tests are not independent from one another (e.g. linkage disequilibrium among markers, see Johnson et al. 2010) or when permutation analysis is problematic (Joo et al. 2016).

## References

Adams WT, Joly RJ (1980) Linkage relationships among twelve allozyme loci in loblolly pine. J Heredity 71:199–202

Alberto FJ, Aitken SN, Alía R (2013) Potential for evolutionary responses to climate change-evidence from tree populations. Global Change Biology 19:1645–1661. doi: https://doi.org/10.1111/gcb.12181

Ali, O. A., O’Rourke, S. M., Amish, S. J., Meek, M. H., Luikart, G., Jeffres, C., & Miller, M. R. (2016). RAD capture (Rapture): flexible and efficient sequence-based genotyping. Genetics, 202(2), 389–400. doi: https://doi.org/10.1534/genetics.115.183665

Álvarez-Castro JM, Carlborg O. A unified model for functional and statistical epistasis and its application in quantitative trait loci analysis. Genetics. 2007; 176: 1151–1167. doi: https://doi.org/10.1534/genetics.106.067348

Anderson JT, LEE C-R, Rushworth CA, et al. (2012) Genetic trade-offs and conditional neutrality contribute to local adaptation. Molecular Ecology 22:699–708. doi: https://doi.org/10.1111/j.1365-294X.2012.05522.x

Arnold SJ (1992) Constraints on phenotypic evolution. The American Naturalist 140:S85–S107. doi: 1 https://doi.org/0.1086/285398

Ashander J, Chevin L-M, Baskett ML (2016) Predicting evolutionary rescue via evolving plasticity in stochastic environments. Proceedings of the Royal Society B: Biological Sciences 283:20161690–10. doi: https://doi.org/10.1098/rspb.2016.1690

Ávila V, Pérez-Figueroa A, Caballero A, et al. (2014) The action of stabilizing selection, mutation, and drift on epistatic quantitative traits. Evolution 68:1974–1987. doi: https://doi.org/10.1111/evo.12413

Bailey SF, Bataillon T (2016) Can the experimental evolution programme help us elucidate the genetic basis of adaptation in nature? Molecular Ecology 25:203–218. doi: https://doi.org/10.1111/mec.13378

Barrett R, Schluter D (2008) Adaptation from standing genetic variation. Trends in Ecology & Evolution 23:38–44. doi: https://doi.org/10.1016/j.tree.2007.09.008

Barton NH (2017) How does epistasis influence the response to selection? Heredity 118:96–109. doi: https://doi.org/10.1038/hdy.2016.109

Barton NH (1990) Pleiotropic models of quantitative variation. Genetics 124:773–782.

Barton NH (1999) Clines in polygenic traits. Genetical Research 74:223–236. doi: https://doi.org/10.1017/S001667239900422X

Barton NH, Etheridge AM, Véber A (2016) The infinitesimal model. bioRxiv 1–54. doi: https://doi.org/10.1101/039768

Beavis W. 1994. The power and deceit of QTL experiments: lessons from comparative QTL studies. Proceedings of the forty-ninth annual corn and sorghum industry research conference. Chicago, IL, USA: American Seed Trade Association, 250–266.

Bérénos et al. (2014) Estimating quantitative genetic parameters in wild populations: a comparison of pedigree and genomic approaches. Molecular Ecology 23: 3434–3451. https://doi.org/10.1111/mec.12827

Berg JJ, Coop G (2014) A Population Genetic Signal of Polygenic Adaptation. PLoS Genetics 10:e1004412. doi: https://doi.org/10.1371/journal.pgen.1004412

Bernardo R, Yu JM. 2007. Prospects for genomewide selection for quantitative traits in maize. Crop Science 47: 1082–1090.

Berry AJ, Ajioka JW, Kreitman M (1991) Lack of polymorphism on the Drosophila fourth chromosome resulting from selection. Genetics 129:1111–1117.

Bessega C, Pometti C, Ewens M et al. (2015) Evidences of local adaptation in quantitative traits in *Prosopis alba* (Leguminosae). Genetica 143: 31. doi:10.1007/s10709-014-9810-5

Blanquart F, Kaltz O, Nuismer SL, Gandon S (2013) A practical guide to measuring local adaptation. Ecology Letters 16:1195–1205. doi: 10.1111/ele.12150

Bontemps et al. (2016) In situ marker-based assessment of leaf trait evolutionary potential in a marginal European beech population. Journal of Evolutionary Biology 29: 514–527.

Boshier D, Broadhurst L, Cornelius J, et al (2015) Is local best? Examining the evidence for local adaptation in trees and its scale. Environmental Evidence 4:1–10. doi: 10.1186/s13750-015-0046-3

Bower AD, Aitken SN (2008) Ecological genetics and seed transfer guidelines for Pinus albicaulis (Pinaceae). American Journal of Botany 95:66–76.

Boyle EA, Li YI, Pritchard JK (2017) An Expanded View of Complex Traits: From Polygenic to Omnigenic. Cell 169:1177–1186. doi: 10.1016/j.cell.2017.05.038

Brandvain Y, Wright SI (2016) The Limits of Natural Selection in a Non-equilibrium World. Trends in Genetics 32:201–210. doi: 10.1016/j.tig.2016.01.004

Breiman, L. (2001). Random Forests. Machine Learning, 45(1), 5–32. http://doi.org/10.1023/A:1010933404324

Budde KB, Heuertz M, Hernandez-Serrano A et al. (2014) In situ genetic association for serotiny, a fire-related trait, in Mediterranean maritime pine *(Pinus pinaster)*. New Phytologist 201:230–241.

Bulmer MG (1980) The mathematical theory of quantitative genetics. Genetical research 19: 1725.

Bürger R (1999) Evolution of Genetic Variability and the Advantage of Sex and Recombination in Changing Environments. Genetics 153:1055–1069.

Bürger R, Akerman A (2011) The effects of linkage and gene flow on local adaptation: A two-locus continent-island model. Theoretical Population Biology 80:272–288. doi: 10.1016/j.tpb.2011.07.002

Bürger R, Lynch M (1995) Evolution and extinction in a changing environment: a quantitative-genetic analysis. Evolution 49:151–163. doi: 10.2307/2410301

Burghardt LT, Young ND, Tiffin P (2017) A Guide to Genome-Wide Association Mapping in Plants. Current Protocols in Plant Biology. doi: 10.1002/cppb.20041

Caballero A, Tenesa A, Keightley PD (2015) The nature of genetic variation for complex traits revealed by GWAS and regional heritability mapping analyses. Genetics 201:1601–1613.

Ćalić I, Bussotti F, Martínez-García PJ, Neale DB (2015) Recent landscape genomics studies in forest trees. Tree Genetics & Genomes 1–7. doi: 10.1007/s11295-015-0960-0

Carlborg Ö, Haley CS (2004) Epistasis: too often neglected in complex trait studies? Nature Reviews Genetics 5:618–625. doi: 10.1038/nrg1407

Carrasco A, Wegrzyn JL, Durán R, et al (2017) Expression profiling in Pinus radiata infected with Fusarium circinatum. Tree Genetics & Genomes 13:1665. doi: 10.1007/s11295-017-1125-0

Carter AJR, Hermisson J, Hansen TF (2005) The role of epistatic gene interactions in the response to selection and the evolution of evolvability. Theoretical Population Biology 68:179–196. doi: 10.1016/j.tpb.2005.05.002

Catchen JM, Hohenlohe PA, Bernatchez L, et al (2017) Unbroken: RADseq remains a powerful tool for understanding the genetics of adaptation in natural populations. Molecular Ecology Resources 17:362–365. doi: 10.1111/1755-0998.12669

Charlesworth, B. 2013. Background selection 20 years on: The Wilhelmine E. Key 2012 Invitational Lecture. Journal of Heredity 104: 161–171.

Charlesworth B, Charlesworth D (2010) Elements of Evolutionary Genetics. Greenwood Village, CO: Roberts and Company Publishers. 734 pp.

Chaves, J. A., Cooper, E. A., Hendry, A. P., Podos, J., De León, L. F., Raeymaekers, J. A. M., et al. (2016). Genomic variation at the tips of the adaptive radiation of Darwin’s finches. Molecular Ecology, 25(21), 5282–5295. http://doi.org/10.1111/mec.13743

Cheverud JM, Routman EJ (1995) Epistasis and its contribution to genetic variance components. Genetics 139:1455–1461.

Chevin L-M (2012) Genetic constraints on adaptation to a changing environment. Evolution 67:708–721. doi: 10.1111/j.1558-5646.2012.01809.x

Chevin L-M, Hoffmann AA (2017) Evolution of phenotypic plasticity in extreme environments. Philosophical Transactions of the Royal Society B: Biological Sciences 372:20160138–12. doi: 10.1098/rstb.2016.0138

Chevin L-M, Lande R, Mace GM (2010a) Adaptation, Plasticity, and Extinction in a Changing Environment: Towards a Predictive Theory. PLoS Biol 8:e1000357–8. doi: 10.1371/journal.pbio.1000357

Chevin L-M, Martin G, Lenormand T (2010b) Fisher’s model and the genomics of adaptation: Restricted pleiotropy, heterogenous mutation, and parallel evolution. Evolution 64:3213–3231. doi: 10.1111/j.1558-5646.2010.01058.x

Chevin L-M, Hospital F (2008) Selective sweep at a quantitative trait locus in the presence of background genetic variation. Genetics 180:1645–1660. doi: 10.1534/genetics.108.093351

Chevin L-M, Lande R (2011) Adaptation to marginal habitats by evolution of increased phenotypic plasticity. Journal of Evolutionary Biology 24:1462–1476. doi: 10.1111/j.1420-9101.2011.02279.x

Cheplick GP (2015) Approaches to Plant Evolutionary Ecology. Oxford University Press, USA.

Civelek M, Lusis AJ (2013) Systems genetics approaches to understand complex traits. Nature Publishing Group 15:34–48. doi: 10.1038/nrg3575

Cockerham CC (1954) An Extension of the Concept of Partitioning Hereditary Variance for Analysis of Covariances among Relatives When Epistasis Is Present. Genetics 39:859–882.

Cohen D, Bogeat-Triboulot M-B, Tisserant E, et al. (2010) Comparative transcriptomics of drought responses in Populus: a meta-analysis of genome-wide expression profiling in mature leaves and root apices across two genotypes. BMC Genomics 11:630. doi: 10.1186/1471-2164-11-630

Collins S, de Meaux J, Acquisti C (2007) Adaptive walks toward a moving optimum. Genetics 176:1089–1099. doi: 10.1534/genetics.107.072926

Comeault AA, Flaxman SM, Riesch R, et al. (2015) Selection on a genetic polymorphism counteracts ecological speciation in a stick insect. Current Biology 25:1975–1981. doi: 10.1016/j.cub.2015.05.058

Comeault AA, Soria-Carrasco V, Gompert Z, et al. (2014) Genome-wide association mapping of phenotypic traits subject to a range of intensities of natural selection in *Timema cristinae*. The American Naturalist 183:711–727. doi: 10.1086/675497

Cornelius J (1994) Heritabilities and additive genetic coefficients of variation in forest trees. Can J For Res 24:372–379. doi: 10.1139/x94-050

Costanza R, d’Arge R, De Groot R, et al. (1997) The value of the world’s ecosystem services and natural capital. Nature 387:253–260. doi: http://dx.doi.org/10.1038/387253a0

Cowen L, Ideker T, Raphael BJ, Sharan R (2017) Network propagation: a universal amplifier of genetic associations. Nat Ecol Evol 18:551–562. doi: 10.1038/nrg.2017.38

Cronn R, Dolan PC, Jogdeo S, et al. (2017) Transcription Through The Eye Of A Needle: Daily And Annual Cycles Of Gene Expression Variation In Douglas-Fir Needles. bioRxiv. doi: 10.1101/117374

Crow JF (2010) On epistasis: why it is unimportant in polygenic directional selection. Philosophical Transactions of the Royal Society B: Biological Sciences 365:1241–1244. doi: 10.1098/rstb.2009.0275

Crow JF (2008) Maintaining evolvability. Journal of genetics 87:349–353. doi: 10.1007/s12041-008-0057-8

Cruickshank TE, Hahn MW (2014) Reanalysis suggests that genomic islands of speciation are due to reduced diversity, not reduced gene flow. Molecular Ecology 23:3133–3157. doi: 10.1111/mec.12796

Csilléry K, Lalagüe H, Vendramin GG, González-Martínez SC, Fady B, Oddou-Muratorio S (2014) Detecting short spatial scale local adaptation and epistatic selection in climate-related candidate genes in European beech *(Fagus sylvatica)* populations. Molecular Ecology 23:4696–4708.

La Torre De AR, Li Z, Van de Peer Y, Ingvarsson PK (2017) Contrasting Rates of Molecular Evolution and Patterns of Selection among Gymnosperms and Flowering Plants. Molecular Biology and Evolution 34:1363–1377. doi: 10.1093/molbev/msx069

De Mita S, Thuillet A-C, Gay L, et al. (2013) Detecting selection along environmental gradients: analysis of eight methods and their effectiveness for outbreeding and selfing populations. Molecular Ecology 22:1383–1399. doi: 10.1111/mec.12182

Devey ME, Fiddler TA, Liu BH, Knapp SJ, Neale DB (1994) An RFLP linkage map for loblolly pine based on a three-generation outbred pedigree. Theor Appl Genet 88:273–278

de Villemereuil P, Gaggiotti OE, Mouterde M, Till-Bottraud I (2015) Common garden experiments in the genomic era: new perspectives and opportunities. Heredity 116:249–254. doi: 10.1038/hdy.2015.93

de Visser JAGM, Cooper TF, Elena SF (2011) The causes of epistasis. Proc Biol Sci 278:3617–3624. doi: 10.1098/rspb.2011.1537

de Vladar HP, Barton N (2014) Stability and response of polygenic traits to stabilizing selection and mutation. Genetics. doi: 10.1534/genetics.113.159111/-/DC1

Devlin B, Roeder K (1999) Genomic control for association studies. Biometrics 55:997–1004.

Dickson, S. P., Wang, K., Krantz, I., Hakonarson, H., & Goldstein, D. B. (2010). Rare variants create synthetic genome-wide associations. PLoS biology, 5(1), e1000294.

Dittmar EL, Oakley CG, Conner JK, et al. (2016) Factors influencing the effect size distribution of adaptive substitutions. Proc Biol Sci 283:20153065–8. doi: 10.1098/rspb.2015.3065

Donohue K, Rubio de Casas R, Burghardt L, Kovach K, Willis CG (2010) Germination, postgermination adaptation, and species ecological ranges. Annual Review of Ecology Evolution and Systematics 41:293–319.

Du, J., & Groover, A. (2010, January). Transcriptional regulation of secondary growth and wood formation. Journal of Integrative Plant Biology.

East EM (1910) A Mendelian interpretation of variation that is apparently continuous. The American Naturalist 44:65–82.

Eckert AJ, Bower AD, Jermstad KD, et al. (2013a) Multilocus analyses reveal little evidence for lineage-wide adaptive evolution within major clades of soft pines (*Pinus* subgenus Strobus). Molecular Ecology 22:5635–5650. doi: 10.1111/mec.12514

Eckert AJ, Maloney PE, Vogler DR, et al. (2015) Local adaptation at fine spatial scales: an example from sugar pine *(Pinus lambertiana, Pinaceae)*. Tree Genetics & Genomes 11:117.

Eckert AJ, Pande B, Ersoz ES, Wright MH, Rashbrook VK, Nicolet CM, Neale DB (2009) High-throughput genotyping and mapping of single nucleotide polymorphisms in loblolly pine (*Pinus taeda* L.). Tree Genetics & Genomes 5:225–234. doi: 10.1007/s11295-008-0183-8

Eckert AJ, Wegrzyn JL, Cumbie WP, et al. (2012) Association genetics of the loblolly pine (Pinus taeda, Pinaceae) metabolome. New Phytologist 193:890–902. doi: 10.1111/j.1469-8137.2011.03976.x

Eckert AJ, Wegryzn JL, Liechty JD, et al. (2013b) The Evolutionary Genetics of the Genes Underlying Phenotypic Associations for Loblolly Pine *(Pinus taeda, Pinaceae)*. Genetics 195:1353–1372. doi: 10.1534/genetics.113.157198/-/DC1

Ehret GB, Lamparter D, Hoggart CJ, Whittaker JC, Beckmann JS, Kutalik Z, Genetic Investigation of Anthropometric Traits Consortium (2012) A multi-SNP locus-association method reveals a substantial fraction of the missing heritability. The American Journal of Human Genetics, 91, 863–871.

Eyre-Walker A, Keightley PD (2007) The distribution of fitness effects of new mutations. Nature Reviews Genetics 8:610–618. doi: 10.1038/nrg2146

Evans, L. M., Kaluthota, S., Pearce, D. W., Allan, G. J., Floate, K., Rood, S. B., & Whitham, T. G. (2016). Bud phenology and growth are subject to divergent selection across a latitudinal gradient in *Populus angustifolia* and impact adaptation across the distributional range and associated arthropods. Ecology and evolution 6:4565–4581. doi: 10.1002/ece3.2222

Falconer DS (1989) Introduction to quantitative genetics, 3d ed. Longman, New York.

Faltus FA (2014) Systems genetics: A paradigm to improve discovery of candidate genes and mechanisms underlying complex traits. Plant Science 223: 45–48. http://dx.doi.org/10.1016/j.plantsci.2014.03.003

Feder JL, Egan SP, Nosil P (2012a) The genomics of speciation-with-gene-flow. Trends in Genetics 28:342–350. doi: 10.1016/j.tig.2012.03.009

Feder JL, Gejji R, Yeaman S, Nosil P (2012b) Establishment of new mutations under divergence and genome hitchhiking. Philosophical Transactions of the Royal Society B: Biological Sciences 367:461–474. doi: 10.1038/nature08480

Feder JL, Nosil P (2010) The efficacy of divergence hitchhiking in generating genomic islands during ecological speciation. Evolution 64:1729–1747. doi: 10.1111/j.1558-5646.2009.00943.x

Feldman M, Lewontin R (1975) The heritability hang-up. Science 190:1163–1168. doi: 10.1126/science.1198102

Felsenstein J (1976) The theoretical population genetics of variable selection and migration. Annu Rev Genet 10:253–280.

Fisher RA (1918) The correlation between relatives on the supposition of Mendelian inheritance. Transactions of the Royal Society of Edinburgh 52:399–433.

Fisher RA (1930) The genetical theory of natural selection: a complete variorum edition. Oxford University Press

Forester BR, Lasky JR, Wagner HH, Urban DL (2017) Using genotype-environment associations to identify multilocus local adaptation. bioRxiv 1–24. doi: 10.1101/129460

Franks SJ, Weber JJ, Aitken SN (2013) Evolutionary and plastic responses to climate change in terrestrial plant populations. Evolutionary Applications 7:123–139. doi: 10.1111/eva.12112

Friedline CJ, Lind BM, Hobson EM, et al. (2015) The genetic architecture of local adaptation I: the genomic landscape of foxtail pine (Pinus balfouriana Grev. & Balf.) as revealed from a high-density linkage map. Tree Genetics & Genomes 11:49. doi: 10.1007/s11295-015-0866-x

Gagnaire P-A, Gaggiotti OE (2016) Detecting polygenic selection in marine populations by combining population genomics and quantitative genetics approaches. Curr Zool 62:603–616. doi: 10.1093/cz/zow088

Gazal S, Finucane HK, Furlotte NA, Loh P, Palamara PF, Liu X, Schoech A, Bulik-Sullivan B, Neale BM, Gusev A, Price A (2017) Linkage disequilibrium–dependent architecture of human complex traits shows action of negative selection. Nat. Gen. 1–7. doi: 10.1038/ng.3954

Gibson G (2012) Rare and common variants: twenty arguments. Nature Reviews Genetics 13:135–145. doi: 10.1038/nrg3118

Gilbert, K. J., & Whitlock, M. C. (2015). *Q*_ST_–*F*_ST_ comparisons with unbalanced half-sib designs. Molecular ecology resources 15:262–267.

Goddard ME, Wray NR, Verbyla K, Visscher PM (2009) Estimating effects and making predictions from genome-wide marker data. Statistical Science 24:517–529.

Gompert, Z., Egan, S. P., Barrett, R. D. H., Feder, J. L., & Nosil, P. (2016). Multilocus approaches for the measurement of selection on correlated genetic loci. Molecular Ecology, 1–18. http://doi.org/10.1111/mec.13867

Gompert, Z., Jahner, J. P., Scholl, C. F., Wilson, J. S., Lucas, L. K., Soria-Carrasco, V., et al. (2015). The evolution of novel host use is unlikely to be constrained by trade-offs or a lack of genetic variation. Molecular Ecology, 24(11), 2777–2793. http://doi.org/10.1111/mec.13199

Goodnight CJ (1988) Epistasis and the effect of founder events on the additive genetic variance. Evolution 42:441–454. doi: 10.2307/2409030

Göring HHH, Terwilliger JD, Blangero J (2001) Large Upward Bias in Estimation of Locus-Specific Effects from Genomewide Scans. The American Journal of Human Genetics 69:1357–1369. doi: 10.1086/324471

Grandtner, M. M. (2005). Elsevier’s Dictionary of Trees: Volume 1: North America. Elsevier.

Grattapaglia D (2017) Status and perspectives of genomic selection in forest tree breeding. In ME Sorrells, Genomic Selection for Crop Improvement (pp. 199–257). Cham, Switzerland: Springer.

Grattapaglia D, Resende MDV (2011) Genomic selection in forest tree breeding. Tree Genetics & Genomes 7: 241–255.

Grattapaglia D, Sederoff R (1994) Genetic linkage maps of Eucalyptus grandis and Eucalyptus urophylla using a pseudo-test cross: mapping strategy and RAPD markers. Genetics 137:1121–1137

Griswold CK (2015) Additive genetic variation and evolvability of a multivariate trait can be increased by epistatic gene action. J Theor Biol 387:241–257. doi: 10.1016/j.jtbi.2015.09.023

Guan, Y., & Stephens, M. (2011). Bayesian variable selection regression for genome-wide association studies and other large-scale problems. Annals of Eugenics, 5(3), 1780–1815. http://doi.org/10.1214/11-AOAS455

Haldane JBS (1930) A mathematical theory of natural and artificial selection. (Part VI, Isolation.). 26:220–230.

Hall D, Hallingbäck HR, Wu HX (2016) Estimation of number and size of QTL effects in forest tree traits. Tree Genetics & Genomes 1–17. doi: 10.1007/s11295-016-1073-0

Hansen TF (2006) The Evolution of Genetic Architecture. Annual Review of Ecology, Evolution, and Systematics 37:123–157. doi: 10.1146/annurev.ecolsys.37.091305.110224

Hansen TF, Pelabon C, Houle D (2011) Heritability is not evolvability. Evolutionary Biology 38:258–277

Hansen TF (2013) Why epistasis is important for selection and adaptation. Evolution 67:3501–3511. doi: 10.1111/evo.12214

Hansen TF (2003) Is modularity necessary for evolvability? Biosystems 69:83–94. doi: 10.1016/S0303-2647(02)00132-6

Hansen TF, Wagner GP (2001) Modeling genetic architecture: a multilinear theory of gene interaction. Theoretical Population Biology 59:61–86. doi: 10.1006/tpbi.2000.1508

Heffner EL, Sorrells ME, Jannink JL (2009) Genomic selection for crop improvement. Crop Science 49: 1–12.

Hemani G, Knott S, Haley C (2013) An Evolutionary Perspective on Epistasis and the Missing Heritability. PLoS Genetics 9:e1003295. doi: 10.1371/journal.pgen.1003295

Henderson CR (1975) Best linear unbiased estimation and prediction under a selection model. Biometrics 31:423–447.

Hendry AP (2002) QST > = = < FST? Trends in Ecology & Evolution 17:502–502. doi: http://dx.doi.org/10.1016/S0169-5347(02)02603-4

Hendry AP (2016) Key Questions on the Role of Phenotypic Plasticity in Eco-Evolutionary Dynamics. Journal of Heredity 107:25–41. doi: 10.1093/jhered/esv060

Hereford J (2009) A Quantitative Survey of Local Adaptation and Fitness Trade-Offs. The American Naturalist 173:579–588. doi: 10.1086/597611

Hermisson J (2009) Who believes in whole-genome scans for selection? Heredity 103:283–284. doi: 10.1038/hdy.2009.101

Hermisson J, Hansen TF, Wagner GP (2003) Epistasis in Polygenic Traits and the Evolution of Genetic Architecture under Stabilizing Selection. American Naturalist 161:708–734. doi: 10.1086/374204

Hermisson J, Pennings PS (2005) Soft sweeps: molecular population genetics of adaptation from standing genetic variation. Genetics 169:2335–2352. doi: 10.1534/genetics.104.036947

Hermisson J, Pennings PS (2017) Soft sweeps and beyond: understanding the patterns and probabilities of selection footprints under rapid adaptation. Methods in Ecology and Evolution 8:700–716. doi: 10.1111/2041-210X.12808

Hill WG (2010) Understanding and using quantitative genetic variation. Philosophical Transactions of the Royal Society B 365:73–85.

Hill WG, Goddard ME, Visscher PM (2008) Data and Theory Point to Mainly Additive Genetic Variance for Complex Traits. PLoS Genetics 4:e1000008–10. doi: 10.1371/journal.pgen.1000008

Hirschhorn JN, Daly MJ (2005) Genome-wide association studies for common diseases and complex traits. Nature Reviews Genetics 6:95–108.

Hoban S, Kelley JL, Lotterhos KE, et al. (2016) Finding the Genomic Basis of Local Adaptation: Pitfalls, Practical Solutions, and Future Directions. The American Naturalist 188:000–000. doi: 10.1086/688018

Hodgins KA, Yeaman S, Nurkowski KA, et al. (2016) Expression Divergence Is Correlated with Sequence Evolution but Not Positive Selection in Conifers. Molecular Biology and Evolution 33:1502–1516. doi: 10.1093/molbev/msw032

Hoffmann AA, Rieseberg LH (2008) Revisiting the Impact of Inversions in Evolution: From Population Genetic Markers to Drivers of Adaptive Shifts and Speciation? Annual Review of Ecology, Evolution, and Systematics 39:21–42. doi: 10.1146/annurev.ecolsys.39.110707.173532

Holliday JA, Aitken SN, Cooke JEK et al. (2017) Advances in ecological genomics in forest trees and applications to genetic resources conservation and breeding. Molecular Ecology 26:706–717.

Holliday, J. A., Wang, T., & Aitken, S. N. (2012). Predicting adaptive phenotypes from multilocus genotypes in Sitka spruce *(Picea sitchensis)* using random forest. G3: Genes, Genomes, Genetics. http://doi.org/10.1534/g3.112.002733/-/DC1

Holliday JA, Zhou L, Bawa R, et al. (2016) Evidence for extensive parallelism but divergent genomic architecture of adaptation along altitudinal and latitudinal gradients in *Populus trichocarpa*. New Phytologist 209:1240–1251.

Hornoy, B., Pavy, N., Gérardi, S., Beaulieu, J., & Bousquet, J. (2015). Genetic Adaptation to Climate in White Spruce Involves Small to Moderate Allele Frequency Shifts in Functionally Diverse Genes. Genome Biology and Evolution, 7(12), 3269–3285. http://doi.org/10.1093/gbe/evv218

Howe GT, Aitken SN, Neale DB, et al. (2003) From genotype to phenotype: unraveling the complexities of cold adaptation in forest trees. Can J Bot 81:1247–1266. doi: 10.1139/b03-141

Huang W, Mackay TFC (2016) The Genetic Architecture of Quantitative Traits Cannot Be Inferred from Variance Component Analysis. PLoS Genetics 12:e1006421–15. doi: 10.1371/journal.pgen.1006421

Huber, C. D., Durvasula, A., Hancock, A. M., & Lohmueller, K. E. (2017). Gene expression drives the evolution of dominance. bioRxiv, 182865. doi: http://dx.doi.org/10.1101/182865

Ingvarsson PK, Hvidsten TR, Street NR (2016) Towards integration of population and comparative genomics in forest trees. New Phytologist 212:338–344. doi: 10.1111/nph.14153

Innan H, Kim Y (2004) Pattern of polymorphism after strong artificial selection in a domestication event. Proceedings of the National Academy of Sciences 101:10667–10672. doi: 10.1073/pnas.0401720101

Isik F, Kumar S, Martínez-García PJ, et al. (2015) Acceleration of Forest and Fruit Tree Domestication by Genomic Selection. Advances in Botanical Research. 74: 93–124. doi: http://dx.doi.org/10.1016/bs.abr.2015.05.002

Iwata et al. (2011) prospects for genomic selection in conifer breeding: a simulation study of Cryptomeria japonica. Tree Genetics & Genomes 7: 747–758.

Jain K, Stephan W (2015) Response of polygenic traits under stabilizing selection and mutation when loci have unequal effects. G3: Genes | Genomes | Genetics 5:1065–1074. doi: 10.1534/g3.115.017970

Jain K, Stephan W (2017) Rapid Adaptation of a Polygenic Trait After a Sudden Environmental Shift. Genetics 206:389–406. doi: 10.1534/genetics.116.196972

Jansen RC, Tesson BM, Fu J, et al. (2009) Defining gene and QTL networks. Current Opinion in Plant Biology 12:241–246. doi: 10.1016/j.pbi.2009.01.003

Jensen JD (2014) On the unfounded enthusiasm for soft selective sweeps. Nature Communications 5:5281. doi: 10.1038/ncomms6281

Jensen JD, Kim Y, DuMont VB, et al. (2005) Distinguishing between selective sweeps and demography using DNA polymorphism data. Genetics 170:1401–1410. doi: 10.1534/genetics.104.038224

Johnson RC, Nelson GW, Troyer JL, et al. (2010) Accounting for multiple comparisons in a genome-wide association study (GWAS). BMC Genomics 11:724.

Jones AG, Bürger R, Arnold SJ (2014) Epistasis and natural selection shape the mutational architecture of complex traits. Nature Communications 5:1–10. doi: 10.1038/ncomms4709

Joo JWJ, Hormozdiari F, Han B, Eskin E (2016) Multiple testing correction in linear mixed models. Genome Biology 17:62.

Josephs EB, Wright SI, et al. (2017a) The Relationship between Selection, Network Connectivity, and Regulatory Variation within a Population of *Capsella grandiflora*. Genome Biology and Evolution 9:1099–1109. doi: 10.1093/gbe/evx068

Josephs EB, Lee YW, Stinchcombe JR, Wright SI (2015) Association mapping reveals the role of purifying selection in the maintenance of genomic variation in gene expression. Proceedings of the National Academy of Sciences 112:15390–15395. doi: 10.1073/pnas.1503027112

Josephs EB, Stinchcombe JR, Wright SI (2017b) What can genome-wide association studies tell us about the evolutionary forces maintaining genetic variation for quantitative traits? New Phytologist 214:21–33. doi: 10.1111/nph.14410

Kaplan NL, Hudson RR, Langley CH (1989) The “hitchhiking effect” revisited. Genetics 123:887–899.

Kawecki TJ (2008) Adaptation to Marginal Habitats. Annual Review of Ecology, Evolution, and Systematics 39:321–342. doi: 10.1146/annurev.ecolsys.38.091206.095622

Kawecki TJ, Ebert D (2004) Conceptual issues in local adaptation. Ecology Letters 7:1225–1241. doi: 10.1111/j.1461-0248.2004.00684.x

Keightley PD, Eyre-Walker A (2010) What can we learn about the distribution of fitness effects of new mutations from DNA sequence data? Philos Trans R Soc Lond, B, Biol Sci 365:1187–1193. doi: 10.1098/rstb.2009.0266

Kempthorne O (1954) The Correlation between Relatives in a Random Mating Population. Proceedings of the Royal Society B: Biological Sciences 143:103–113. doi: 10.1098/rspb.1954.0056

Kimura M (1983) The neutral theory of molecular evolution. Cambridge University Press

Kinloch Jr, B. B., Parks, G. K., & Fowler, C. W. (1970). White pine blister rust: simply inherited resistance in sugar pine. Science, 193–195.

Kirkpatrick M, Barton NH (1997) Evolution of a species’ range. The American Naturalist 150:1–23. doi: 10.1086/286054

Kirkpatrick M, Barton NH (2006) Chromosome Inversions, Local Adaptation and Speciation. Genetics 173:419–434. doi: 10.1534/genetics.105.047985

Kopp M, Hermisson J (2009a) The Genetic Basis of Phenotypic Adaptation I: Fixation of Beneficial Mutations in the Moving Optimum Model. Genetics 182:233–249. doi: 10.1534/genetics.108.099820

Kopp M, Hermisson J (2009b) The genetic basis of phenotypic adaptation II: the distribution of adaptive substitutions in the moving optimum model. Genetics 183:1453–1476. doi: 10.1534/genetics.109.106195

Kopp M, Matuszewski S (2013) Rapid evolution of quantitative traits: theoretical perspectives. Evolutionary Applications 7:169–191. doi: 10.1111/eva.12127

Kremer A, Le Corre V (2012) Decoupling of differentiation between traits and their underlying genes in response to divergent selection. Heredity 108:375–385. doi: 10.1038/hdy.2011.81

Kremer A, Ronce O, Robledo-Arnuncio JJ, et al. (2012). Long-distance gene flow and adaptation of forest trees to rapid climate change. Ecology Letters 15:378–392.

Krutovsky KV, Neale DB (2005) Nucleotide diversity and linkage disequilibrium in cold-hardiness- and wood quality-related candidate genes in Douglas fir. Genetics 171:2029–2041. doi: 10.1534/genetics.105.044420

Lamichhaney S, Berglund J, Almén MS, et al. (2015) Evolution of Darwin’s finches and their beaks revealed by genome sequencing. Nature 518:371–375. doi: 10.1038/nature14181

Lamy JB, Bouffier L, Burlett R, Plomion C, Cochard H, Delzon S (2011) Uniform selection as a primary force reducing population genetic differentiation of cavitation resistance across a species range. PLoS One 6:e23476.

Lamy JB, Plomion C, Kremer A, Delzon S (2012) QST< FST as a signature of canalization. Molecular Ecology 21:5646–5655.

Lande R (1980) The genetic covariance between characters maintained by pleiotropic mutations. Genetics 94:203–215.

Lande R (2009) Adaptation to an extraordinary environment by evolution of phenotypic plasticity and genetic assimilation. Journal of Evolutionary Biology 22:1435–1446. doi: 10.1111/j.1420-9101.2009.01754.x

Lande R, Arnold S (1983) The measurement of selection on correlated characters. Evolution, 37, 1210–1226. doi: 10.1111/j.1558-5646.1983.tb00236.x

Langlet O (1971) Two hundred years genecology. Taxon 20:653–721. doi: 10.2307/1218596

Latta RG (1998) Differentiation of allelic frequencies at quantitative trait loci affecting locally adaptive traits. The American Naturalist 151:283–292. doi: 10.1086/286119

Latta RG (2003) Gene flow, adaptive population divergence and comparative population structure across loci. New Phytologist 161:51–58. doi: 10.1046/j.1469-8137.2003.00920.x

Laurent S, Pfeifer SP, Settles ML, et al. (2016) The population genomics of rapid adaptation: disentangling signatures of selection and demography in white sands lizards. Molecular Ecology 25:306–323. doi: 10.1111/mec.13385

Lauteri M, Pliura A, Monteverdi MC, Brugnoli E, Villani F, Eriksson G (2004) Genetic variation in carbon isotope discrimination in six European populations of *Castanea sativa* Mill. originating from contrasting localities. Journal of evolutionary biology 17:1286–1296.

Le Corre V, Kremer A (2012) The genetic differentiation at quantitative trait loci under local adaptation. Molecular Ecology 21:1548–1566. doi: 10.1111/j.1365-294X.2012.05479.x

Le Rouzic A, Álvarez-Castro JM (2016) Epistasis-induced evolutionary plateaus in selection responses. American Naturalist 188:E134–E150. doi: 10.1086/688893

Leempoel, K., Duruz, S., Rochat, E., Widmer, I., Orozco-terWengel, P., & Joost, S. (2017). Simple rules for an efficient use of Geographic Information Systems in molecular ecology. Frontiers in Ecology and Evolution 5: 1–10.

Legendre, P., & Legendre, L. F. (2012). Numerical ecology (Vol. 24). Elsevier.

Leimu R, Fischer M (2008) A meta-analysis of local adaptation in plants. PLoS ONE 3:e4010. doi: 10.1371/journal.pone.0004010.s001

Leinonen T, McCairns RJS, O’Hara RB, Merilä J (2013) QST-FST comparisons: evolutionary and ecological insights from genomic heterogeneity. Nature Reviews Genetics 14:179–190. doi: 10.1038/nrg3395

Leinonen PH, Sandring S, Quilot B, et al. (2009). Local adaptation in european populations of *Arabidopsis lyrata* (Brassicaceae). American Journal of Botany 96:1129–1137.

Leiserson M, Eldridge JV, Ramachandran S (2013) Network analysis of GWAS data. Current opinion in Genetics and Development 23:602–610 doi: 10.1016/j.gde.2013.09.003

Leitch AR, Leitch IJ (2012) Ecological and genetic factors linked to contrasting genome dynamics in seed plants. New Phytologist 194:629–646. doi: 10.1111/j.1469-8137.2012.04105.x.

Lenormand T (2002) Gene flow and the limits to natural selection. Trends in Ecology & Evolution 17:183–189. doi: 10.1016/S0169-5347(02)02497-7

Lewontin RC, Krakauer J (1973) Distribution of gene frequency as a test of the theory of the selective neutrality of polymorphisms.

Li Y, Suontoma M, Burdon RD, Dungey HS (2017) Genotype by environment interactions in forest tree breeding: review of methodology and perspectives on research and application. Tree Genetics & Genomes 13:60. doi: 10.1007/s11295-017-1144-x

Lind BM, Friedline CJ, Wegrzyn JL, et al. (2017) Water availability drives signatures of local adaptation in whitebark pine (*Pinus albicaulis* Engelm.) across fine spatial scales of the Lake Tahoe Basin, USA. Molecular Ecology 26:3168–3185 doi: 10.1111/mec.14106

Loehle C (1988) Tree life history strategies: the role of defenses. Canadian Journal of Forest Research 18: 209–222.

Long AD, Langley CH (1999) The power of association studies to detect the contribution of candidate genetic loci to variation in complex traits. Genome Research 9:720–731.

Lopez GA, Potts BM, Vaillancourt RE, Apiolaza LA (2003). Maternal and carryover effects on early growth of Eucalyptus globulus. Canadian Journal of Forest Research, 33(11), 2108–2115. http://doi.org/10.1139/X03-132

Liu J-J (2017) Profiling methyl jasmonate-responsive transcriptome for understanding induced systemic resistance in whitebark pine (*Pinus albicaulis*). Plant Mol Biol 0:0–0. doi: 10.1007/s11103-017-0655-z&domain=pdf

Liu J-J, Schoettle AW, Sniezko RA, et al. (2016) Genetic mapping of *Pinus flexilis* major gene (Cr4) for resistance to white pine blister rust using transcriptome-based SNP genotyping. BMC Genomics 17:1–12. doi: 10.1186/s12864-016-3079-2

Lotterhos KE, Hodges K, Yeaman S, Degner J, Aitken S (2017) Modular environmental pleiotropy of genes involved in local adaptation to climate despite physical linkage. bioRxiv doi: https://doi.org/10.1101/202481

Lotterhos KE, Whitlock MC (2014) Evaluation of demographic history and neutral parameterization on the performance of *F*_ST_ outlier tests. Molecular Ecology 23:2178–2192. doi: 10.1111/mec.12725

Lotterhos KE, Whitlock MC (2015) The relative power of genome scans to detect local adaptation depends on sampling design and statistical method. Molecular Ecology 24:1031–1046.

Lowry DB, Hoban S, Kelley JL, et al. (2016) Breaking RAD: an evaluation of the utility of restriction site-associated DNA sequencing for genome scans of adaptation. Molecular Ecology Resources 17:142–152. doi: 10.1111/1755-0998.12635

Lowry DB, Hoban S, Kelley JL, et al. (2017) Responsible RAD: Striving for best practices in population genomic studies of adaptation. Molecular Ecology Resources 17:366–369. doi: 10.1111/1755-0998.12677

Lynch M, Walsh B (1998) Genetics and analysis of quantitative traits. Sinauer Sunderland, MA

Mackay T (2001) The genetic architecture of quantitative traits. Annual review of genetics, 35, 303–339.

Mackay TFC (2014) Epistasis and quantitative traits: using model organisms to study gene-gene interactions. Nature Reviews Genetics 15:22–33. doi: 10.1038/nrg3627

Mackay TFC, Stone EA, Ayroles JF (2009) The genetics of quantitative traits: challenges and prospects. Nature Reviews Genetics 10:565–577. doi: 10.1038/nrg2612

MacPherson A, Hohenlohe PA, Nuismer SL (2015) Trait dimensionality explains widespread variation in local adaptation. Proc Biol Sci 282:20141570–20141570. doi: 10.1098/rspb.2014.1570

Mahalovich MF, Hipkins VD (2011) Molecular genetic variation in whitebark pine (Pinus albicaulis Engelm.) in the Inland West. In: Keane, Robert E.; Tomback, Diana F.; Murray, Michael P.; and Smith, Cyndi M., eds. 2011. The future of high-elevation, five-needle white pines in Western North America: Proceedings of the High Five Symposium. 28-30 June 2010; Missoula, MT. Proceedings RMRS-P-63. Fort Collins, CO: U.S. Department of Agriculture, Forest Service, Rocky Mountain Research Station. 376 p.

Mähler N, Wang J, Terebieniec BK, et al. (2017) Gene co-expression network connectivity is an important determinant of selective constraint. PLoS Genetics 13:e1006402–33. doi: 10.1371/journal.pgen.1006402

Mahler DL, Weber MG, Wagner CE, Ingram T (2017) Pattern and Process in the Comparative Study of Convergent Evolution. American Naturalist 190:S13–S28. doi: 10.1086/692648

Mäki-Tanila A, Hill WG (2014) Influence of gene interaction on complex trait variation with multilocus models. Genetics 198:355–367. doi: 10.1534/genetics.114.165282

Martin G, Lenormand T (2006) A general multivariate extension of Fisher’s geometrical model and the distribution of mutation fitness effects across species. Evolution 60:893–16. doi: 10.1554/05-412.1

Martin G, Lenormand T (2008) The distribution of beneficial and fixed mutation fitness effects close to an optimum. Genetics 179:907–916. doi: 10.1534/genetics.108.087122

Matuszewski S, Hermisson J, Kopp M (2015) Catch me if you can: Adaptation from standing genetic variation to a moving phenotypic optimum. Genetics 200:1255–1274. doi: 10.1534/genetics.115.178574/-/DC1

Matuszewski S, Hermisson J, Kopp M (2014) Fisher’s Geometric Model with a moving optimum. Evolution 68:2571–2588. doi: 10.1111/evo.12465

Mátyás C (1996) Climatic adaptation of trees: rediscovering provenance tests. Euphytica 92:45–54.

Maynard Smith JH, Haigh J (1974) The hitch-hiking effect of a favourable gene. Genetical Research 23:23–35.

McCandlish DM, Stoltzfus A (2014) Modeling evolution using the probability of fixation: history and implications. The Quarterly review of biology 89:225–252. doi: 10.1086/677571

McKay JK, Latta RG (2002) Adaptive population divergence: markers, QTL and traits. Trends in Ecology & Evolution 17:285–291.

McKinney GJ, Larson WA, Seeb LW, Seeb JE (2017) RADseq provides unprecedented insights into molecular ecology and evolutionary genetics: comment on Breaking RAD by Lowry et al. (2016). Molecular Ecology Resources 17:356–361. doi: 10.1111/1755-0998.12649

Mei W, Stetter MG, Gates DJ, Stitzer MC, Ross-Ibarra J (2017) Adaptation in plant genomes: bigger is different. bioRxiv https://doi.org/10.1101/196501

Meier JI, Sousa VC, Marques DA, et al. (2016) Demographic modelling with whole-genome data reveals parallel origin of similar Pundamilia cichlid species after hybridization. Molecular Ecology 26:123–141. doi: 10.1111/mec.13838

Messer PW, Petrov DA (2013) Population genomics of rapid adaptation by soft selective sweeps. Trends in Ecology & Evolution 28:659–669. doi: 10.1016/j.tree.2013.08.003

Meuwissen TH, Hayes BJ, Goddard ME. 2001. Prediction of total genetic value using genome-wide dense marker maps. Genetics 157: 1819–1829.

Mitton JB, Grant MC, Yoshino AM (1998) Variation in allozymes and stomatal size in pinyon (*Pinus edulis*, Pinaceae), associated with soil moisture. American Journal of Botany 85:1262–1265.

Mitton JB, Williams CG (2006) Gene Flow in Conifers. In: Landscapes, Genomics, and Transgenic Conifers. (ed Williams CG), pp. 147–168. Springer Netherlands, Dordrecht.

Mizrachi E, Verbeke L, Christie N, et al. (2017) Network-based integration of systems genetics data reveals pathways associated with lignocellulosic biomass accumulation and processing. Proc Natl Acad Sci 114:1195–1200. doi: 10.1073/pnas.1620119114

Moreno G (1994) Genetic architecture, genetic behavior, and character evolution. Annual review of ecology and systematics 25:31–44.

Morgenstern EK (1996) Geographic variation in forest trees: genetic basis and application of knowledge in silviculture. UBC press.

Morse AM, Peterson DG, Islam-Faridi MN, et al. (2009) Evolution of genome size and complexity in *Pinus*. PLoS ONE 4:e4332.

Moser G, Lee SH, Hayes BJ, Goddard ME, Wray NR, Visscher PM (2015) Simultaneous discovery, estimation and prediction analysis of complex traits using a Bayesian mixture model. PLoS Genetics, 11, e1004969.

Namkoong G (1979) Introduction to Quantitative Genetics in Forestry. Technical Bulletin No. 1588. Washington, D. C. USDA Forest Service. 342 pp.

Neale DB, Kremer A. (2011). Forest tree genomics: growing resources and applications. Nature Reviews Genetics 12:111–122.

Neale DB, Langley CH, Salzberg SL, Wegrzyn JL (2013) Open access to tree genomes: the path to a better forest. Genome Biology 14:6 14:120. doi: 10.1186/gb-2013-14-6-120

Neale DB, Martínez-García PJ, La Torre De AR, et al. (2017) Novel Insights into Tree Biology and Genome Evolution as Revealed Through Genomics. Ann Rev of Plant Biol 68:457–483. doi :10.1146/annurev-arplant-042916-041049

Neale DB, Savolainen O (2004) Association genetics of complex traits in conifers. Trends in Plant Science 9:325–330.

Nelson RM, Pettersson ME, Carlborg Ö (2013) A century after Fisher: time for a new paradigm in quantitative genetics. Trends in Genetics 29:669–676. doi: 10.1016/j.tig.2013.09.006

Nilsson-Ehle H (1909) Kreuzungsuntersuchungen an Hafer und Weizen. Lunds Universitets Arsskrift 5:1–122.

Nystedt B, Street NR, Wetterbom A, et al. (2013) The Norway spruce genome sequence and conifer genome evolution. Nature 497:570–584.

Ohta T (1982) Linkage disequilibrium with the island model. Genetics 101:139–155.

Ohta, T. 1992. The nearly neutral theory of molecular evolution. Annu. Rev. Ecol. Syst. 23:263–286.

Ohta, T. 1996. The current significance and standing of neutral and nearly neutral theories. BioEssays 18: 673–684.

Oldfield, S., Lusty, C., & MacKinven, A. (1998). The world list of threatened trees. World Conservation Press.

Orr HA (1998) The population genetics of adaptation: the distribution of factors fixed during adaptive evolution. Evolution 52:935. doi: 10.2307/2411226

Orr HA (2005) The genetic theory of adaptation: a brief history. Nature Reviews Genetics 6:119–127. doi: 10.1038/nrg1523

Orr HA (2003) The Distribution of Fitness Effects Among Beneficial Mutations. Genetics 163:1519–1526. doi: 10.1101/SQB.1951.016.01.026

Orr HA (2006) The distribution of fitness effects among beneficial mutations in Fisher’s geometric model of adaptation. J Theor Biol 238:279–285. doi: 10.1016/j.jtbi.2005.05.001

Orr HA (2001) The “sizes” of mutations fixed in phenotypic evolution: a response to Clarke and Arthur. Evolution & Development 3:121–123. doi: 10.1046/j.1525-142x.2001.003003121.x

Orr HA (2000) Adaptation and the cost of complexity. Evolution 54:13–20. doi: 10.1111/j.0014-3820.2000.tb00002.x

Ortiz-Barrientos D, Engelstädter J, Rieseberg LH (2016) Recombination Rate Evolution and the Origin of Species. Trends in Ecology & Evolution 31:226–236. doi: 10.1016/j.tree.2015.12.016

Ovaskainen O, Karhunen M, Zheng CH, Cano Arias JM, Merilä J (2011) A new method to uncover signatures of divergent and stabilizing selection in quantitative traits. Genetics 189:621–632.

Paaby AB, Rockman MV (2013) The many faces of pleiotropy. Trends in Genetics 29:66–73. doi: 10.1016/j.tig.2012.10.010

Paixão T, Barton NH (2016) The effect of gene interactions on the long-term response to selection. Proc Natl Acad Sci USA 113:4422–4427. doi: 10.1073/pnas.1518830113

Palmé AE, Pyhajarvi T, Wachowiak W, Savolainen O (2009) Selection on Nuclear Genes in a Pinus Phylogeny. Molecular Biology and Evolution 26:893–905. doi: 10.1093/molbev/msp010

Parchman TL, Gompert Z, Mudge J, et al. (2012) Genome-wide association genetics of an adaptive trait in lodgepole pine. Molecular Ecology 21:2991–3005. doi: 10.1111/j.1365-294X.2012.05513.x

Parchman TL, Jahner JP, Uckele K, Galland LM (forthcoming) RADseq approaches and applications for forest tree genetics.

Pavlidis P, Metzler D, Stephan W (2012) Selective sweeps in multilocus models of quantitative traits. Genetics 192:225–239. doi: 10.1534/genetics.112.142547

Patterson HD, Thompson R (1971) Recovery of interblock information when block sizes are unequal. Biometrika 58:545–554.

Pennings PS, Hermisson J (2006a) Soft sweeps III: the signature of positive selection from recurrent mutation. PLoS Genetics 2:e186. doi: 10.1371/journal.pgen

Pennings PS, Hermisson J (2006b) Soft Sweeps II--Molecular Population Genetics of Adaptation from Recurrent Mutation or Migration. Molecular Biology and Evolution 23:1076–1084. doi: 10.1093/molbev/msj117

Petit RJ, Hampe A (2006) Some evolutionary consequences of being a tree. Annual review of ecology 37:187–214.

Phillips PC (2008) Epistasis—the essential role of gene interactions in the structure and evolution of genetic systems. Nature Reviews Genetics 9:855–867.

Piepho HP, Möhring J, Melchinger AE, Büchse A (2008) BLUP for phenotypic selection in plant breeding and variety testing. Euphytica 161:209–228.

Pickrell JK, Berisa T, Liu JZ, Ségurel L (2016) Detection and interpretation of shared genetic influences on 42 human traits. Nature Genetics 48:709–717. doi: 10.1038/ng.3570

Platt A, Vilhjalmsson BJ, Nordborg M (2010) Conditions under which genome-wide association studies will be positively misleading. Genetics 186:1045–1052.

Plomion C, Bastien C, Bogeat-Triboulot M-B, et al. (2016) Forest tree genomics: 10 achievements from the past 10 years and future prospects. Annals of Forest Science 73:77103. doi: 10.1007/s13595-015-0488-3

Postma FM, Ågren J (2016) Early life stages contribute strongly to local adaptation in Arabidopsis thaliana. Proceedings of the National Academy of Sciences 113:7590–7595. doi: 10.1073/pnas.1606303113

Price AL, Patterson NJ, Plenge RM, Weinblatt ME, Shadick NA, Reich D (2006) Principal components analysis corrects for stratification in genome-wide association studies. Nature Genetics 38:904–909.

Prout T, Barker JSF (1993) *F* statistics in *Drosophila buzzatii: selection, population size and inbreeding*. Genetics 134: 369–375.

Prunier J, Laroche J, Beaulieu J, Bousquet J (2011) Scanning the genome for gene SNPs related to climate adaptation and estimating selection at the molecular level in boreal black spruce. Molecular Ecology 20:1702–1716. doi: 10.1111/j.1365-294X.2011.05045.x

Prunier J, Verta J-P, MacKay JJ (2015) Conifer genomics and adaptation: at the crossroads of genetic diversity and genome function. New Phytologist 209:44–62.

Quesada, T., Li, Z., Dervinis, C., et al. (2008). Comparative analysis of the transcriptomes of *Populus trichocarpa* and *Arabidopsis thaliana* suggests extensive evolution of gene expression regulation in angiosperms. New Phytologist 180:408–420.

Ralph P, Coop G (2010) Parallel adaptation: one or many waves of advance of an advantageous allele? Genetics 186:647–668. doi: 10.1534/genetics.110.119594

Rellstab C, Gugerli F, Eckert AJ, et al. (2015) A practical guide to environmental association analysis in landscape genomics. Molecular Ecology 24:4348–4370. doi: 10.1111/mec.13322

Remington DL (2015) Alleles versus mutations: Understanding the evolution of genetic architecture requires a molecular perspective on allelic origins. Evolution 69:3025–3038. doi: 10.1111/evo.12775

Resende et al. (2012a) Accelerating the domestication of trees using genomic selection: accuracy of prediction models across ages and environments. New Phytologist 193: 617–624.

Resende et al. (2012b) Genomic selection for growth and wood quality in Eucalyptus: capturing the missing heritability and accelerating breeding for complex traits in forest trees. New Phytologist 194: 116–128.

Resende et al. (2012c) Accuracy of genomic selection methods in a standard data set of loblolly pine (*Pinus taeda* L.). Genetics 190: 1503–1510.

Riesch R, Muschick M, Lindtke D, et al. (2017) Transitions between phases of genomic differentiation during stick-insect speciation. Nat ecol evol 1:1–13. doi: 10.1038/s41559-017-0082

Ritland, K., Krutovsky, K.V., Tsumura, Y., Pelgas, B., Isabel, N. and Bousquet, J., 2011. Genetic mapping in conifers. In: Genetics, genomics and breeding of conifers, pp.196–238.

Ritland K, Ritland C (1996) Inferences about quantitative inheritance based on natural population structure in the yellow monkeyflower, Mimulus guttatus. Evolution 50:1074–1082.

Rockman, M. V. (2012, January). The QTN program and the alleles that matter for evolution: All that’s gold does not glitter. Evolution 66:1–17.

Rodíguez-Quilón I, Santos-del-Blanco L, Serra-Varela MJ, Koskela J, González-Martínez SC, Alía R (2016) Capturing neutral and adaptive genetic diversity for conservation in a highly structured tree species. Ecological Applications 26:2254–2266. doi:

Romero IG, Ruvinsky I, Gilad Y (2012) Comparative studies of gene expression and the evolution of gene regulation. Nature Publishing Group 13:505–516. doi: 10.1038/nrg3229

Roschanski AM, Csilléry K, Liepelt S et al. (2016) Evidence of divergent selection at landscape and local scales in *Abies alba* Mill. in the French Mediterranean Alps. Molecular Ecology 25:776–794.

Savolainen O, Lascoux M, Merilä J (2013) Ecological genomics of local adaptation. Nature Reviews Genetics 14:807–820. doi: 10.1038/nrg3522

Savolainen O, Pyhäjärvi T, Knürr T (2007) Gene Flow and Local Adaptation in Trees. Annual Review of Ecology, Evolution, and Systematics 38:595–619. doi: 10.1146/annurev.ecolsys.38.091206.095646

Schoville SD, Bonin A, Francois O, et al. (2012) Adaptive Genetic Variation on the Landscape: Methods and Cases. Annual Review of Ecology, Evolution, and Systematics 43:23–43. doi: 10.1146/annurev-ecolsys-110411-160248

Schrider DR, Kern AD (2016) S/HIC: Robust Identification of Soft and Hard Sweeps Using Machine Learning. PLoS Genetics 12:e1005928–31. doi: 10.1371/journal.pgen.1005928

Schrider DR, Mendes FK, Hahn MW, Kern AD (2015) Soft shoulders ahead: spurious signatures of soft and partial selective sweeps result from linked hard sweeps. Genetics 200:267–284. doi: 10.1534/genetics.115.174912/-/DC1

Schrider DR, Shanku AG, Kern AD (2016) Effects of linked selective sweeps on demographic inference and model selection. Genetics 204:1207–1223. doi: 10.1534/genetics.116.190223/-/DC1

Searle SR, Casella G, McCulloch CE (1992) Variance Components. New York, NY: John Wiley & Sons, Inc. 528 pp.

Silva-Junior O, Faria DA, Grattapaglia D (2015) A flexible multi-species genome-wide 60K SNP chip developed from pooled resequencing of 240 Eucalyptus tree genomes across 12 species. New Phytologist 206:1527–1540. doi: 10.1111/nph.13322

Simons YB, Bullaughey K, Hudson RR, Sella G (2017) A model for the genetic architecture of quantitative traits under stabilizing selection. bioRxiv 1–76. doi: arXiv:1704.06707

Siol M, Wright S, Barrett S (2010) The population genomics of plant adaptation. New Phytologist 188: 313–332.

Slate J (2005) Quantitative trait locus mapping in natural populations: progress, caveats and future directions. Molecular Ecology 14:363–379. doi: 10.1111/j.1365-294X.2004.02378.x

Slatkin M (1987) Gene flow and the geographic structure of natural populations. Science 236:787–793. doi: 10.1126/science.3576198

Slatkin M (1975) Gene flow and selection in a two-locus system. Genetics 81:787–802.

Smith SD (2016) Pleiotropy and the evolution of floral integration. New Phytologist 209:80–85. doi: 10.1111/nph.13583

Sork VL, Aitken SN, Dyer RJ, et al. (2013) Putting the landscape into the genomics of trees: approaches for understanding local adaptation and population responses to changing climate. Tree Genetics & Genomes 9:901–911. doi: 10.1007/s11295-013-0596-x

Speed, D., & Balding, D. J. (2014). MultiBLUP: improved SNP-based prediction for complex traits. Genome Research, 24(9), 1550–1557. http://doi.org/10.1101/gr.169375.113

Spitze K (1993) Population structure in *Daphnia obtusa:* quantitative genetic and allozyme variation. Genetics 135:367–374.

Spencer CCA, Su Z, Donnelly P, Marchini J (2009) Designing genome-wide association studies: sample size, power, imputation, and the choice of genotyping chip. PLoS Genetics 5:e1000477.

St Clair JB, Mandel NL, Vance-Borland KW (2005) Genecology of Douglas fir in western Oregon and Washington. Annals of Botany 96:1199–1214.

Stephan, W. (2010) Genetic hitchhiking versus background selection: the controversy and its implications. Philos Trans R Soc Lond B Biol Sci 365: 1245–1253.

Stephan W (2015) Signatures of positive selection: from selective sweeps at individual loci to subtle allele frequency changes in polygenic adaptation. Molecular Ecology 25:79–88. doi: 10.1111/mec.13288

Stephens M, Balding DJ (2009) Bayesian statistical methods for genetic association studies. Nature Reviews Genetics 10:681–690.

Stinchcombe JR, Hoekstra HE (2008) Combining population genomics and quantitative genetics: finding the genes underlying ecologically important traits. Heredity 100:158–170. doi: 10.1038/sj.hdy.6800937

Stölting KN, Paris M, Meier C, Heinze B, Castiglione S, Bartha D, Lexer C (2015) Genome-wide patters of differentiation and spatially varying selection between postglacial recolonization lineage of *Populus alba* (Salicaceae), a widespread forest tree. New Phytologist 207:723–734. doi: 10.1111/nph.13392

Storey JD, Tibshirani R (2003) Statistical significance for genomewide studies. Proceedings of the National Academy of Sciences of the United States of America 100:9440–9445.

Strickler SR, Bombarely A, Mueller LA (2012) Designing a transcriptome next-generation sequencing project for a nonmodel plant species1. American Journal of Botany 99:257–266. doi: 10.3732/ajb.1100292

Suren H, Hodgins KA, Yeaman S, Nurkowski KA, Smets P, Rieseberg LH, Aitken SN, Holliday JA (2016) Exome capture from the spruce and pine giga-genomes. Molecular Ecology 16:1136–1146.

Tan, B., Grattapaglia, D., Wu, H. X., & Ingvarsson, P. K. (2017). Genomic prediction reveals significant non-additive effects for growth in hybrid Eucalyptus. bioRxiv, 1–35. http://doi.org/10.1101/178160

Temesgen B, Brown GR, Harry DE, Kinlaw CS, Sewell MM, Neale DB (2001) Genetic mapping of expressed sequence tag polymorphism (ESTP) markers in loblolly pine (Pinus taeda L.). Theor Appl Genet 102:664–675

Tenaillon O (2014) The Utility of Fisher’s Geometric Model in Evolutionary Genetics. Annual Review of Ecology, Evolution, and Systematics 45:179–201. doi: 10.1146/annurev-ecolsys-120213-091846

Thavamanikumar S, Southerton SG, Bossinger G, Thumma BR (2013) Dissection of complex traits in forest trees — opportunities for marker-assisted selection. Tree Genetics & Genomes 9:627–639. doi: 10.1007/s11295-013-0594-z

Tiffin P, Ross-Ibarra J (2014) Advances and limits of using population genetics to understand local adaptation. Trends in Ecology & Evolution 29:673–680. doi: 10.1016/j.tree.2014.10.004

Tigano A, Friesen VL (2016) Genomics of local adaptation with gene flow. Molecular Ecology 25:2144–2164. doi: 10.1111/mec.13606

Timpson NJ, Greenwood CMT, Soranzo N, et al. (2017) Genetic architecture: the shape of the genetic contribution to human traits and disease. Nature Publishing Group 1–15. doi: 10.1038/nrg.2017.101

Turelli M, Barton NH (1994) Genetic and statistical analyses of strong selection on polygenic traits: what, me normal? Genetics 138:913–941.

Vasquez-Gross, H. A., Yu, J. J., Figueroa, B., Gessler, D. D., Neale, D. B., & Wegrzyn, J. L. (2013). CartograTree: connecting tree genomes, phenotypes and environment. Molecular Ecology Resources 13: 528–537.

Via S, Lande R (1985) Genotype-environment interaction and the evolution of phenotypic plasticity. Evolution 39:505–522. doi: 10.1111/j.1558-5646.1985.tb00391.x

Vialette-Guiraud ACM, Andres-Robin A, Chambrier P, et al. (2016) The analysis of Gene Regulatory Networks in plant evo-devo. Journal of Experimental Botany 67:2549–2563. doi: 10.1093/jxb/erw119

Vitezica ZG, Legarra A, Toro MA, Varona L (2017) Orthogonal Estimates of Variances for Additive, Dominance, and Epistatic Effects in Populations. Genetics 206:1297–1307. doi: 10.1534/genetics.116.199406

Vizcaíno-Palomar N, Revuelta-Eugercios B, Zavala MA, Alia R, González-Martínez SC (2014) The role of population origin and microenvironment in seedling emergence and early survival in Mediterranean maritime pine (*Pinus pinaster* Aiton). PLoS ONE 9:e109132.

Wachowiak W, Trivedi U, Perry A, Cavers S (2015) Comparative transcriptomics of a complex of four European pine species. BMC Genomics 16:1–9. doi: 10.1186/s12864-015-1401-z

Wadgymar SM, Lowry DB, Gould BA, et al. (2017) Identifying targets and agents of selection: innovative methods to evaluate the processes that contribute to local adaptation. Methods in Ecology and Evolution 8:738–749. doi: 10.1111/2041-210X.12777

Wagner GP, Altenberg L (1996) Perspective: complex adaptations and the evolution of evolvability. Evolution 50:967. doi: 10.2307/2410639

Wagner GP, Kenney-Hunt JP, Pavlicev M, et al. (2008) Pleiotropic scaling of gene effects and the “cost of complexity.” Nature 452:470–472. doi: 10.1038/nature06756

Wagner GP, Pavlicev M, Cheverud JM (2007) The road to modularity. Nature Reviews Genetics 8:921–931. doi: 10.1038/nrg2267

Wagner GP, Zhang J (2011) The pleiotropic structure of the genotype–phenotype map: the evolvability of complex organisms. Nature Publishing Group 12:204–213. doi: 10.1038/nrg2949

Wang L, Beissinger TM, Lorant A, Ross-Ibarra C, Ross-Ibarra J, Hufford M (2017) The interplay of demography and selection during maize domestication and expansion. bioRxiv doi: https://doi.org/10.1101/114579

Wang Z, Liao BY, Zhang J (2010) Genomic patterns of pleiotropy and the evolution of complexity. Proceedings of the National Academy of Science 107:18034–18039. doi: 10.1073/pnas.1004666107

Wegrzyn JL, Lee JM, Tearse BR, Neale DB (2008) TreeGenes: A Forest Tree Genome Database. International Journal of Plant Genomics 2008:1–7. doi: 10.1155/2008/412875

Wegrzyn JL, Main D, Figueroa B, et al. (2012) Uniform standards for genome databases in forest and fruit trees. Tree Genetics & Genomes 8:549–557. doi: 10.1007/s11295-012-0494-7

Welch JJ, Waxman D (2003) Modularity and the cost of complexity. Evolution 57:1723–13. doi: 10.1554/02-673

Wellenreuther M, Hansson B (2016) Detecting Polygenic Evolution: Problems, Pitfalls, andPromises. Trends in Genetics 32:155–164. doi: 10.1016/j.tig.2015.12.004

Whitlock MC (1999) Neutral additive genetic variance in a metapopulation. Genetical Research 74:215–221.

Whitlock MC (2003) Fixation probability and time in subdivided populations. Genetics 164:767–779.

Whitlock MC, Gilbert KJ (2012) Q_ST_ in a hierarchically structured population. Molecular Ecology Resources 12:481–483.

Whitlock, M. C. and F. Guillaume. 2009. Testing for spatially divergent selection: Comparing Q_S_τ to F_ST_. Genetics 183: 1055–1063.

Whitlock MC, Lotterhos KE (2015) Reliable detection of loci responsible for local adaptation: Inference of a null model through trimming the distribution of F_ST_. The American Naturalist 186:S24–S36. doi: 10.1086/682949

Whitlock MC, Phillips PC, Moore FB (1995) Multiple fitness peaks and epistasis. Annual Review of Ecology and Systematics 26:601–629. doi: annurev.es.26.110195.003125

Wong CK, Bernardo R (2008) Genomewide selection in oil palm: increasing selection gain per unit time and cost with small populations. Theor Appl Genet 116: 815–824.

Wortley, A. H., & Scotland, R. W. (2004). Synonymy, sampling and seed plant numbers. Taxon, 53(2), 478–480.

Wright S (1932) The roles of mutation, inbreeding, crossbreeding, and selection in evolution. Proceedings of the Sixth International Congress on Genetics, Vol. 1, No. 6. (1932), pp. 356–366.

Wright S (1931) Evolution in Mendelian populations. Genetics 16:97–159.

Xu S (2003) Theoretical Basis of the Beavis Effect. Genetics 165:2259–2268. doi: 10.1038/hdy.1992.131

Yang J, Benyamin B, McEvoy BP, et al. (2010) Common SNPs explain a large proportion of the heritability for human height. Nature Genetics 42:565–569. doi: 10.1038/ng.608

Yang, J., S. Mezmouk, A. Baumgarten, E. S. Buckler, K. E. Guill, M. D. McMullen, and J. Ross-Ibarra. 2017. Incomplete dominance of deleterious alleles contribute substantially to trait variation and heterosis in maize. bioRxiv. Doi: https://doi.org/10.1101/086132

Yeaman S (2013) Genomic rearrangements and the evolution of clusters of locally adaptive loci. Proceedings of the National Academy of Sciences 110:E1743–51. doi: 10.1073/pnas.1219381110

Yeaman S (2015) Local Adaptation by Alleles of Small Effect. The American Naturalist 186:S74–S89. doi: 10.1086/682405

Yeaman S, Hodgins KA, Lotterhos KE, et al. (2016) Convergent local adaptation to climate in distantly related conifers. Science 353:1431–1433. doi: 10.1126/science.aaf7812

Yeaman S, Jarvis A (2006) Regional heterogeneity and gene flow maintain variance in a quantitative trait within populations of lodgepole pine. Proceedings of the Royal Society B: Biological Sciences 273:1587–1593. doi: 10.1534/genetics.166.2.1053

Yeaman S, Otto SP (2011) Establishment and maintenance of adaptive genetic divergence under migration, selection, and drift. Evolution 65:2123–2129. doi: 10.1111/j.1558-5646.2011.01277.x

Yeaman S, Whitlock MC (2011) The genetic architecture of adaptation under migration-selection balance. Evolution 65:1897–1911. doi: 10.1111/j.1558-5646.2011.01269.x

Yoder JB, Tiffin P (2017) Effects of Gene Action, Marker Density, and Timing of Selection on the Performance of Landscape Genomic Scans of Local Adaptation. Journal of Heredity 1–13. doi: 10.1093/jhered/esx042

Yu J, Pressoir G, Briggs WH, Vroh Bi I, Yamasaki M et al. (2005) A unified mixed-model method for association mapping that accounts for multiple levels of relatedness. Nature Genetics 38:203–208.

Zhang X-S (2012) Fisher’s geometric model of fitness landscape and variance in fitness within a changing environment. Evolution 66:2350–2368. doi: 10.1111/j.1558-5646.2012.01610.x

Zhang M, Zhou L, Bawa R, Suren H, Holliday JA (2016) Recombination rate variation, hitchhiking, and demographic history shape deleterious load in poplar. Molecular Biology and Evolution. 33:2899–2910. doi: https://doi.org/10.1093/molbev/msw169

Zhong S, Dekkers JC, Fernando RL, Jannink JL (2009) Factors affecting accuracy from genomic selection in populations derived from multiple inbred lines: a barley case study. Genetics 182:355–364.

Zhou, X., Carbonetto, P., & Stephens, M. (2013). Polygenic modeling with Bayesian sparse linear mixed models. PLoS Genetics, 9(2), e1003264. http://doi.org/10.1371/journal.pgen.1003264

Zinkgraf M, Liu L, Groover A, Filkov V (2017) Identifying gene coexpression networks underlying the dynamic regulation of wood-forming tissues in Populus under diverse environmental conditions. New Phytologist 214:1464–1478. doi: 10.1111/nph.14492

Zöllner S, Pritchard JK (2007) Overcoming the Winner’s Curse: Estimating Penetrance Parameters from Case-Control Data. The American Journal of Human Genetics 80:605–615. doi: 10.1086/512821

Zuk O, Hechter E, Sunyaev SR (2012) The mystery of missing heritability: Genetic interactions create phantom heritability. Proceedings of the National Academy of Science 109:1193–1198. doi: 10.1073/pnas.1119675109/-/DCSupplemental

## Supplemental References

Devlin B, Roeder K (1999) Genomic control for association studies. Biometrics 55:997–1004.

Johnson RC, Nelson GW, Troyer JL, et al (2010) Accounting for multiple comparisons in a genome-wide association study (GWAS). BMC Genomics 11:724.

Pritchard JK, Pickrell JK, Coop G (2010) The Genetics of Human Adaptation: Hard Review Sweeps, Soft Sweeps, and Polygenic Adaptation. Current Biology 20:R208–R215. doi: 10.1016/j.cub.2009.11.055

Wegrzyn JL, Eckert AJ, Choi M, et al (2010) Association genetics of traits controlling lignin and cellulose biosynthesis in black cottonwood (*Populus trichocarpa*, Salicaceae) secondary xylem. New Phytologist 188:515–532. doi: 10.1111/j.1469-8137.2010.03415.x

Zöllner S, Pritchard JK (2007) Overcoming the Winner’s Curse: Estimating Penetrance Parameters from Case-Control Data. The American Journal of Human Genetics 80:605–615.

